# Cellular protein delivery through membrane potential driven water pores

**DOI:** 10.64898/2026.02.24.707441

**Authors:** Jonathan Franke, Palina Dubatouka, Amin Yourdkhani, Sahil Soni, Tillmann Utesch, Juliana Serrano, Tolga Soykan, Martin Lehmann, Han Sun, Jan Vincent V. Arafiles, Christian P. R. Hackenberger

## Abstract

Providing immediate access for functional proteins inside living cells would unlock unprecedented control over cellular processes; however, commonly used endocytic delivery suffers from endosomal trapping and degradation. One of the most powerful non-endosomal delivery methods uses cell surface anchored cell penetrating peptide (CPP)-additives that allow proteins to enter cells directly. Nevertheless, the underlying molecular mechanism involved in direct entry via crossing the cell membrane (protein translocation through the cell) and the major driving forces remain controversially discussed. Here, we provide a stepwise molecular picture on how CPP-additives enable uptake of protein cargoes through direct membrane translocation. CPP-additives accumulate on the cell surface in nucleation zones, locally hyperpolarizing the membrane, and induce transient water pores that allow selective CPP-protein entry without compromising membrane integrity.

These fundamental mechanistic insights provide a firm basis for rationally optimizing delivery strategies using highly cationic CPPs, ultimately resulting in innovative and smart protein delivery strategies to advance therapeutic protein applications.

**Abstract Figure:** 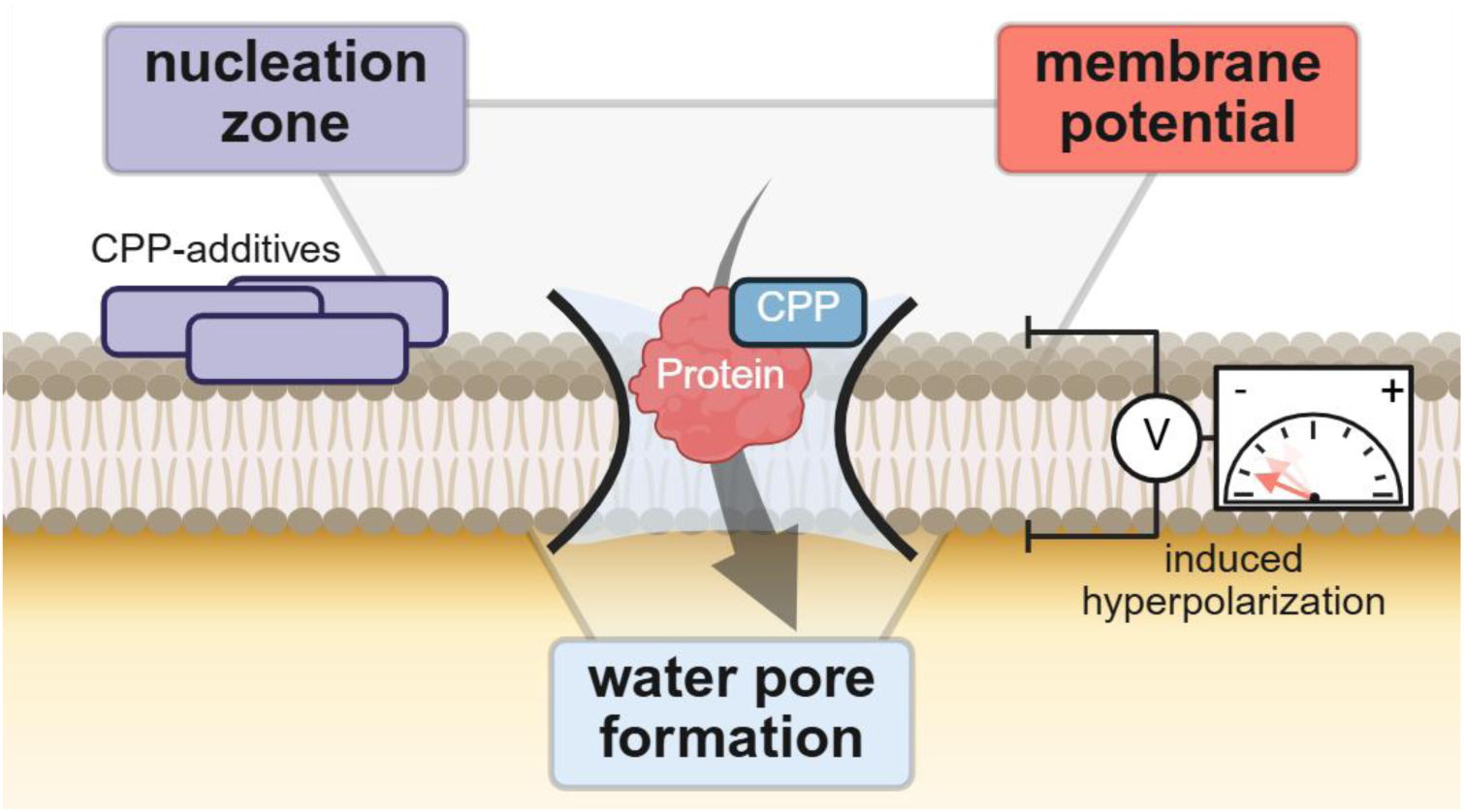

## Introduction

Cellular delivery is a key challenge in the molecular life sciences, given the growing demand for peptide- and protein-based biopharmaceuticals acting on intracellular targets. Unlocking the full therapeutic potential of these biomacromolecules requires robust, versatile, and efficient technologies capable of mediating their safe and selective transport across the plasma membrane into the cytosol.^[1–3]^

Biomacromolecules can enter cells through either endocytic pathways or direct translocation across the plasma membrane, whereas conventional delivery strategies — such as liposomes, polymeric nanoparticles, and other vesicular carriers — primarily rely on endocytic uptake.^[4]^ This is accompanied by endosomal entrapment, lysosomal degradation, and recycling which altogether severely restrict the cytosolic bioavailability of these payloads.^[5–7]^ In contrast, direct translocation provides a more promising route, allowing biomacromolecules to bypass the endosomal system and access their site of action in the cytoplasm.^[8, 9]^ Current physical approaches to achieve direct membrane permeabilization (e.g., electroporation^[10, 11]^, mechanoporation^[12]^, microfluidic squeezing^[13]^) provide membrane crossing but are largely confined to *in vitro* or *ex vivo* use and often compromise cell viability.^[1]^

Cell-penetrating peptides (CPPs) have emerged as a powerful alternative for facilitating the cellular entry of functional macromolecules.^[14–16]^ Conjugation of CPPs to proteins can promote direct penetration of the cellular membrane; however, cyclic CPPs as well as higher concentrations of the CPP-conjugate are needed to achieve efficient non-endosomal delivery into the cytosol.^[16–18]^ We recently developed a modular approach for achieving a more efficient cytoplasmic delivery of native proteins in living cells using tailored Arg-rich CPP-additives at low concentrations (Fig. 1a).^[19, 20]^ In this system, linear CPP-additives transiently associate with the plasma membrane to form localized nucleation zones, which prime the membrane for the direct translocation of CPP–protein conjugates. Such CPP-additive strategy enables cytosolic delivery within minutes while bypassing endosomal uptake and thereby establishing a promising route for functional protein delivery, as exemplified in the delivery of a nanobody reconstituting cystic fibrosis transmembrane conductance regulator (CFTR).^[22]^ In addition, a fully reversible protein bioconjugation strategy on amines (BioRAM^[21]^) can be used for CPP-protein conjugation, achieving full recovery of the native protein inside cells, as recently demonstrated for an active opioid receptor targeting nanobody.^[23]^

**Fig. 1.**
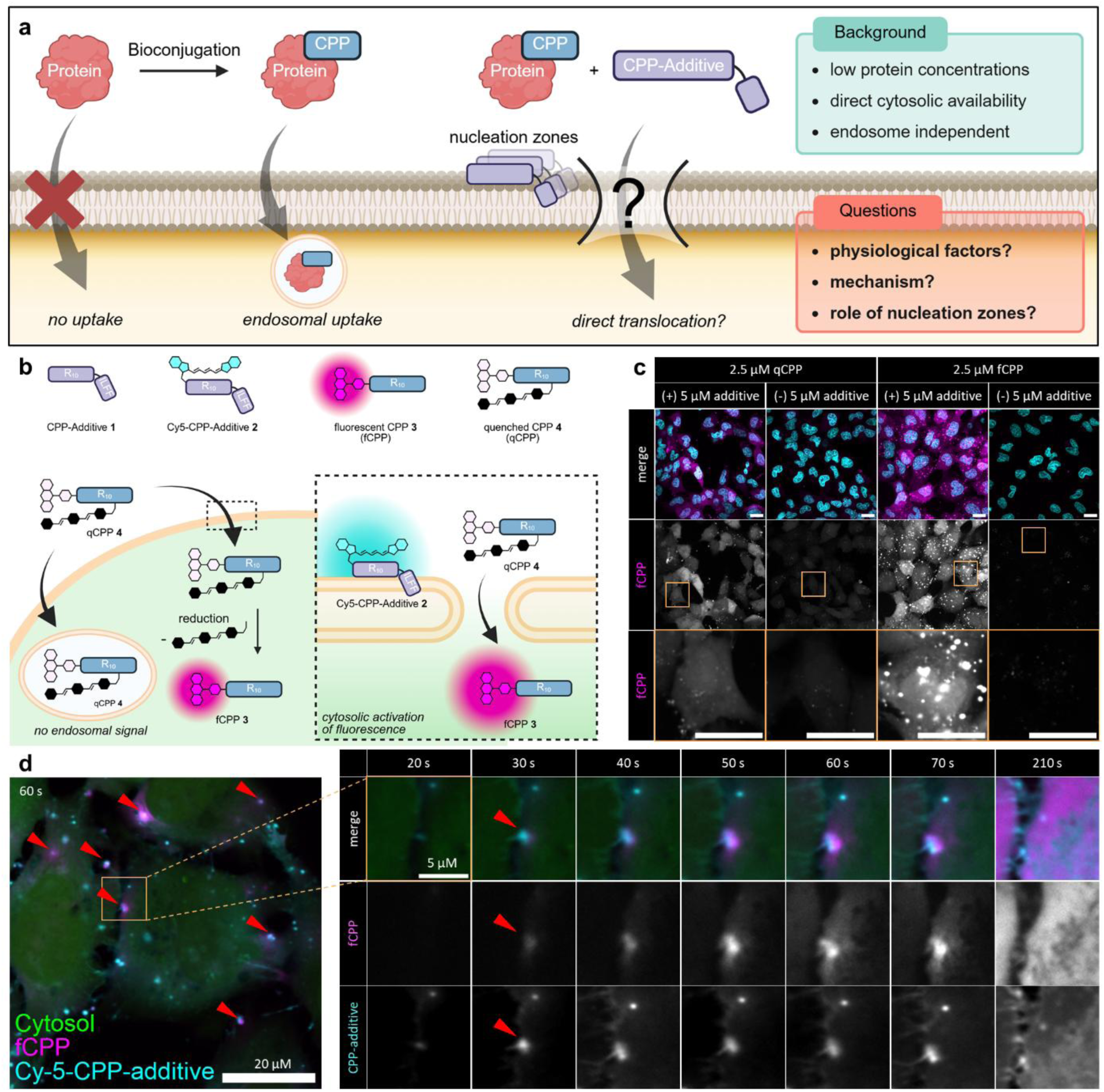
CPP-additive mediated uptake and quenched probe design to study direct translocation. a) Current understanding of CPP-additive mediated protein delivery schematically illustrated. b) Schematic representation of peptide structures **1**−**4** and CPP-additive mediated entry of qCPP **4** and intracellular activation of fluorescence through reduction to fCPP **3**. c) Incubation of HeLa cells with 2.5 µM fCPP **3** or qCPP **4** in the presence and absence of 5 µM CPP-additive **1** for 30 min in serum-free FluoroBrite Dulbecco’s Modified Eagle Medium (FB DMEM). Nuclei were co-stained with Hoechst (cyan). Scale bar = 20 µm. **d**) Time-lapse microscopy of 2.5 µM qCPP **4** uptake mediated by 2.5 µM Cy5-CPP-additive **2** (green: cytosolic marker 5-chloromethylfluorescein diacetate (CMFDA), magenta: fCPP **4** (λ_abs, max_ = 552 nm, λ_em, max_ = 578 nm), cyan – Cy5-CPP-additive **2** (λ_abs, max_ = 651 nm, λ_em, max_ = 670 nm)). Hela cells treated with qCPP **4** and CPP-additive **2,** 2.5 µM each in serum-free FB DMEM. Red arrows indicate nucleation zones, which serve as entry points for qCPP **4**. Highlighted region of interest is displayed at different time points, visualizing the development of activated fCPP after uptake.

Previous mechanistic work has identified several key parameters influencing the direct translocation of small arginine-rich polycationic CPPs. These factors include membrane lipid composition and architecture^[24–26]^, peptide concentration^[18, 27]^, presence of counterions^[28]^, glycocalyx composition^[29, 30]^, and the membrane potential^[31–33]^. Among these, membrane potential has been proposed as a decisive driving force: interaction of polycationic CPPs with negatively charged membrane surface can induce transient depolarization events and the formation of nanoscopic water pores, facilitating peptide entry.^[34, 35]^ While such models have advanced our understanding of how short polyarginine CPPs can traverse cell membranes, the direct translocation mechanisms for larger CPP-protein-conjugates in a macroscopic cellular environment remains elusive.^[36, 37]^

In the present study, we integrate biochemical, biophysical and computational evidence to elucidate the molecular mechanism underlying direct translocation of CPP–protein facilitated by CPP-additives. We focus on the interplay between nucleation zone, membrane potential modulation, and transient water pore formation to delineate how these parameters collectively govern spontaneous cytosolic entry of peptide and protein cargoes. This mechanistic framework provides the first unified account of for combining molecular driving forces in CPP-based delivery strategies, offering guidelines toward more efficient and non-destructive intracellular delivery of therapeutic and research-relevant proteins.

## Results

### CPP-additive mediated uptake through nucleation zones

Based on previous findings from our group^[19, 20]^ and others^[9, 38–40]^, we hypothesize that CPP-additives play a pivotal role in nucleation zones formation, thereby facilitating the uptake of additional CPP substrates. Although nucleation zones have been repeatedly described in literature, their specific contribution to cellular uptake remains poorly understood. Confocal fluorescence microscopy is a commonly used tool for investigating cellular such processes, as it enables temporally resolved localization of fluorescently labeled molecules. Nevertheless, to delineate the impact of CPP-additive mediated nucleation zones on the direct cytosolic availability of peptide and protein cargoes, we first had to devise specifically tailored fluorescent probes.

Our previously reported CPP-additive **1** is a decaarginine sequence with a *N*-terminal thiol reactive disulfide and *C*-terminal hydrophobic amino acid sequence (ILFF), designed to anchor the peptide additive on the cell surface (Fig. 1b).^[19]^ In order to simultaneously monitor localization of the CPP-additive and a CPP-conjugated model-cargo within a short timeframe, we obtained CPP-additive **2** equipped with Cyanine5 (Cy5), and a fluorescent CPP-cargo **3** (fCPP, 5,6-carboxytetramethylrhodamine (TAMRA-labeled R_10_) for live-cell confocal time-lapse microscopy (Fig. 1b).^[20]^ Upon addition of CPP-additive **2** and fCPP **3** to HeLa cells, CPP-additive **2** immediately accumulates on the cell surface. Besides general cell surface association, we observed areas of CPP-additive accumulation appearing as puncta with intense Cy5 signal, which we consider as nucleation zones (Supplementary Fig. 1).^[8, 9]^ In parallel, fCPP **3** quickly diffuses inside the cells; however, the high fluorescence background prevents clear distinction between intra- and extracellular signals, hence limiting the ability to correlate these nucleation zones with entry sites of TAMRA-R_10_ **3**. To mitigate this limitation, we designed a quenched CPP-cargo (qCPP) **4** that displays a turn-on fluorescence upon encountering the reductive intracellular environment. qCPP **4** was synthesized by forming a disulfide between fCPP **3** and a diazo-containing Black-Hole-Quencher (BHQ-2) (Extended Data Fig. 1a). Quenching by BHQ-2 occurs via Förster resonance energy transfer due to the overlap of the absorption spectrum of the BHQ-2 with the emission of the TAMRA fluorophore upon excitation.^[41]^ Reduction of the disulfide bond under physiological conditions liberates the BHQ-2 from the construct, resulting in the generation of fluorescent fCPP **3** (Extended Data Fig. 1b). Although BHQ-2 increases hydrophobicity of the qCPP **4**, membrane disruption and acute toxicity was not observed for the qCPP **4** at treatment concentrations (2.5 µM) used in this study (Supplementary Fig. 2). While permeability of the whole construct would likely be affected, we do not expect a change in its uptake mechanism.^[42]^

The cellular uptake of the qCPP **4** (2.5 µM) was initially tested in HeLa cells by incubation for 30 min in presence and absence of the CPP-additive **1** (2.5 µM, Fig. 1c). Gratifyingly, we observed diffuse fluorescent TAMRA-signals in the cytosol, indicative of cytosolic activation and successful cargo transport. Moreover, cells were devoid of endosomal signals as endocytosed qCPP **4** remains quenched because of the oxidizing lumen inside the endosome.^[43]^ Still, after incubation periods longer than 5 min, few punctate signals appear, which is most likely a consequence of endocytosis of fCPP **3** formed by extracellular reduction of qCPP **4**.^[44]^ In the absence of CPP-additive **1**, qCPP **4** (2.5 µM) shows markedly reduced cytosolic signal, yet uptake is not completely abolished compared to fCPP **3** treated cells, probably due to the attachment of the hydrophobic BHQ-2 altering permeation properties as explained above. On the other hand, direct treatment with unquenched fCPP **3** (2.5 µM) and CPP-additive **1** (2.5 µM) showed both punctate endosomal as well as diffused cytosolic signals (Fig. 1c).

Next, we applied the qCPP **4** on HeLa cells in combination with the fluorescently labeled Cy5-CPP-additive **2** and monitored the respective fluorescent signals using time-lapse microscopy. The Cy5-CPP-additive **2** accumulates on the cell surface and forms nucleation zones within 20 seconds after addition, in accordance with previous results (red arrow, Fig. 1d).^[19, 20]^ This event is immediately followed by the emergence of fCPP **3** signal progressively developing from several distinct intracellular sites as a result of qCPP **4** uptake and cytosolic reduction to fCPP **3** (Supplementary Video 1). Using an automated Fiji macro, we quantified the colocalization of Cy5-CPP-additive **2** and fCPP **3** signals over time (see Supplementary Information for detailed procedure). From 30 to 90 seconds, the ratio of fCPP signal to CPP-additive signal increased, suggesting a colocalization of the signals within the region of interest (Extended Data Fig. 2a). This is followed by the fCPP **3** signal quickly spreading throughout the whole cell, resulting in a decreased overlap of fCPP **3** and Cy5-CPP-additive **2** signals, indicated by a decrease of the quotient. This result parallels the qualitative observation in which the fCPP **3** signal evolves from the fluorescent puncta of Cy5-CPP-additive **2** on the cell surface (Please refer to Supplementary Fig. 3 for discussion of out-of-plane localization).

Using the extracellularly quenched probe qCPP **4** allowed unambiguous discrimination of extracellular and intracellular cytosolic signals. We demonstrate that qCPP **4** primarily enters through the extracellular nucleation zones of Cy5-CPP-additives **2,** followed by fast cytosolic reduction to fluorescent fCPP **3**. We conclude that nucleation zones formed by CPP-additive **2**, serve as direct entry points for qCPP **4** into cells.

### Membrane potential as driving force for CPP-additive mediated uptake

Previous studies (e.g. Rothbard et al^[33]^, Hallaj et al^[34]^, Gao et al^[35]^, and Trofimenko et al.^[45]^) have suggested that the cell membrane potential (V_m_) plays a key role in the translocation of highly positively charged CPPs; however, these studies have not probed dynamic changes in V_m_ upon CPP exposure, especially in the context of externally added CPPs.

Therefore, to monitor V_m_ over time in the presence of CPP-additives and cargoes, we performed cell microscopy experiments taking advantage of the fluorescent dye DiBAC_4_(3) (bis-(1,3-dibutylbarbituric acid) trimethine oxonol),^[46]^ which allows a continuous V_m_ read-out based on its distribution across membrane (Extended Data Fig. 3b-c).^[47–49]^ We equilibrated HeLa cells with DiBAC_4_(3) after which, 2.5 µM qCPP **4** and 5 µM CPP-additive **1** were added together. The change in DiBAC_4_(3) fluorescence and the intracellular fluorescence of fCPP **3** (after cytosolic reduction of qCPP **4)** were recorded simultaneously at 37 °C (Fig. 2a). The fluorescence signal was normalized to the intensity of frame 0 before addition of the CPPs. Compared to the medium treated cells, we observed a significant decrease in DiBAC_4_(3) signal for cells co-treated with qCPP **4** and CPP-additives **1** (Fig. 2b-c). The DiBAC_4_(3) signal reaches a minimum between 80−120 seconds before recovering to the baseline level, suggesting induced hyperpolarization upon addition of qCPP **4** and CPP-additive **1**. Based on literature, the shift compared to the baseline of 20% corresponds to a change in membrane potential of ∼ −20 mV from the resting potential.^[49, 50]^ After experimental evidence that addition of CPPs temporarily affects V_m_, we tested the effect of abrogating the membrane potential on CPP-additive-mediated uptake. Gramicidin is commonly used to permanently depolarize cells by acting as an ionophore through the formation of channels in the membrane lipid bilayer.^[51]^ Upon pretreatment of HeLa cells with DiBAC_4_(3) dye, incubation of gramicidin (1.2 µg/mL) for 5 min and subsequent addition of qCPP **4** and Cy5-CPP-additive **2**, we observed accumulation of Cy5-CPP-additive **2** on the cell surface; however, we found no uptake of qCPP **4** (Supplementary Fig. 4). This finding suggests that the permanent depolarization blocks qCPP **4** uptake into HeLa cells.

**Fig 2.**
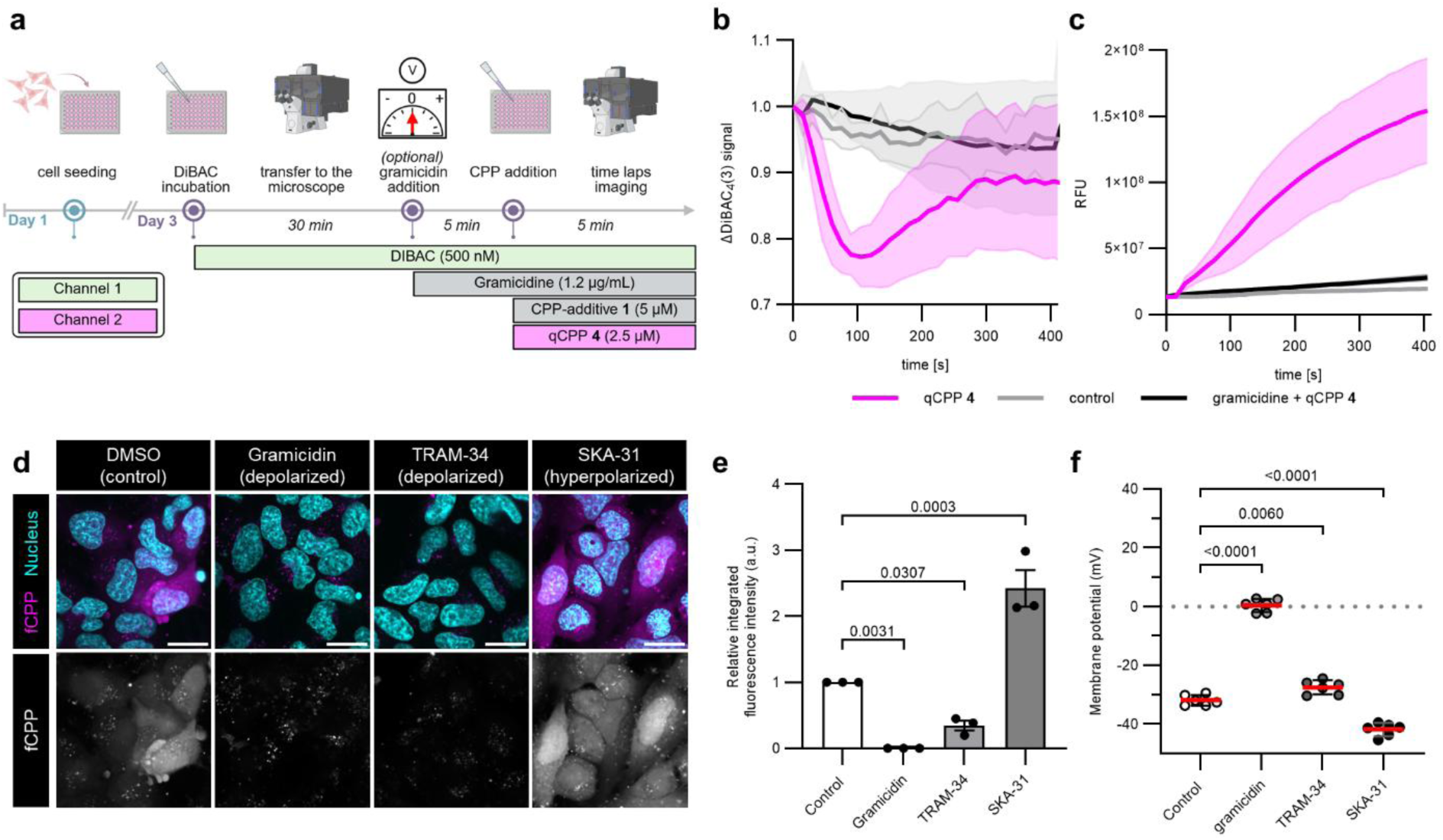
The role of membrane potential (V_m_) for CPP-additive mediated uptake. a) Workflow for time-lapse microscopy to simultaneously monitor V_m_ (DiBAC_4_(3) fluorescence (Channel 1), and qCPP **4** uptake (Channel 2). b) Change in membrane potential measurement acquired through quantification of intracellular signal of channel 1 (DiBAC_4_(3)); control = FB-DMEM medium. Data represent mean intracellular fluorescence of three regions of interest (ROIs) normalized to t_0_=1 ± standard error (SE) (N=3). c) Intracellular signal of channel 2 (qCPP **4**→fCPP **3**) measured over time. Data represent mean intracellular fluorescence of three ROIs normalized to t_0_=0 ± SE (N=3). d) Confocal images of HeLa cells pre-treated with 1 µg/mL gramicidin, 20 µM TRAM-24, or SKA-31, and incubated with 2.5 µM qCPP **4** and 5 µM CPP-additive **1** for 30 min. Scale bars = 20 µm. e) Median nuclear fluorescence intensities of fCPP **3** after treatment with 2.5 µM qCPP **4** in 600 cells across three biological replicates normalized against the control. Data were normalized (non-treated control=1, gramicidin treated=0), after qualitative validation of no diffused signal in gramicidin treated cells. Data presented as mean ± SE. *P*-values were determined by one-way analysis (ANOVA) followed by Dunnett’s post hoc test. f) Intracellular DiBAC_4_(3) fluorescence-based membrane potential measurement in HeLa cells treated with 1 µg/mL gramicidin, 20 µM TRAM34, or 20 µM SKA-31. Red line represents the mean ± SE of six individual data points taken across three biological replicates. *P*-values were determined by one-way analysis (ANOVA) followed by Dunnett’s post hoc test.

We then questioned whether further modulation of the resting V_m_ could affect the CPP-additive mediated direct translocation efficiency of qCPP **4**. To test this, we pre-treated HeLa cells with TRAM-34 (calcium-activated potassium channel inhibitor)^[52]^ and SKA-31 (calcium-activated potassium channel activator)^[53]^ and quantified fCPP **3** signal in the presence of CPP-additive **1**(Fig. 2d-f). Similar to previous reports, we observed that increasing the resting V_m_ through SKA-31 treatment (from −32 mV to −40 mV) resulted in increased uptake (almost 2-fold), whereas decreasing the resting V_m_ through TRAM-34 treatment (from −32 mV to −26 mV) decreased uptake by a factor of 2 (Fig. 2e-f).^[31, 54]^ It is clear that CPP-additive–mediated delivery of qCPP **4** is tightly coupled to membrane polarization in the resting state: lowering polarization consistently suppresses direct translocation, whereas increasing polarization promotes it.^[31, 54]^

Our data demonstrates that CPP-additive treatment causes a temporary hyperpolarization event associated with CPP uptake. This uptake is found to be dependent on resting membrane potential (V_m_) with more polarized cells exhibiting increased uptake.

### Temporary hyperpolarization induced water pore formation

The observation of peptides entering through nucleation zones and temporary membrane hyperpolarization raises the question of their mechanistic relationship. Previous studies have proposed water pore formation as principal entry mechanism for cationic CPPs. These simulations, however, relied on coarse-grained models^[31]^ or were performed over relatively short timescale^[35]^, and importantly did not maintain a continuous membrane voltages throughout the entire CPP translocation process. To extend these efforts, we performed sub-microsecond atomistic MD simulations of CPP permeation processes in POPC lipid bilayers using the Computational Electrophysiology (CompEL) setup (Extended Fig. 3a). In this framework, a defined ion imbalance across the membrane that naturally generates a membrane potential is maintained.^[45]^ We propose that this configuration mimics the local environment of nucleation zones, where the accumulation of CPP-additives induces a strong local transmembrane potential which we observed experimentally as a global hyperpolarization event in Fig. 2b.

We simulated here cationic linear R_10_ and compared it to G_10_, a non-penetrating control. Pre-screening simulations of 100 ns across different ionic imbalances, showed that an ionic imbalance of 18e, corresponding to a membrane voltage of ∼ 2 V, consistently induced pore formation in both models (Supplementary Fig. 5). Despite the high voltage, the overall system remained energetically stable. Similar high voltages have been also applied in previous simulation studies^[31, 35]^, where they were necessary to accelerate pore formation and CPP translocation within sub-microsecond timescales. We therefore selected a transmembrane voltage of ∼2 V for subsequent in-depth production simulations. It should be noted that while the precise local transmembrane potentials within nucleation zones remain unknown, they are likely substantially higher than the average global membrane potential measured in cells.

Across seven independent 500 ns MD production runs for each peptide, pore formation was consistently observed (Fig. 3a-b). This indicates that the pore formation is primarily driven by the applied membrane potential rather than by specific peptide properties, as G_10_ formed pores under the same transmembrane potentials as the R_10_. The process is initiated by the intrusion of water into the hydrophobic core of the membrane, resulting in the formation of a transient water column. Subsequently, protrusion of lipid headgroups from both membrane leaflets towards the membrane center promotes the formation of a hydrophilic membrane pore. Upon entry of R_10_ into the pore, the pore diameter decreases progressively over time (Supplementary Fig. 6), highlighting the critical role of sustained transmembrane voltage in maintaining a large water pore. Notably, direct peptide translocation occurred exclusively for cationic R_10_, but not for G_10_, indicating that electrophoretic drift of highly positively charged peptides along the transmembrane electric field constitutes the primary driving force for translocation (Fig. 3a). The translocation mechanism observed during simulations is exemplarily illustrated in Fig. 3c and Supplementary video 2.

**Fig. 3.**
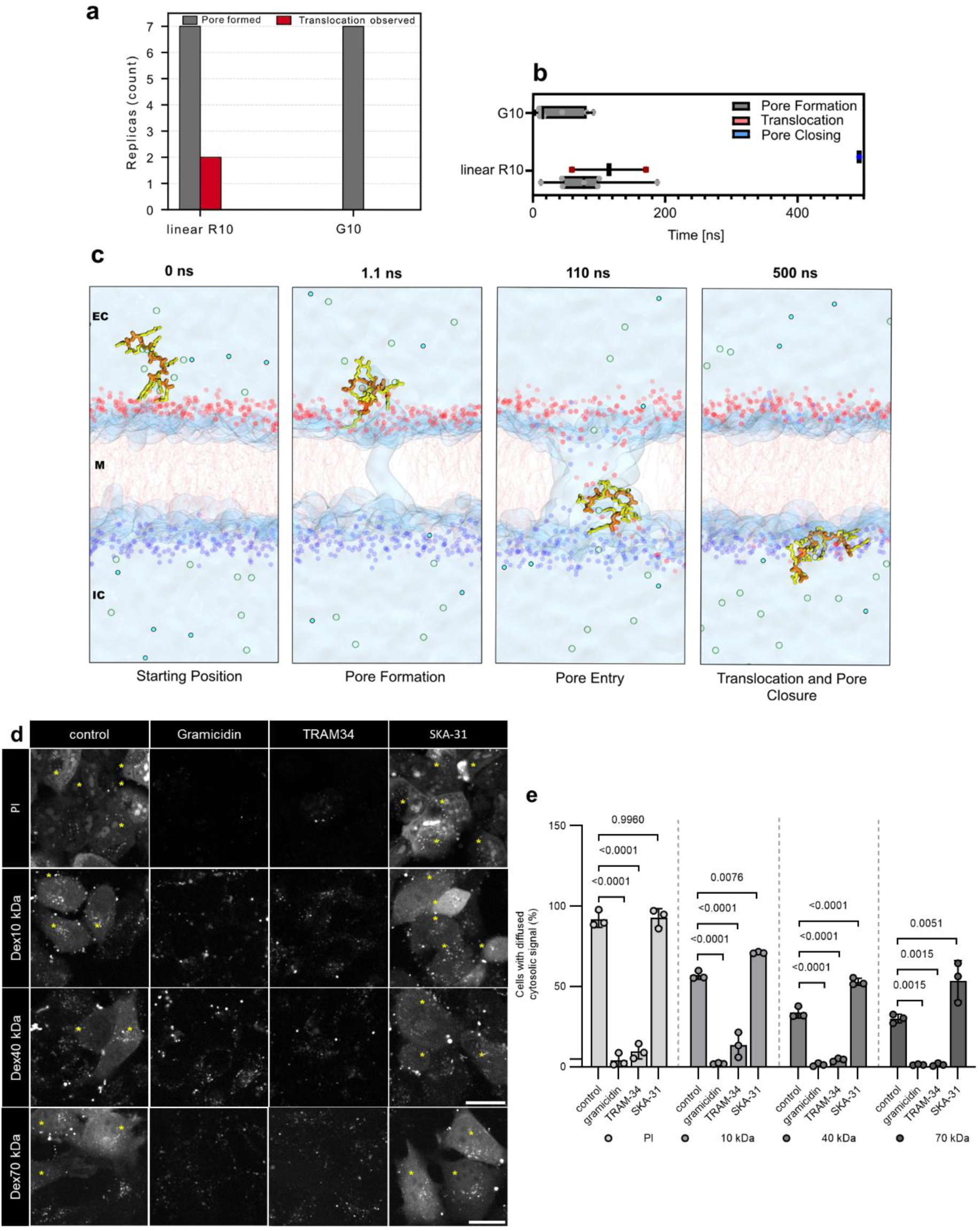
Pore formation and CPP translocation pathway revealed by atomistic MD simulations and fluid marker studies. a) Summary of pore formation and CPP translocation events observed across seven independent simulation runs of 500 ns. b) Mean times for pore formation, CPP translocation, and pore closure during simulations, shown as bar plots. Individual data points correspond to single simulation replicas, and error bars denote the standard deviation. c) Representative translocation mechanism of linear R_10_ observed in the simulations. d) Representative confocal images of HeLa cells pre-treated with 1 µg/mL gramicidin, 10 µM of TRAM-34 or SKA-31, incubated with 50 µg/mL propidium iodide (PI), 400 µg/mL 10 kDa, 40 kDa, 70kDa neutral TMR-dextran with 5 µM CPP-additive **1** for 15 min in serum free FB DMEM. Yellow asterisks indicate cells with diffuse cytosolic signals. Scale bar = 20 µm. Incubation time was reduced to 15 min since direct translocation was experimentally found to happen quickly and longer incubation time only increases background signal through endosomal uptake. e) Quantification of the cells with diffused cytosolic signal (see Supplementary Information). Data presented as mean ± SE of three biological replicates. *P*-values were determined by one-way analysis (ANOVA) followed by Dunnett’s post hoc test.

We further quantified the minimum pore diameter (defined as the narrowest diameter of the pore), which was previously suggested to be around 2 nm.^[31]^ Interestingly, the largest minimum pore size was observed for G_10_ (Extended Data Fig. 3c), which did not translocate and consequently did not transiently occupy the pore. Nevertheless, this difference vanishes when considering the maximal pore size, with no notable differences observed between the CPP-cargoes, all of which consistently exhibit maximal pore diameters of approximately 3.5−4 nm (Supplementary Fig. 7). The larger pore sizes observed here compared to earlier reports likely arise from the continuous application of transmembrane voltages. Following peptide translocation, pore closure was observed in most of the 500 ns simulations (Fig. 2b). We attribute that the reduction in membrane potential caused by partial dissipation of the ion imbalance after the charged peptides translocated between the two compartments induced the pore closing event (Supplementary Fig. 8).

Overall, the MD simulations reveal that transient pore formation is primarily driven by the applied transmembrane potential. The data also indicates that the positive charge on the CPP is the driving force for translocation through the water pore, with the transmembrane electric field acting as a selectivity filter for positively charged cargoes. Moreover, we observed computationally a relatively large pore size of up to 4 nm, which would allow the transport of small protein cargos.

Next, we experimentally validated the pore size using fluid-phase markers of defined hydrodynamic radius. We incubated HeLa cells with different fluid markers such as propidium iodide (PI, small fluorescent molecule, molecular weight = 668 Da) and TAMRA-labeled neutral dextrans (10 kDa ∼ 1.86 nm, 40 kDa ∼ 4.8 nm and 70 kDa ∼ 6.5 nm)^[55]^, in the presence of CPP-additive **1** (5 µM). Confocal images of treated cells showed diffused fluorescent signals following fluid marker uptake in the presence of CPP-additive **1** (yellow asterisk, Fig. 3d), while in the absence of CPP-additive **1** no such cytosolic signal (Supplementary Fig. 10) was observed. The signal intensity and the proportion of cells exhibiting a diffuse cytosolic TAMRA fluorescence decreased with increasing size of the fluid marker, and increased with increasing concentrations, indicating that direct translocation of these fluid markers through transient water pores is influenced by both molecular size and concentration (Fig. 3d-e, Supplementary Fig. 11). In addition, modulation of V_m_ influenced the delivery efficiencies of all fluid-phase markers as previously observed for qCPP **4** uptake; hyperpolarization enhanced and depolarization reduced the number of cells with diffused cytosolic signals, and complete depolarization (V_m_ = 0 mV) prevented the uptake of all fluid markers (Fig. 3d-e). We conclude, that exogenous cargos up to 6.5 nm size can translocate into cells in presence of CPP-additive **1**.^[55–57]^ Furthermore, fluid markers only translocated in polarized cells (V_m_ < 0) in the presence of CPP-additive, strengthening our hypothesis on direct translocation through water pores driven by temporary hyperpolarization of the membrane.

### Direct translocation of CPP-modified proteins

We demonstrated the interplay of nucleation zones, membrane potential and water pore formation for CPP-peptide cargoes. To investigate whether CPP-modified proteins enter cells through the same mechanism as qCPP **4** in the presence of CPP-additive **1**, we first employed the green fluorescent protein-binding nanobody GBP1 (∼ 15 kDa) in HeLa cells. Nanobodies in general represent a class of small protein-binders that can be generated with high affinity and specificity for a given target, making them attractive for divers intracellular therapeutic applications.^[22, 58]^ However, in this case, GBP1 was chosen as model protein due to the lack of specific intracellular accumulation and localization sites in endogenous non-GFP containing cells, acting as a reference protein for direct translocation. GBP1 **A** was expressed and labeled with TAMRA (GBP1 **B**-TAMRA) using an established sortase-mediated ligation protocol and further modified with CPPs using BioRAM (TAMRA-GBP1 **C**, Extended Data Fig. 4a-b). BioRAM is a recently developed approach that enables the straightforward, reversible modification of native amines, allowing efficient cytosolic delivery while preserving protein integrity.^[21]^

First, we tested the fluorescently labeled, CPP-modified GBP1 **C** in a time-lapse microscopy set up. We hardly observed development of fluorescent signal from nucleation zones formed by the Cy5-CPP-additive **2**, due to the high background fluorescence, that was also persistent for the non-quenched fCPP **3** (Supplementary Fig. 12). We, therefore, adopted the concept of a quenched fluorophore onto the protein GBP1 **A**, to gain selective activation of the fluorescence upon successful intracellular protein delivery. TAMRA-ER-C(BHQ-2)-ERERLPETGG peptide **12** was *N*-terminally ligated to GBP1 **A** using sortase-mediated ligation yielding quenched GBP1 **D**, which was further modified using BioRAM-R_8_ resulting in the cell-permeable, quenched GBP1 **E** (Fig. 4a, Extended Fig. 5). Addition of 2.5 µM quenched, cell permeable GBP1 **E** in the presence of 2.5 µM Cy5-CPP-additive **2** resulted in immediate protein entry into the cell, followed by reductive cleavage of the Quencher yielding fluorescent GBP1(Fig. 4a, Supplementary Video 3). During time-lapse microscopy we observed positions within the confocal layer that temporarily show high fluorescent signal, followed by diffusion into the entire cell, similar to the peptide probe qCPP **4** (Fig. 4b-c). These points of entry correlate with CPP-additive **2** nucleation zone signals. The fluorescent signal of fluorescent GBP1 **E** is markedly reduced when cells are depolarized by preincubation with gramicidin.

**Fig. 4.**
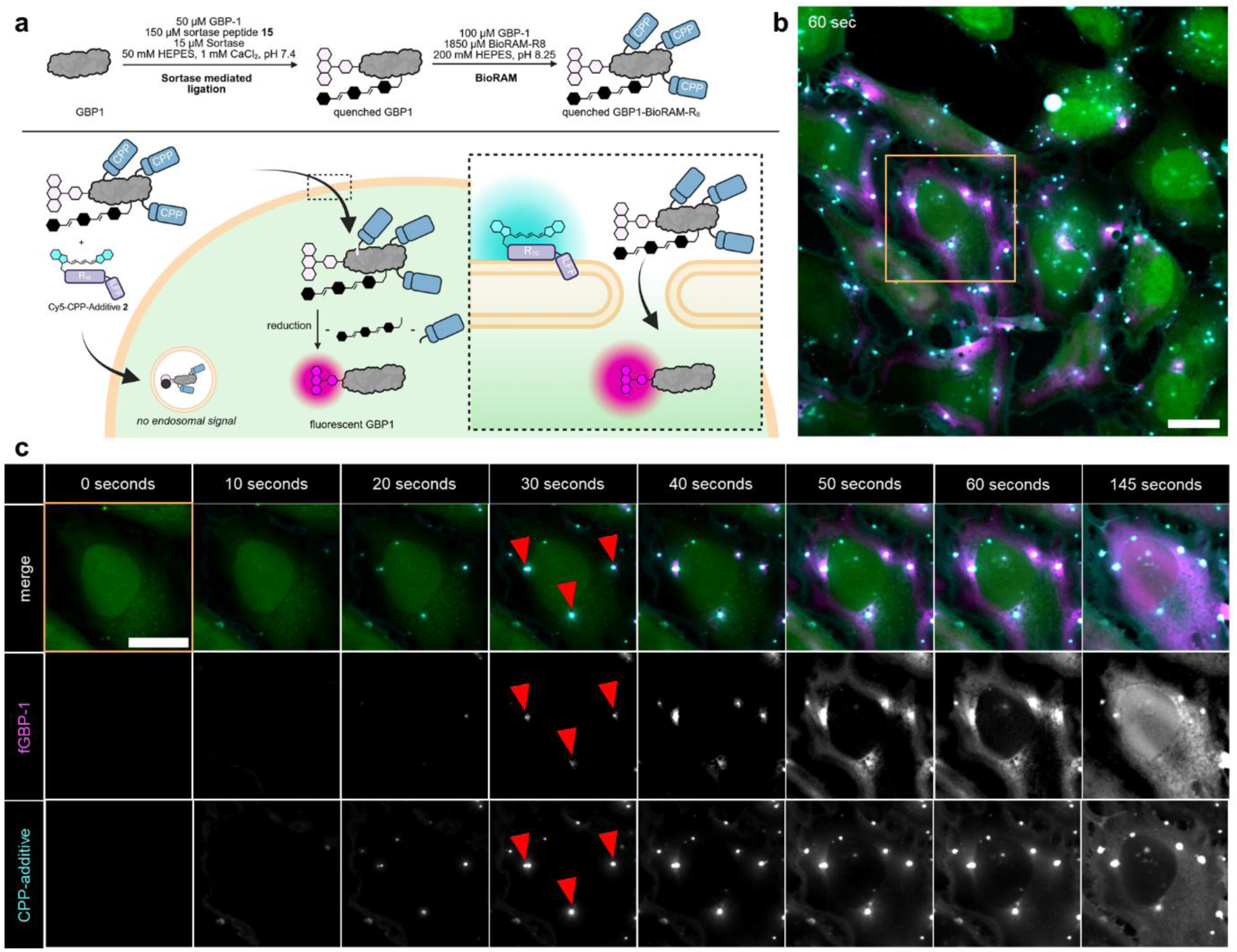
Live tracking of protein entry. a) Schematic representation of quenched GBP1 **E** uptake and intracellular activation of fluorescence. (Extended Data Fig. 5). b) Time lapse microscopy experiment monitoring cellular entry of quenched, CPP modified GBP1 **E**. Composite image (green: cytosolic marker CMFDA, magenta: GBP1 **E**, cyan: Cy5-CPP-additive **2**) of the 60 seconds frames of HeLa cells treated with 2.5 µM GBP1 **E** and 2.5 µM of Cy5-CPP-additive **2**. c) Highlighted region of interest from the time lapse microscopy from section C in 10 second intervals. Nucleation zones, which serve as point of entry for GBP1 **E** are marked as red arrows. Scale bar = 20 µm.

To enable quantitative analysis of time dependency and membrane potential effects on protein delivery via fluorescence microscopy, we used BioRAM-conjugated NLS-mCherry (NLS-mCherry **G**) as a model cargo. NLS-mCherry was chosen because it is intrinsically fluorescent, easy to express, and highly stable, eliminating the need for additional chemical labeling and providing a robust read out. Upon cytosolic delivery, the NLS-mCherry **G** construct localizes to the nucleus due to the genetically encoded nuclear localization sequence (NLS), which facilitates high throughput quantification using our optimized microscopy protocol by directly translating the fluorescent nuclear signal into delivery efficiency (Supplementary methods section NLS-mCherry-BioRAM uptake).^[21]^

We first analyzed the time dependent CPP-additive mediated uptake of NLS-mCherry **G** over 5, 15, 30, 60, and 90 minutes by measuring total cellular uptake while differentiating endosomal and nuclear fluorescence (Fig. 5b and 5c, Supplementary Fig. 13). Protein uptake via direct translocation (here, nuclear fluorescence) was complete within the first 5 minutes showing no significant increase thereafter, consistent with the rapid entry and early saturation observed in the time-lapse microscopy. In contrast, endosomal signals progressively increased over longer incubation time, plateauing after 60 minutes of incubation. The constant nuclear fluorescence beyond 5 min further indicates that endosomal escape makes no contribution to cytosolic delivery within this timeframe, in line with our previous observation that BioRAM-proteins delivered using CPP-additives **1** do not induce endosomal rupture.^[21]^ Next, we explored the membrane potential influence on the CPP-mediated uptake of proteins. We incubated untreated, gramicidin-, TRAM-34-, and SKA-31-treated HeLa cells with CPP-additive **1** and NLS-mCherry **G** for 15 min and performed quantification of fluorescent signal within the nucleus (Fig. 5d, e). Both qualitative confocal microscopy images and automated quantitative microscopy^[21]^ showed higher fluorescence intensity in HeLa cells treated with hyperpolarization agent SKA-31, while depolarizing cells with TRAM-34 and gramicidin show reduced uptake. To determine whether the direct translocation of NLS-mCherry **G** linearly correlates with V_m_, we performed the protein uptake experiment in 20 mM HEPES buffer with increasing KCl concentration, which allows proportional modulation of cell membrane potential (Extended Data Fig. 6b).^[31, 39, 54]^ In accordance with previous reports, increasing KCl concentration led to depolarization, which also led to decreased nuclear fluorescence signal (Extended Data Fig. 6b). Finally, we assessed the direct translocation of NLS-mCherry **G** in different cells lines with varying resting V_m_ (Fig. 5f). While there is a good correlation between membrane potential and protein uptake within individual cell lines HeLa, U2OS, A549 (Fig. 5h, Extended Data Fig. 6b-d), it is difficult to cross-correlate resting V_m_ with protein uptake between different cell lines (Fig. 5g). Each cell line differs in properties, such as lipid membrane composition^[59]^, glycocalyx complexity^[29, 30]^, and cell-surface proteins^[60]^, which may contribute the susceptibility of a specific cell line for direct translocation. Nonetheless, V_m_ emerges from our data as a critical determinant of direct translocation efficiency of proteins.

**Fig. 5.**
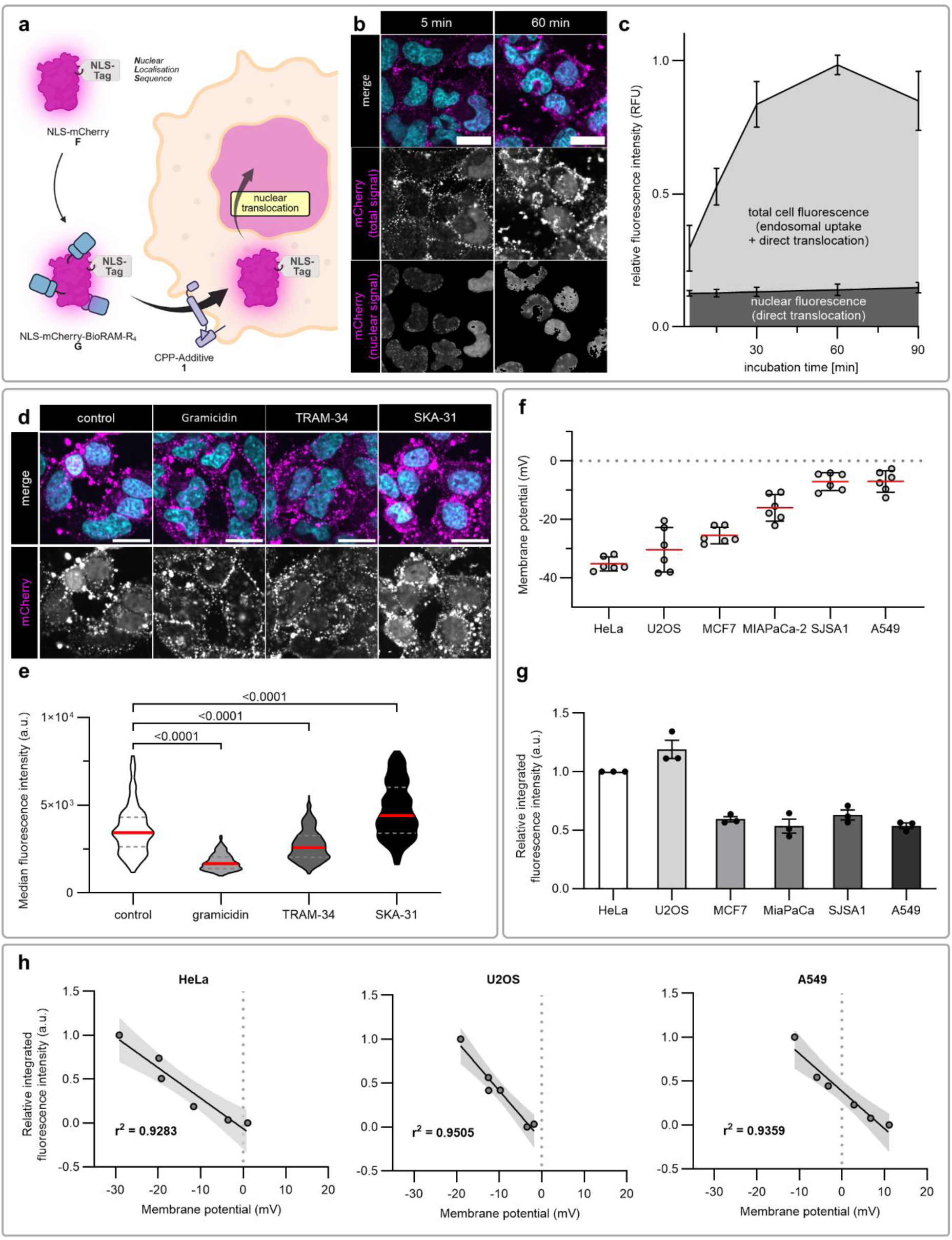
Protein uptake is governed by membrane potential across cell lines. a) Schematic representation of NLS-mCherry protein delivery. BioRAM modification and CPP-additive mediated translocation followed by nuclear translocation of the protein. b) Representative microscopy images of HeLa cells after delivery of NLS-mCherry **G** for the quantitative assessment in (c). Incubation of HeLa cells with 2.5 µM NLS-mCherry **G** and 5 µM CPP-additive **1** in serum free FB DMEM. Nuclei co-stained with Hoechst (cyan). Scale bars = 20 µm. Full panel is displayed in Supplementary Fig. 13. c) Quantification of total intracellular signal over incubation time and the subfraction of nuclear fluorescence of intact protein as a result of direct translocation. Data is represented as mean ±SD of triplicate of independent biological replicates, 200 cells each. d) Microscopy images showing uptake of 2.5 µM NLS-mCherry **G** and 5 µM CPP-additive **1** in serum free FB DMEM in HeLa cells that were pretreated with membrane potential modulators. Pretreatment using of 1.2 µg/mL gramicidin, 20 µM TRAM-34 or 20 µM SKA-31 in serum free FB DMEM for 5 min. Nuclei co-stained with Hoechst (cyan). Scale bars = 20 µm. e) Representative violin-plot quantifying the nuclear fluorescence intensities of around 270 cells from images from (d). Data shows median value (red line) and quartile values (grey line). *P*-values were determined by one-way analysis (ANOVA) followed by Dunnett’s multiple comparison test. f) Membrane potential determined using by DIBAC_4_(3) and FACS. Data shows individual readings across three biological replicates with mean values (red line) and SD. g) Quantification of intracellular delivery of NLS-mCherry **G** in the presence of 5 µM CPP-additive **1** for 15 min. h) Correlation of membrane potential with delivery efficacy within respective cell lines. Data fit by linear regressions and 95% confidence intervals (gray filled area) and corresponding coefficient of determinations (r^2^). Replicate data for (e) and Complete set of data for (h) are found in Extended Data Fig. 4.

Finally, we tested the hypothesis, derived from our computational simulations, that CPP-additive mediated delivery is selectively for positively charged cargos (Fig. 4). To investigate this, we compared fluorescence intensity of cells treated with neutral TAMRA-labeled fluid markers and the CPP-modified, positively charged TAMRA-GBP1 **D** at 50 µg/µL in the presence of 5 µM CPP-additive **1**. Despite smaller size and higher labeling (1-2 dyes/10 kDa dextran) of 10 kDa dextran (according to manufactures Manual “Dextran Conjugates”, mp01800, Thermo Fisher), cells treated with GBP1 **D** (1 dye/protein) showed 50−70-fold higher fluorescence (Fig. 6a), indicating that positive charge strongly potentiates uptake. To extend this concept, we tested for unspecific co-delivery of bystander proteins. To this end, we fluorescently labeled GBP1 with the spectrally distinct FITC (FITC-GBP1 **H**) and modified it with CPPs using BioRAM (FITC-GBP1 **I**) (Extended Data Fig. 4). We incubated cells with 2.5 µM fluorescently labeled GBP1 (TAMRA-GBP1 **B,** FITC-GBP1 **H**) together with the cell-permeable spectral counterpart (TAMRA-GBP1 **C,** FITC-GBP1 **I**) in the presence of 5 µM CPP-additive **1** (Fig. 6b). Besides endosomal uptake, we only observed direct translocation (here, diffused cytosolic fluorescent signal) for the CPP-modified GBP1 **C** and **I**. To further challenge our system for nonspecific protein uptake, we tested the delivery of CPP-modified GBP1 **C**/**I** (2.5 µM) in the presence of 20-fold excess of non CPP-modified GBP1 **B**/**H** (50 µM). We only observed endosomal uptake and no evidence of direct translocation of bystander proteins (Supplementary Fig. 14). These findings demonstrate that conjugation of CPPs to the cargo protein markedly enhances cellular uptake. Non-specific delivery of unmodified proteins via CPP-additive-induced water pores was not detected under our conditions, though such bystander delivery effects may become more relevant at higher concentrations (> 50 µM of protein) or smaller cargoes.

**Fig. 6.**
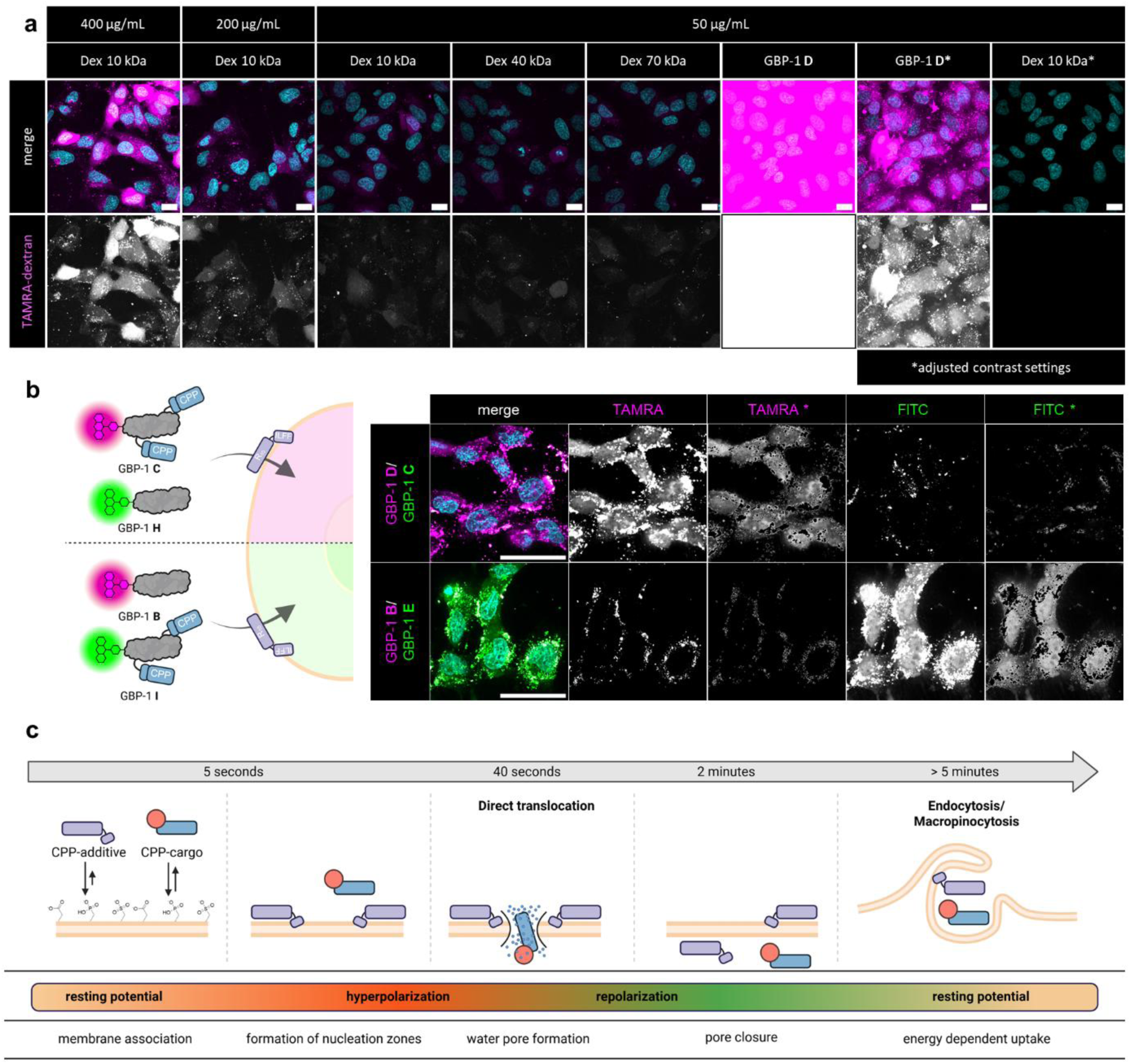
Specific CPP-protein uptake and mechanistic Framework. a) Representative confocal images of HeLa cells treated with TAMRA labeled dextran of different size and concentration with 5 µM CPP-additive **1** for 15 min in serum free FB DMEM. Comparison to treatment with CPP modified TAMRA-GBP1 **C**. Intracellular signal is 50−70-fold increased comparing GBP1 **C** over 10kDa dextran. b) Co-delivery of fluorescently labeled CPP-modified GBP1 (**C** and **I**) over non-CPP-modified GBP1 (**B** and **H**) at 2.5 µM each with 5 µM CPP-additive **1** for 15 min in serum free FB DMEM. Nuclei were counterstained with Hoechst. (green: GBP1 **H**/**I**, magenta: GBP1 **B**/**C**, cyan: nucleus). *endosome exclusion was performed using a Fiji macro by thresholding of high-intensity signal. Scale bars = 20 µm. c) Schematic representation of the proposed 5-step mechanism for CPP-additive mediated direct translocation of CPP-cargo.

Taken together, we conclude that CPP-additive-mediated delivery of both peptide and protein proceeds through membrane potential driven water pores within nucleation zones, with specificity for CPP-modified proteins.

### Mechanistic framework for the direct translocation of proteins

Based on our experimental findings, we propose the following five-step model for CPP-additive-mediated direct translocation of CPP-cargo (peptide/protein) (Fig. 6c).

CPP-additives initially associate with the cell surface through ionic interactions of its cationic arginine residues with the negatively charged membrane membrane surface (step 1).^[61]^ Being optimized for enhanced membrane retention through its hydrophobic anchor and thiol-reactive moiety, CPP-additives then promote the formation of nucleation zones (step 2).^[19, 20, 42]^ These nucleation zones appear as discrete punctate regions on the plasma membrane, exhibiting markedly longer residence times compared to CPP-cargo conjugates. Accumulation of CPP-additive positive charges on the membrane induces a temporary hyperpolarization event by locally increasing the membrane potential, which subsequently drives the formation of transient water pores (step 3; pore size: simulation up to 4 nm, experimental up to 6.5 nm). CPP-cargo (peptides or protein) enters the cell primary through these water pores seconds after addition to the cells. The subsequent influx of positively charged molecules restores the cell’s resting membrane potential, leading to the closure of the water pore and termination of direct translocation (step 4). Thereafter, remaining extracellular CPPs are internalized via endocytosis at much longer timescales. (step 5).

#### Step 1 – Initial membrane association

In line with previous studies on arginine-rich CPPs, our data support an initial binding step dominated by electrostatic attraction between the polycationic CPP-additives and anionic components of the plasma membrane.^[27, 62–65]^ This interaction concentrates CPP-additives on the membrane surface and is a prerequisite for subsequent spatial organization. Although our simulations employ compositionally simplified lipid bilayers, prior MD work has demonstrated that lipid composition and surface charge density strongly modulate membrane stability and the free energy barrier for peptide binding and insertion.^[66]^ We therefore anticipate that local membrane defects further promote CPP-additive accumulation and primes specific regions of the cell surface for nucleation zone formation.^[19, 39]^

#### Step 2 – Nucleation zone formation

Once bound, CPP-additives are designed to remain at the membrane via a synergy of electrostatic interactions, hydrophobic contacts, and thiol-anchoring to membrane-proximal cysteine residues or thiol-reactive motifs. This multivalent retention promotes additive clustering into nucleation zones, providing highly concentrated CPP patches that are observed as discrete, long-lived puncta on the cell surface. Such clustering is consistent with previous reports that multimerization^[15]^, co-delivery^[67]^, or formation of peptide-rich coacervates^[68, 69]^ enhances CPP-mediated uptake by surpassing local concentration thresholds required for efficient translocation.^[70]^ In our framework, CPP-additive-rich nucleation zones distinguish from more uniformly distributed CPPs that remain below the critical local density.

#### Step 3 – Hyperpolarization and water pore formation

The build-up of positive charge within nucleation zones is expected to induce a pronounced local hyperpolarization, previously also termed “megapolarization” (V_m_ ≤ –150 mV)^[31]^ for CPP-induced pore formation. While microscopy analysis report only changes about ΔV_m_ ∼ 20 mV in the apparent V_m_, these likely reflect an average over multiple nucleation zones on multiple cells within the region of interest. We propose that accumulation of CPP-additives within individual nucleation zones can temporarily drive local membrane potential toward values sufficient to lower the energetic barrier for the formation of water pores, reminiscent of what happens during electroporation.^[71]^

#### Step 4 – Direct translocation of CPP-cargo and pore closure

Once water pores form, positively charged CPP-cargo molecules experience a strong inward electrophoretic driving force due to their net positive charge, favoring entry through these transient openings. The efficient cytosolic delivery of CPP-modified proteins and peptides, over neutral fluid marker (Dextran, G_10_), indicates that translocation is a charge-driven passage through defined, short-lived pores. Importantly, permanently depolarized cells show no uptake, as they can’t hyperpolarize which would allow for water pore formation. This tight interplay between temporary polarization, pore formation, and cargo translocation demonstrates, that CPP-mediated delivery is not a consequence of an uncontrolled membrane rupture. Following the initial direct translocation, influx of external cations and CPP-cargo through the water pores gradually restores the resting membrane potential, promoting pore closure and terminating further direct translocation.^[54]^ Others have identified biochemical processes linked to ion homeostasis that may also influence pore termination.^[31, 54]^

#### Step 5 – Termination of translocation and endocytic uptake

At later time points, remaining extracellular CPP-additives and -cargo fail to induce the critical hyperpolarization required for direct translocation and are therefore predominantly internalized via endocytosis, leading to punctate intracellular signals that distinct from the early cytosolic fluorescence of direct entry. This temporal segregation aligns with the commonly observed concentration-dependence of CPP uptake, in which crossing a threshold enables rapid, energy-independent translocation, whereas subthreshold conditions lead to slower, endocytosis-dominated internalization.

Our molecular dynamics simulations provide a mechanistic view of CPP-additive–induced pore formation but are inherently constrained by spatial and temporal limitations. Only small membrane patches corresponding to subsections of a nucleation zone can be probed on nano- to microsecond timescales, and the simulated compositions omit key features of the native plasma membrane, such as membrane proteins, full lipid complexity, and the precise local CPP density. Consequently, we focused our models on voltage-driven CPP–membrane interactions rather than attempting to reproduce the full biochemical environment. Previous computational studies of CPP-mediated permeabilization have adopted related strategies but differ substantially in methodology, employing either coarse-grained representations to access larger length scales or shorter all-atom simulations without maintaining ion imbalance over time.^[31, 35]^ In contrast, our approach preserves an ion gradient throughout, enabling sustained hyperpolarization and the observation of larger, more stable pores that mirror the enhanced translocation capacity of CPP-additives.

Although this work centers on a CPP-additive–based delivery platform, the mechanistic framework likely extends to other polycationic systems capable of direct membrane translocation at high local concentrations, including guanidinium-rich polymers^[72–74]^ and cationic lipid nanostructures^[75]^. In all such systems, local charge accumulation, membrane clustering, and reaching of critical concentration thresholds appear to be recurrent design motifs that promote transient pore formation and efficient cargo entry.^[9]^ Our framework suggests that engineering nucleation zones, through tailored hydrophobicity, charge density and anchoring groups, may serve as a general strategy to sensitize membranes towards reversible, membrane potential-driven pore formation. In this context, CPP-additives function not just as carriers but as modulators of the membrane potential, enabling cytosolic delivery of otherwise impermeable proteins at low treatment concentrations.

## Conclusion

We present the first mechanistic understanding for CPP-additive mediated protein delivery via direct translocation. We demonstrate rapid and selective entry of CPPs and CPP-proteins at discrete nucleation zones formed by CPP-additive accumulation without disruption of the membrane. Computational studies complement experimental evidence to support the decisive role of membrane potential and suggest the formation of transient water pores, large enough to accommodate proteins.

The uptake hinges on local, temporary hyperpolarization, requiring a critical peptide concentration threshold, which is likely achieved through the accumulation of CPP-additives on the cell membrane. This enables intracellular translocation through water pores of further CPPs/CPP-proteins along the transmembrane electric gradient at low concentrations.

The transient nature of the water pore formation as well as the enhanced entry of CPP-cargoes reinforce CPP-additive mediated direct translocation as a viable entry route for proteins besides current technologies for drug delivery. These principles possibly extend to other polycationic CPP systems and further add on the understanding of multicomponent delivery systems on a macroscopic level. To fully realize the potentials of this strategy, future works must address cell selectivity, triggered delivery and formulation strategies.

## Acknowledgment

We thank Kristin Kemnitz-Hassanin and Ines Kretzschmar for their technical assistance with cell culture, protein expression and peptide synthesis. Furthermore, we thank Andreas L. Marzinzik (Novartis) for providing the SJSA-1 cell line. J.V.V.A. was funded by an Alexander von Humboldt Fellowship and J.F. by a Chemiefonds fellowship of the Fonds der Chemischen Industrie (FCI). C.P.R.H. and H.S. acknowledge support from the Deutsche Forschungsgemeinschaft (DFG): CRC 1449 project-ID 431232613 and CRC 1349 project-ID 387284271 to C.P.R.H. and RTG2473 ‘Bioactive Peptides’ project-ID 392923329 to C.P.R.H. and H.S. Graphical abstract and figure panels created with BioRender.com.

## Extended Data

**Extended Data Fig. 1.**
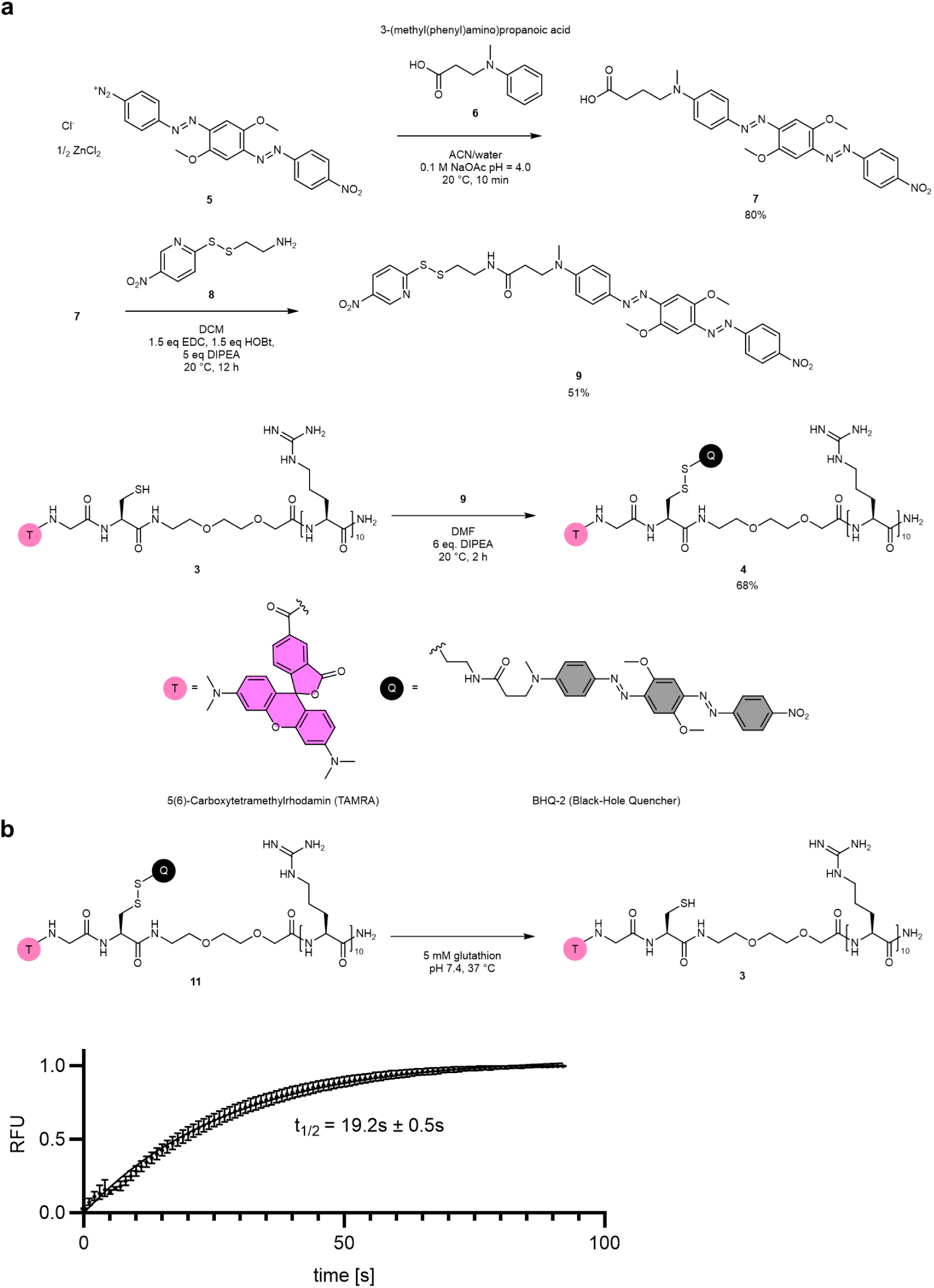
Synthesis of qCPP and *in vitro* fluorescence activation. a) Disulfide-activated BHQ-2 **9** was synthesized in two steps by azo coupling of Fast Black K diazonium salt **5** with 3-(methyl(phenyl)amino)propanoic acid **6**.^[76]^ Cysteamine was separately activated using 2,2-Dithiobis(5-nitropyridine) **8** and coupled with BHQ-2 acid **7** using EDC, HOBt and DIPEA in DCM. The peptide sequence **3** was synthesized and *N*-terminally modified by solid-phase peptide synthesis using 3 eq. TAMRA acid, 3 eq. HATU, 10 eq. DIPEA in DMF, cleaved from the resin and purified via preparative HPLC. The disulfide activated BHQ-2 **9** was then reacted to the free cysteine of the purified peptide **3** in DMF with 6 eq. DIPEA. b) We tested the reduction of the disulfide bond *in vitro* under physiological conditions, cleaving the BHQ-2 from the fCPP **4** to activate the fluorescence properties of the fCPP **4**. The speed of this activation was determined by incubation of 500 nM fCPP **4** and 5 mM glutathione (GSH) in phosphate buffered saline (PBS) at pH 7.4 at 37 °C mimicking the reducing, cytosolic environment in cells. After addition of glutathione the fluorescence (ex.: 552 nm; em.: 579 nm) was measured every second over 200 seconds. Half of the fluorescence activation was reached after 19 seconds. The increase of fluorescence of the quenched state versus the fully activated state (plateau) corresponds to 60-fold. The fast speed of the activation and the efficient quenching in the quenched state renders the fCPP **4** suitable for intracellular application including tracking of cellular uptake of the peptide probe by activation of fluorescence with high temporal resolution. Data is represented as mean ± SD of three replicates

**Extended Data Fig. 2.**
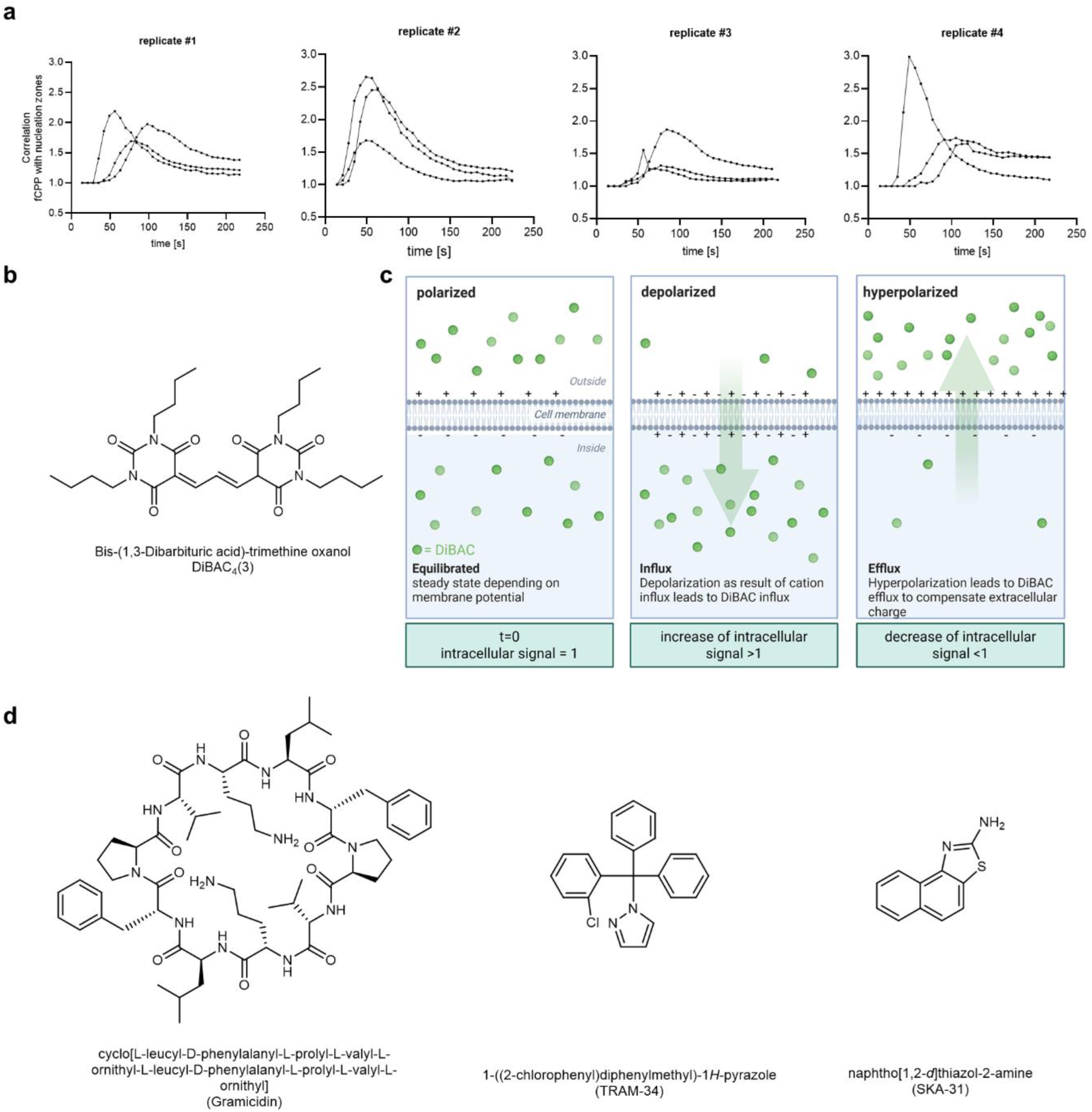
Colocalization of fCPP and CPP-additive signal over time and live-cell assessment of membrane potential. a) Colocalization of fCPP **3** and CPP-additive **2** signal in time-lapse confocal microscopy. Shown are replicates 1-4 from the correlation experiment quantifying the fCPP **1** fluorescence (RFP channel) with regions of intense Cy5-CPP-additive **2** signal relative to area outside of those regions (=correlation factor; see Supporting information). A correlation factor >1 indicates preferential colocalization of fCPP with CPP-additive–defined nucleation zone. The ratio of fCPP intensities inside vs. outside these regions was plotted over time for individual fixed-position images. Each replicate corresponds to an independently treated cell sample. b) Chemical structure of DiBAC_4_(3) as membrane potential indicator. c) schematic representation of the DiBAC_4_(3) dye for indirect membrane potential measurements. The distribution of DiBAC_4_ (3) depends on Vm gradient described by the Nernst equation, where depolarization (Vm = 0) results in the uptake of the dye and its binding to intracellular proteins or membranes exhibiting a red spectral shift facilitating microscopy analysis. Conversely, membrane hyperpolarization (Vm < resting membrane potential) is indicated as a decrease in DiBAC_4_ (3) fluorescence d) Chemical structures of the membrane potential modulators used in this study.

**Extended Data Fig. 3.**
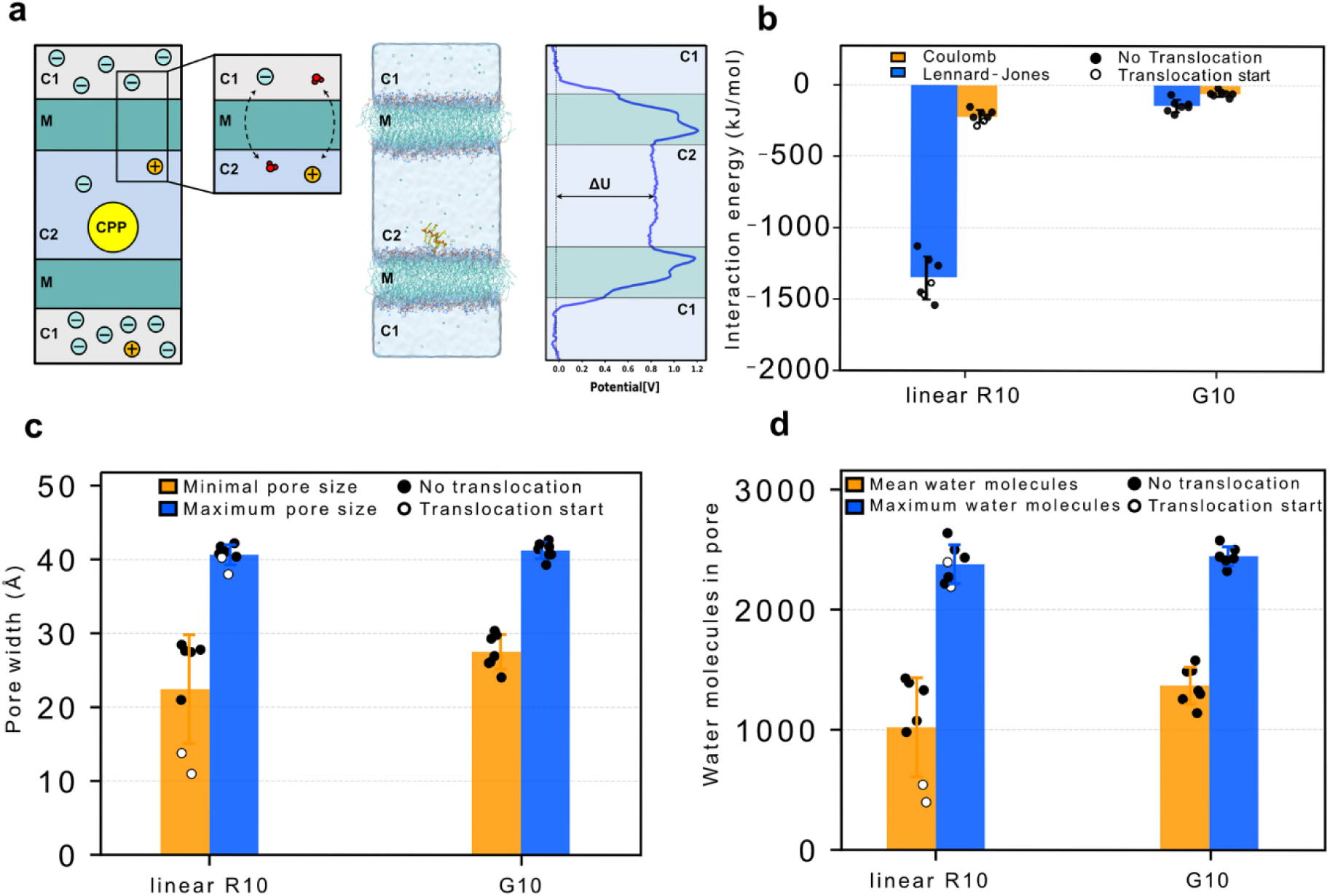
Computational characterization of CPP-membrane interactions and pore dynamics under a transmembrane voltage. a) Computational setup of the two-membrane system, in which ions are separated by two membranes (M) into compartments C1 and C2. The CPP is shown in yellow. The ionic imbalance is maintained using the CompEL protocol according to Kutzner et al.^[45]^, and the resulting transmembrane potential (ΔU) arises from the ion imbalance between C1 and C2. b) CPP-membrane interaction energies decomposed into Coulombic and Lennard-Jones (LJ) contributions, extracted from the simulations and grouped according to whether CPP translocation occurred. c) Minimum and maximal pore radii in Å and d) mean and maximal water occupancy, quantified as the number of water molecules within the pore per nanosecond, derived from seven independent 500 ns simulation runs. Simulations runs in which peptide translocation occurred are indicated by black circles, whereas runs without translocation are shown as white cycles. Bars denote averages across the seven simulations, with individual data point shown together with the standard deviation

**Extended Data Fig. 4.**
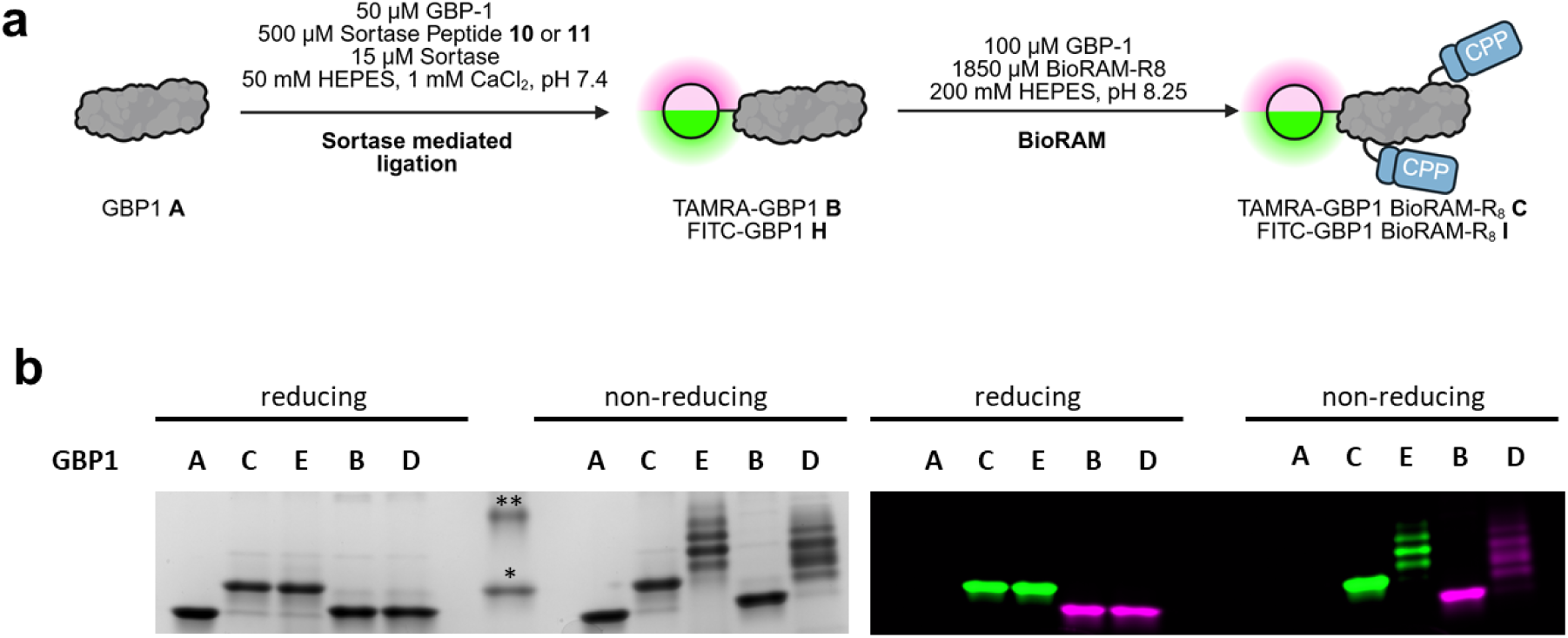
Generation of fluorescently labeled, cell permeable GBP1 nanobodies. a) Chemoenzymatic modification of GBP1 **A** by sortase yielding GBP1 **B**/**H**, followed by BioRAM modification creating homogenous labeled GBP1 **C**/**I** with an average modification of 2.2−2-5 R_8_ per protein. Fluorescently labled GBP-1 presents three solvent-exposed lysines that can be modified using the amine-selective bioconjugation strategy BioRAM (*N*-terminus is fluorescently labeled, therefore inaccessible). BioRAM-R_8_ ensures high cell permeability even at low modification degree and therefore was chosen over the shorter BioRAM-R_4_/R_6_. b) SDS-PAGE analysis of GBP1 proteins generated. Reducing samples include β-mercaptoethanol causing the reductive cleavage CPP-modifications from the protein. Coomassie stained gel (left) and fluorescent gel (right). **15 kDa, *10 kDa, full gel image is displayed Supporting Fig. S15.

**Extended Data Fig. 5.**
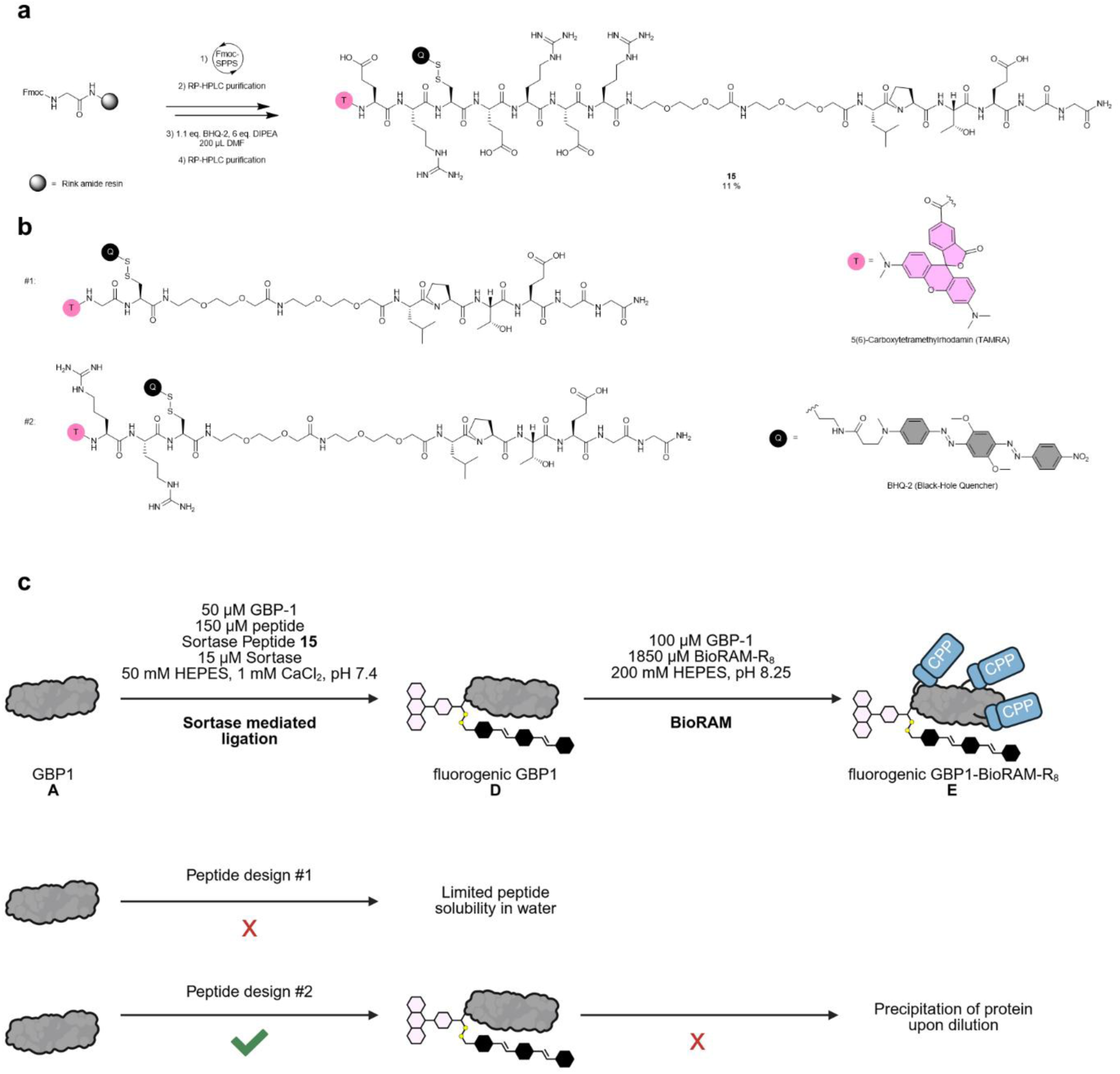
Generation of cell-permeable and quenched GBP1 **E**. a) Synthesis of peptide (TAMRA-ER-C(BHQ-2)-ERERLPETGG) for sortase mediated ligation following general procedure for solid phase peptide synthesis (SPPS) (for detailed information refer to supporting information). b) Previous designs that were synthesized and tested for chemoenzymatic labeling of GBP1 **A**. c) Chemoenzymatic modification of GBP1 **A** by sortase yielding GBP1 **D**, followed by BioRAM modification creating homogenous labeled GBP1 **E** with an average modification of 2.92 R_8_ per protein. GBP-1 presents three solvent-exposed lysines that can be labeled using the amine-selective bioconjugation strategy BioRAM. BioRAM-R_8_ ensures high cell permeability even at low modification degree and therefore was chosen over the shorter BioRAM-R_4_/R_6_. Labeling using earlier designs of the sortase peptide lacked solubility for sortase modification (#1: TAMRA-C(BHQ-2)-LPETGG) or reduced stability of the protein leading to precipitation of the bioconjugate (#2: TAMRA-RR-C(BHQ-2)-LPETGG) most likely due to solvation effects. Incorporation of six hydrophobic amino acid with balanced charge distribution led to a highly soluble peptide and stable bioconjugates.

**Extended Data Fig. 6.**
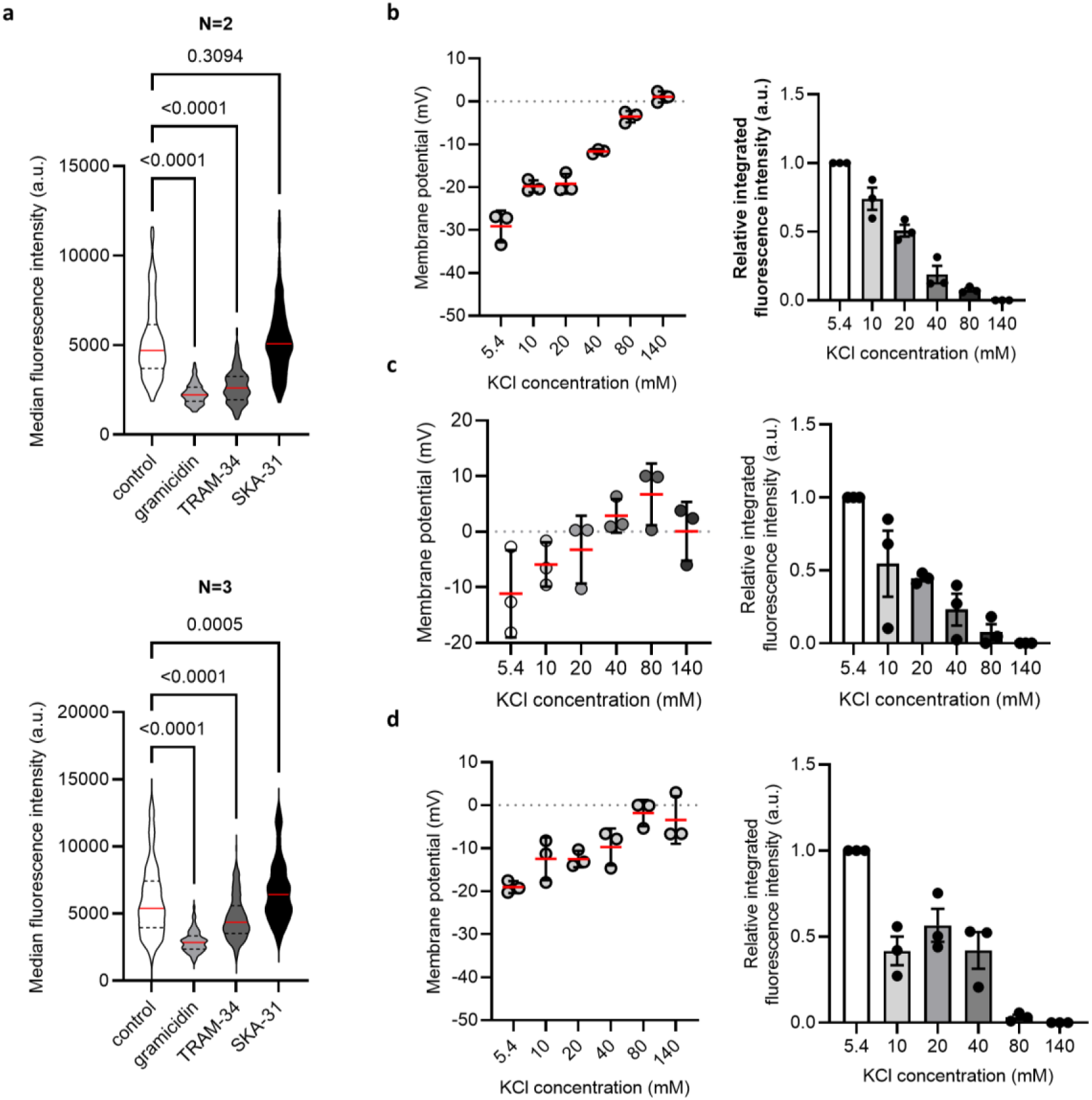
Cell membrane potential correlation with protein delivery efficacy. a) Independent replicate data (N=2, N=3) of the NLS-mCherry **G** uptake in HeLa cells pretreated with gramicidine, TRAM-34, and SKA-31. Red line indicates the mean and dotted gray lines the quartile values. *P*-values are calculated by one-way ANOVA with Dunnet’s post hoc analysis. (b-d) Membrane potential modulation at different KCl concentration. (left) DiBAC-based membrane potential measurement using FACS and (right) NLS-mCherry **G** uptake in b) HeLa cells, c) A549, and d) U2OS cells. Data presented as mean ± standard error of three independent experiments.

