## Supplementary Information for "Cellular protein delivery through membrane potential driven water pores"

---

[1] Jonathan Franke, Palina Dubatouka, Amin Yourdkhani, Tillmann Utesch, Sahil Soni, Juliana Serrano, Tolga Soykan, Martin Lehmann, Jan Vincent V. Arafiles, Han Sun, Christian P. R. Hackenberger
Leibniz-Forschungsinstitut für Molekulare Pharmakologie (FMP)
Robert-Rössle-Strasse 10, 13125 Berlin (Germany)


[2] Jonathan Franke, Palina Dubatouka, Christian P. R. Hackenberger
Department of Chemistry, Humboldt Universität zu Berlin
Brook-Taylor-Straße 2, 12489 Berlin (Germany)

[3] Amin Yourdkhani, Sahil Soni, Han Sun
Institut für Chemie, Technische Universität Berlin
Strasse des 17. Juni 124, 10623 Berlin (Germany)

[4] Juliana Serrano
Institute of Chemistry and Biochemistry, Freie Universität Berlin
(Germany)

### Supplementary figures

#### Supplementary Fig. 1. Time-lapse experiment using fCPP 3.

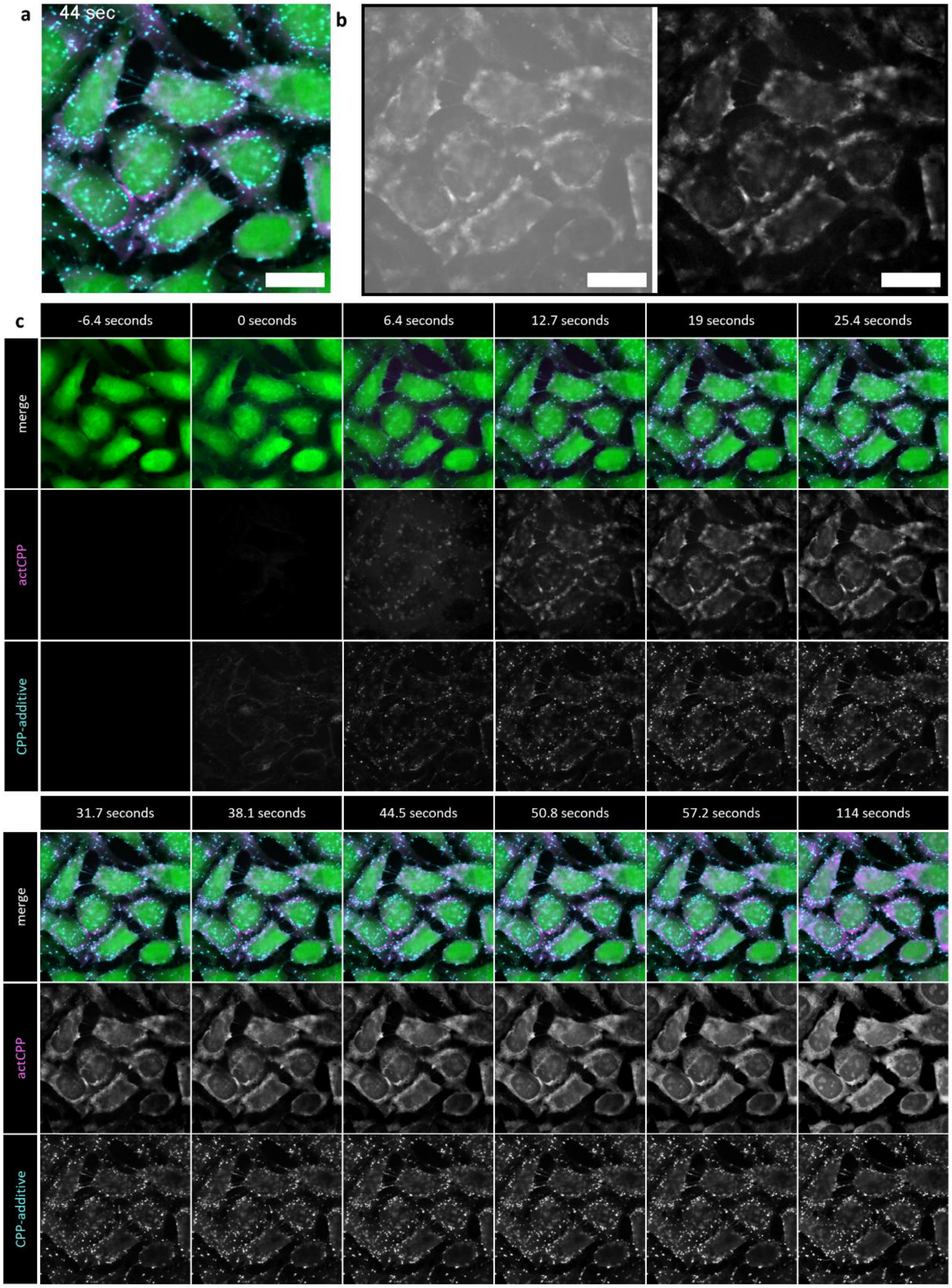

HeLa cells were incubated with 2.5 µM of fCPP **3** and 2.5 µM of Cy5-CPP-additive **2** in serum-free FluoroBrite DMEM at 37 °C, 5% $CO_2$ and imaged every 6.35 seconds. Scale bar = 20 µm. (green – cytosolic marker CMFDA, magenta – fCPP **3**, cyan – Cy5-CPP-additive **2**). a) Representative composite image at 44 seconds. b) RFP (fCPP) channel at 19 seconds, highlighting the unadjusted (left) adjusted (right) contrast to account for high background fluorescence. c) Time-lapse montage of the incubation series.

#### Supporting Fig. 2. Toxicity assessment of qCPP 4 on HeLa cells.

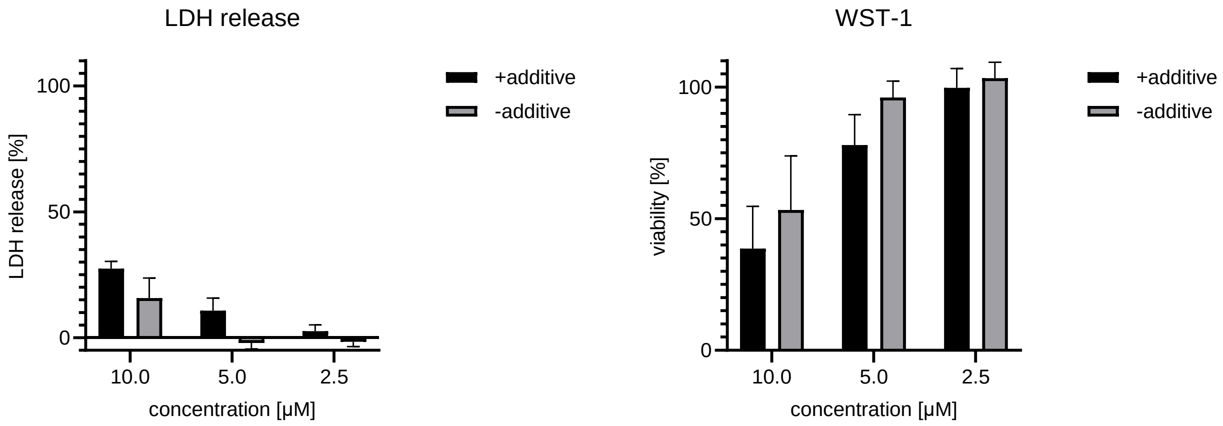

HeLa cells were treated with 10, 5, 2.5 µM of qCPP **4** in the presence and absence of 5 µM CPP-additive **1**. After 1 hour the cell supernatant was tested for lactate dehydrogenase (LDH) activity using a membrane integrity assay kit from Promega. After 24 h viability was tested by measuring mitochondrial dehydrogenase activity WST-1. Data is represented as mean ±SD of three independent biological replicates (N=3).

#### Supplementary Fig. 3. Discussion of out-of-plane localization.

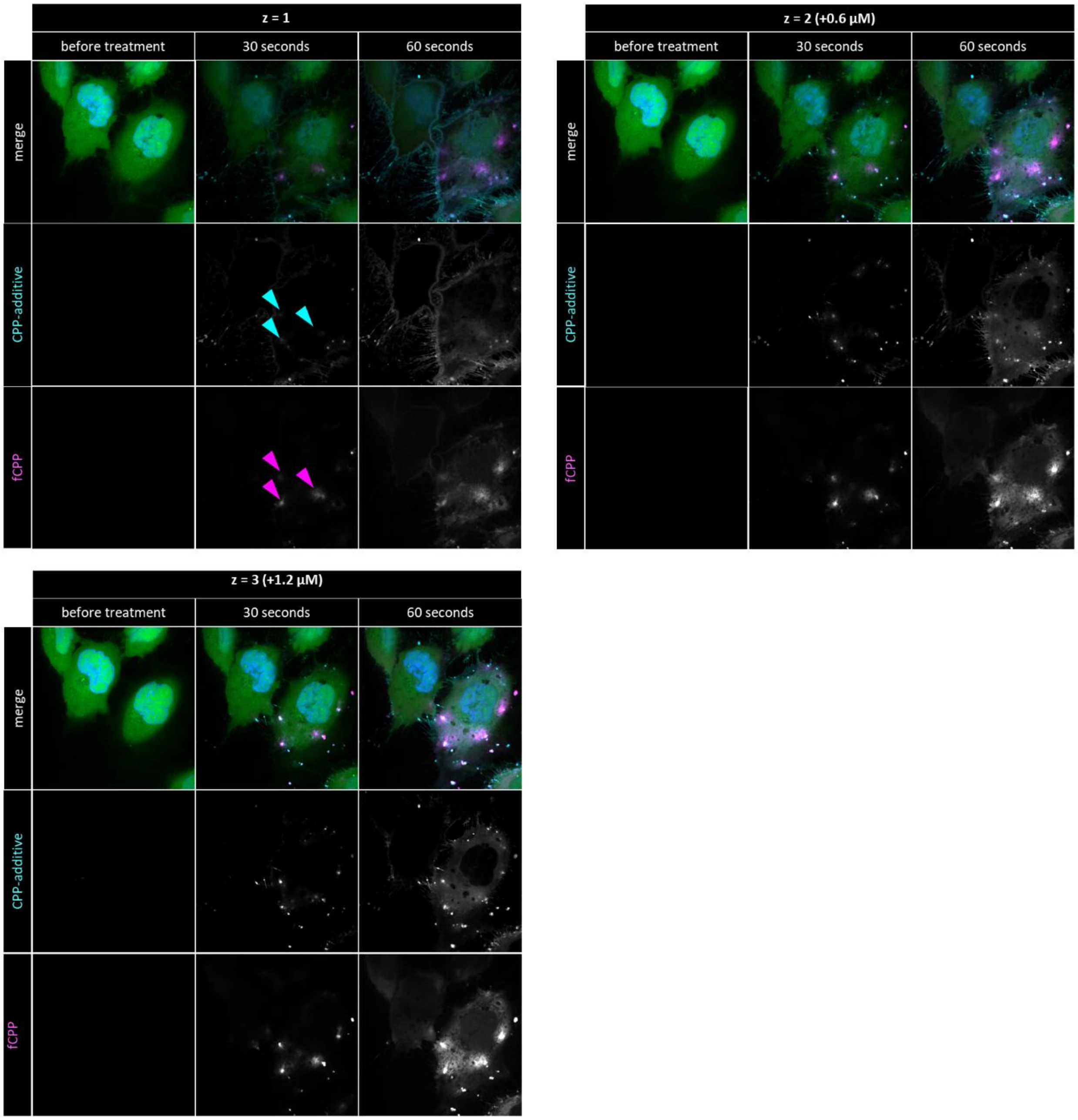

Time lapse images of CPP-additive mediated qCPP entry. HeLa cells were stained using cytosol marker (CMFDA) and nuclear stain (Hoechst). qCPP **4** and Cy5-CPP-additive **2** were added to a final concentration of 2.5 µM. Three z positions were captured for the region of interest every 30 seconds. The cyan arrows indicate the associated extracellular accumulations of Cy5-CPP-additive, which are only well resolved in plane z=3. The magenta arrows point towards influx of qCPP **4**, followed by reduction to fCPP **3** and intracellular activation of fluorescent signal visible in all z-planes.

**Discussion:**

It is important to note that the confocal fluorescence microscopy applied in this study captures only a single optical section. Within the imaged plane, we observe pronounced nucleation zones at multiple positions along the cell periphery, which likely represent primary entry sites, while additional, smaller nucleation events may remain unresolved due to limited sensitivity and out-of-plane localization. Although the exact number of CPP-additive molecules required to induce water pore formation under physiological conditions is unknown, and individual pores cannot be resolved at the single-molecule level in spinning disk confocal time-lapse experiments, the qualitative observation of qCPP **4** influx from nucleation zones and intracellular activation to fCPP **3** together with quantitative image analysis identifies these zones as the dominant entry route. The immediate appearance of fCPP **3** signal inside cells within seconds after probe application strongly argues for direct translocation across the plasma membrane, most likely via transient water pore formation within nucleation zones, as no alternative pathway such as vesicular trafficking would be expected to proceed on this timescale.

#### Supplementary Fig. 4: Time lapse images of CPP-additive mediated delivery of fCPP in gramicidin pre-treated HeLa cells.

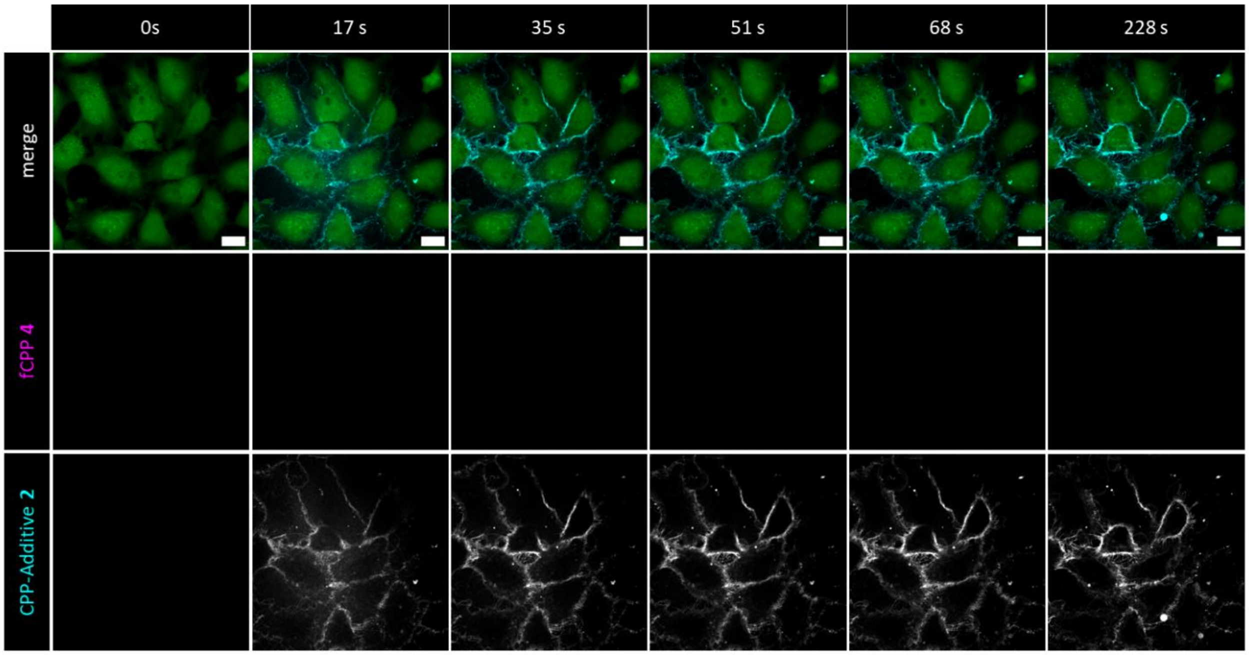

Composite time-lapse images image (green – cytosolic marker CMFDA, magenta – fCPP **3**, cyan – Cy5-CPP-additive **2**) of HeLa cells incubated for 5 min with 1.2 µg/mL gramicidin before addition of qCPP **4** and Cy5-CPP-additive **2** to a final concentration of 2.5 µM each. Time lapse images were recorded ever 8.5 s. CPP-additive shows cell surface accumulation but no fCPP signal. Therefore, we conclude that gramicidin blocks CPP uptake. CPP-additive **2** still associates with the cell surface but does not show formation of nucleation zones. Scale bar = 20 µm.

#### Supplementary Fig. 5: Pore formation in relation to transmembrane potential.

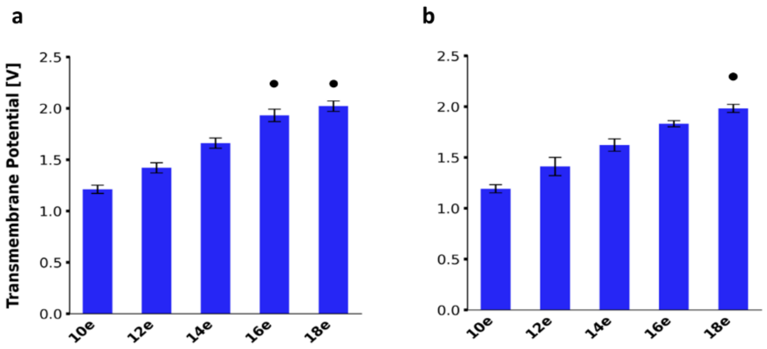

Transmembrane potentials derived from 100 ns simulations of pre-screened systems, calculated as 10 ns time-averaged values for a) linear $R_{10}$ and b) Membrane only. Simulation runs that resulted in pore formation are indicated by black dot. Error bars denote the standard deviation.

#### Supplementary Fig. 6: Pore width over time.

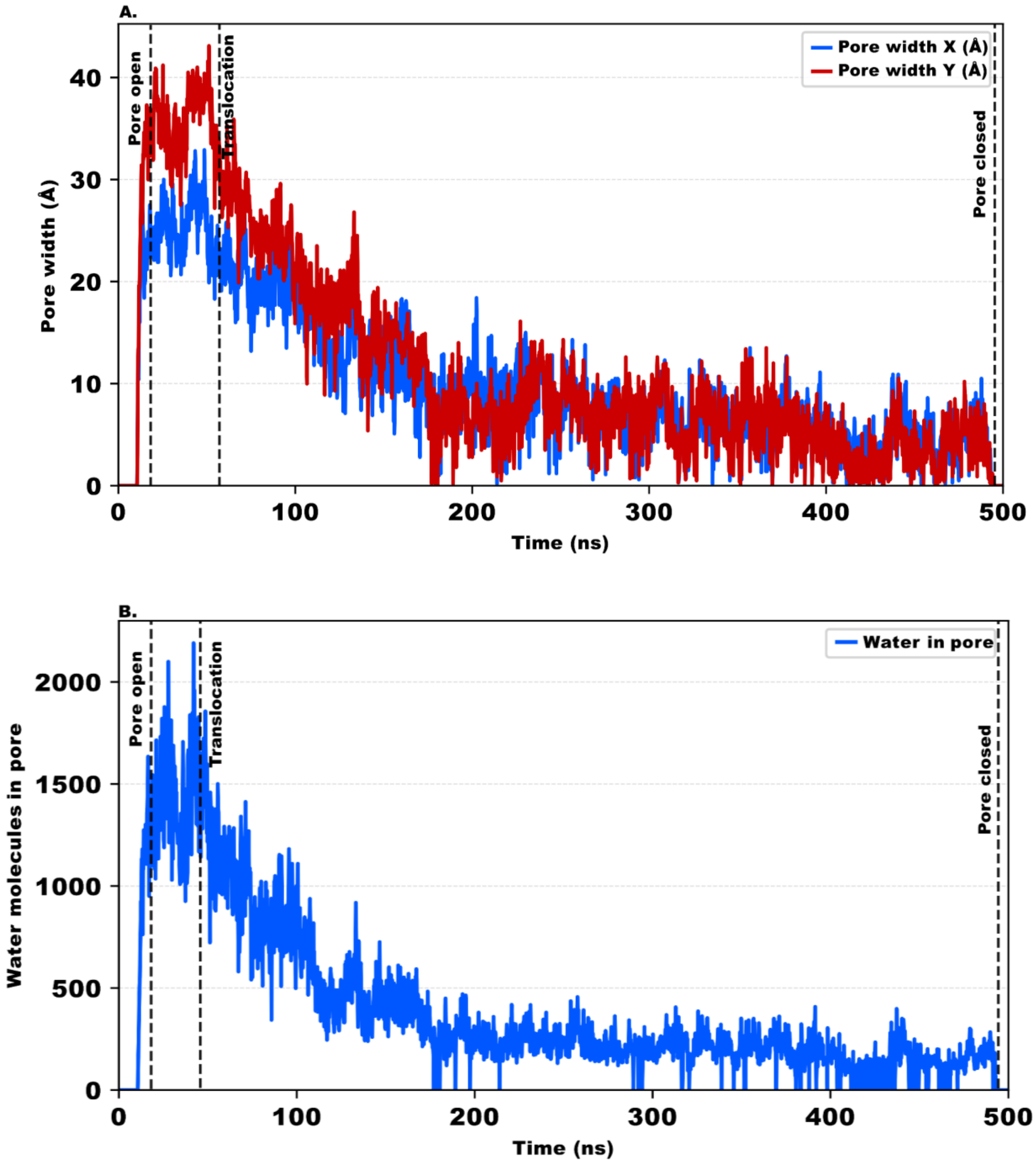

a) Time evolution of the minimum pore width along the X and Y directions during $R_{10}$ translocation. b) Number of water molecules inside the pore as a function of time. Vertical dashed lines indicate the beginning of pore opening, peptide translocation, and pore closure.

#### Supplementary Fig. 7: Strength of peptide-membrane interaction.

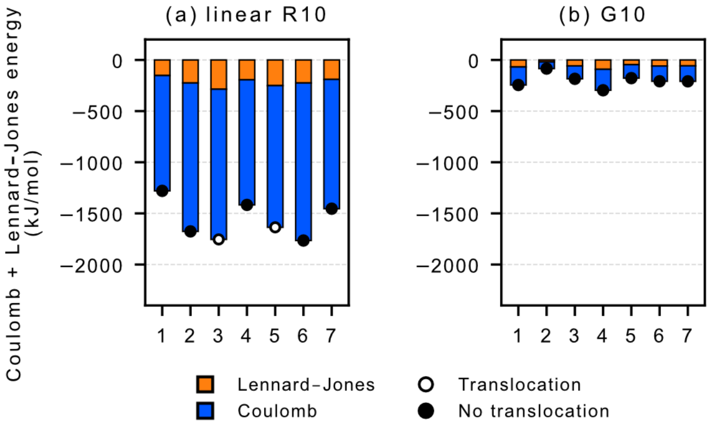

Mean peptide-membrane interaction energies decomposed into Lennard-Jones (orange) and Coulombic (blue) contributions for individual simulation replicas of linear $R_{10}$ and $G_{10}$. Stacked bars represent the total interaction energy. Open circles indicate replicas in which peptide translocation occurred, whereas filled circles denote non-translocating runs.

**Supplementary Fig. 8: Electrostatic potential profiles illustrating the reduction in transmembrane potential following peptide translocation.**

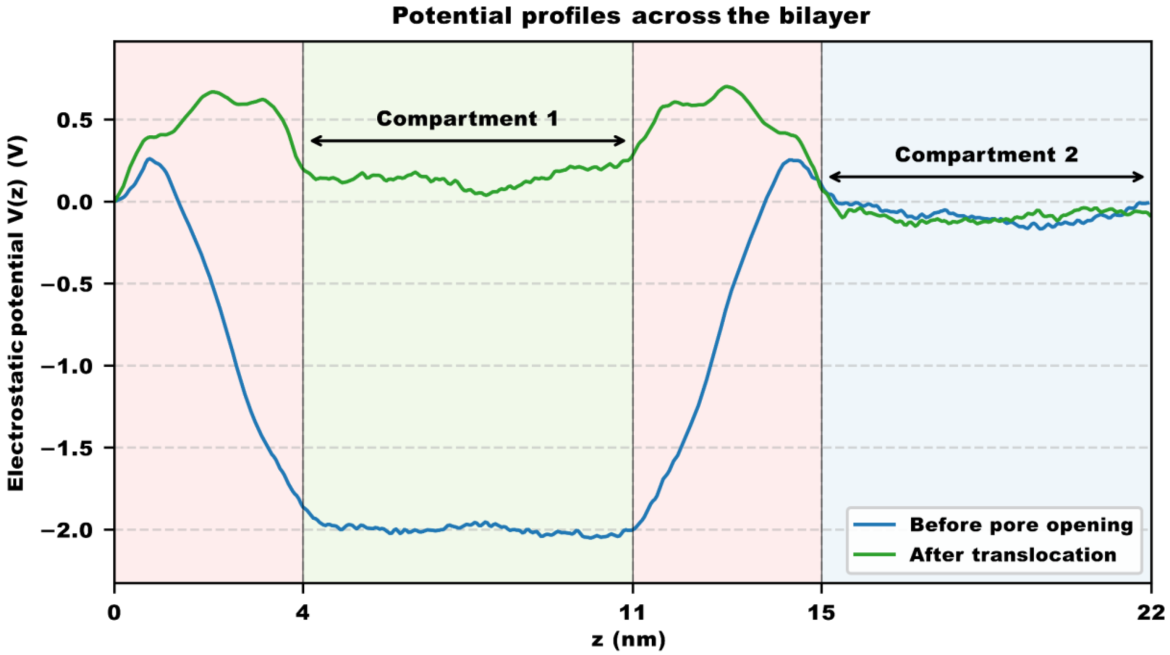

The electrostatic potential, V(z), was calculated along the z-axis across the double-membrane system for simulation segments before pore opening (blue) and after CPP translocation (green). The two membrane slabs are shown in light red, Compartment 1 in light green, and Compartment 2 in light blue.

#### Supplementary Fig. 9: Evaluation of further polyarginine derivatives.

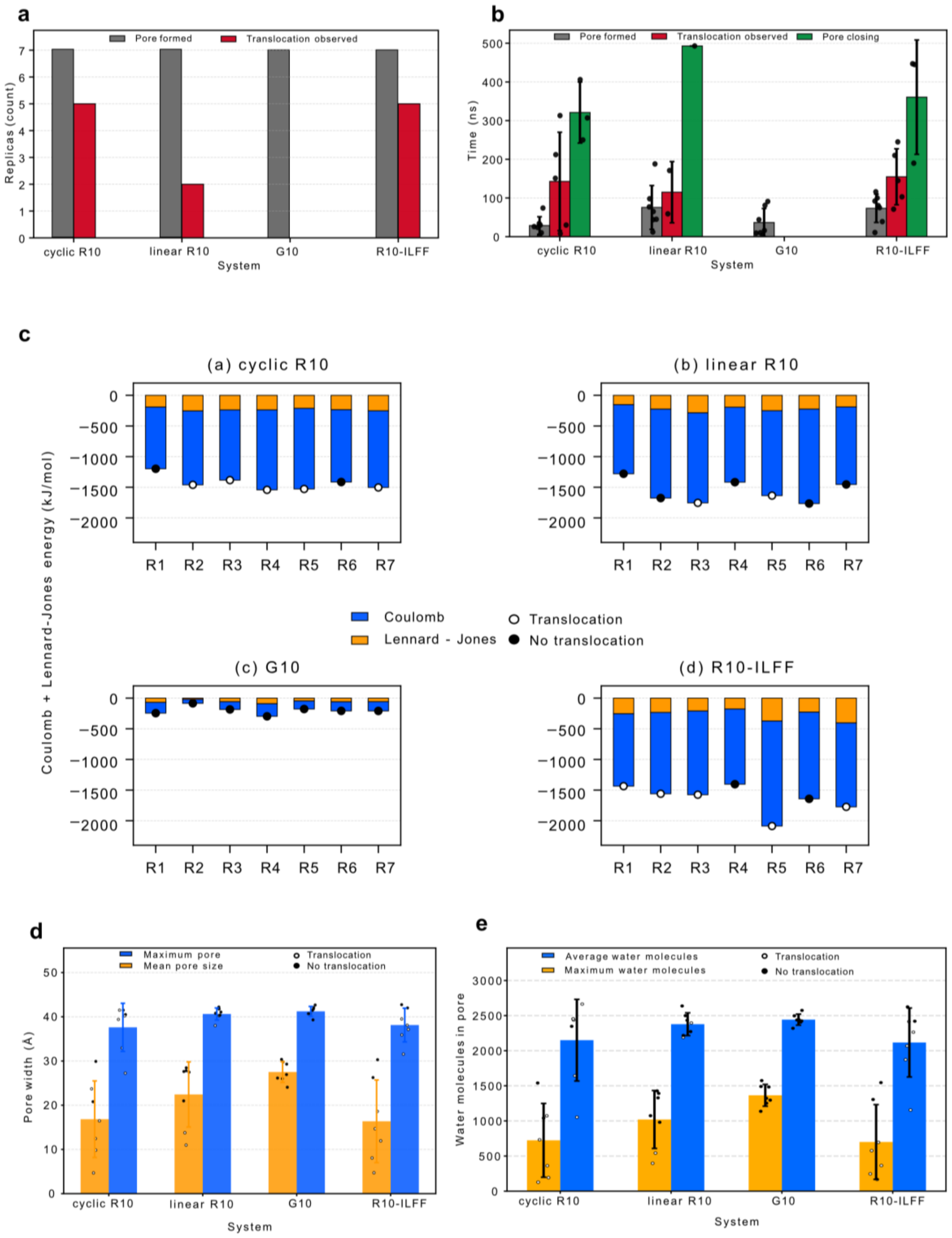

Discussion:

In addition to linear polyarginine ($R_{10}$), we modeled $R_{10}$-ILFF, a hydrophobic but non–membrane-anchored polyarginine, and cyclic $R_{10}$ ($cR_{10}$), which has been used for delivery of peptides and proteins in our previous studies. Despite their structural differences, $R_{10}$-ILFF and $cR_{10}$ exhibit similar behaviour during simulations comparable to linear $R_{10}$ and serve as additional examples of cationic peptide cargos.

a) Summary of pore formation and CPP translocation events observed across seven independent simulation runs of 500 ns. b) Mean times for pore formation, CPP translocation, and pore closure during simulations, shown as bar plots. Individual data points correspond to single simulation replicas, and error bars denote the standard deviation. c) Mean peptide–membrane interaction energies decomposed into Lennard–Jones (orange) and Coulombic (blue) contributions for individual simulation replicas (R1-7) of cyclic $R_{10}$, linear $R_{10}$, $G_{10}$, and $R_{10}$-ILFF. Stacked bars represent the total interaction energy. Open circles indicate replicas in which peptide translocation occurred, whereas filled circles denote non-translocating runs. d) Minimum and maximal pore radii in Å and e) mean and maximal water occupancy, quantified as the number of water molecules within the pore per nanosecond, derived from seven independent 500 ns simulation runs. Simulations runs in which peptide translocation occurred are indicated by black circles, whereas runs without translocation are shown as white cycles. Bars denote averages across the seven simulations, with individual data point shown together with the standard deviation.

#### Supplementary Fig. 10: Uptake of fluid marker in the absence of CPP-additive.

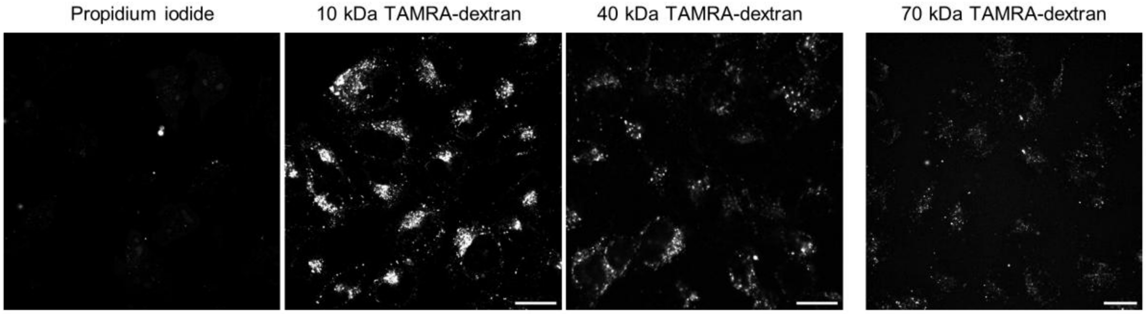

Confocal images (RFP channel) of HeLa cell that were incubated with 50 µg/mL propidium iodide, 400 µg/mL 10 kDa, 40 kDa, or 70 kDa TAMRA-dextran in serum-free FluoroBrite DMEM for 15 min. 70 kDa TAMRA-dextran was aquired at the same microscopy settings but on a different day. Scale bar = 20 µm.

#### Supplementary Fig. 11: Concentration dependent CPP-mediated uptake of fluid marker.

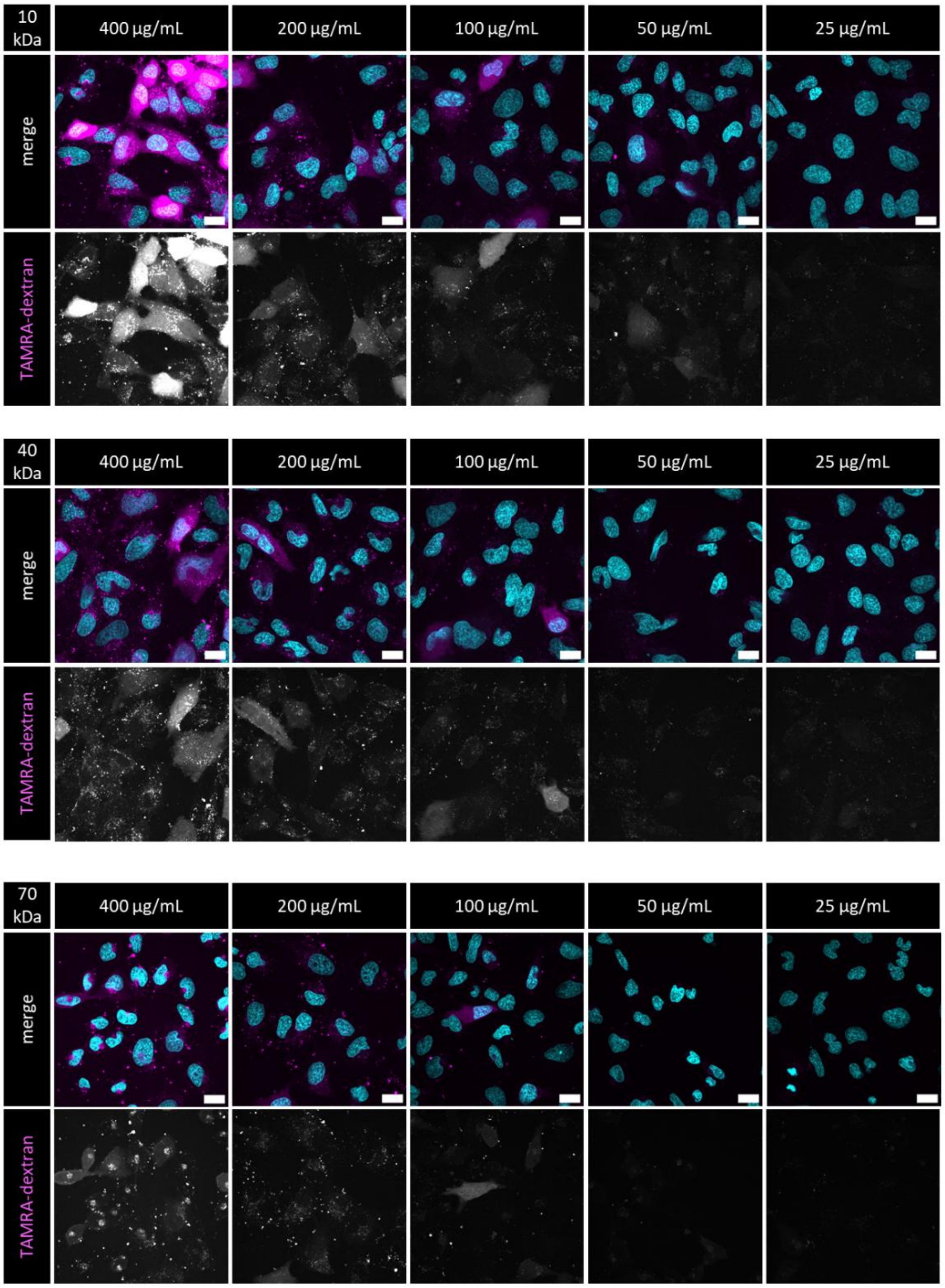

HeLa cells were incubated with the indicated concentration of TAMRA labeled dextran (top panel: 10 kDa; bottom panel: 40 kDa) as fluid marker in the presence of 5 µM CPP-additive **1** for 15 min. Nuclei were counterstained using Hoechst (cyan). Cells were imaged with a confocal spinning disc microscope under the same lase settings. Contrast settings are aligned for all panels.

#### Supplementary Fig. 12: Time lapse microscopy experiment monitoring cellular entry of GBP1 B.

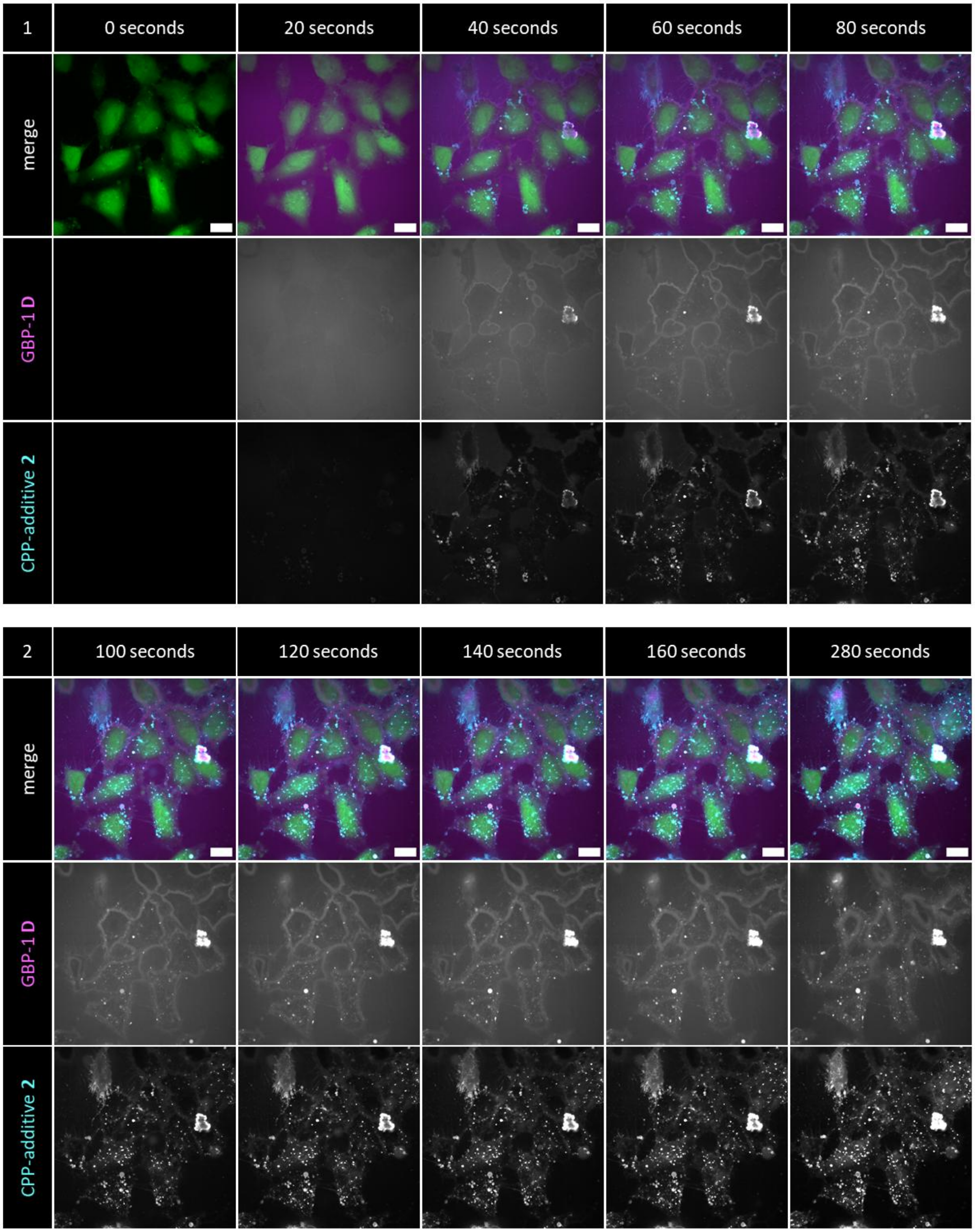

Composite images (green – cytosolic marker CMFDA, magenta – GBP1 **B**, cyan – Cy5-CPP-additive **2**) of HeLa cells treated with 2.5 µM GBP1 **B** and 2.5 µM of Cy5-CPP-additive **2** in serum-free FB DMEM. Scale bar = 20 µm.

#### Supplementary Fig. 13: Time resolved incubation of NLS-mCherry.

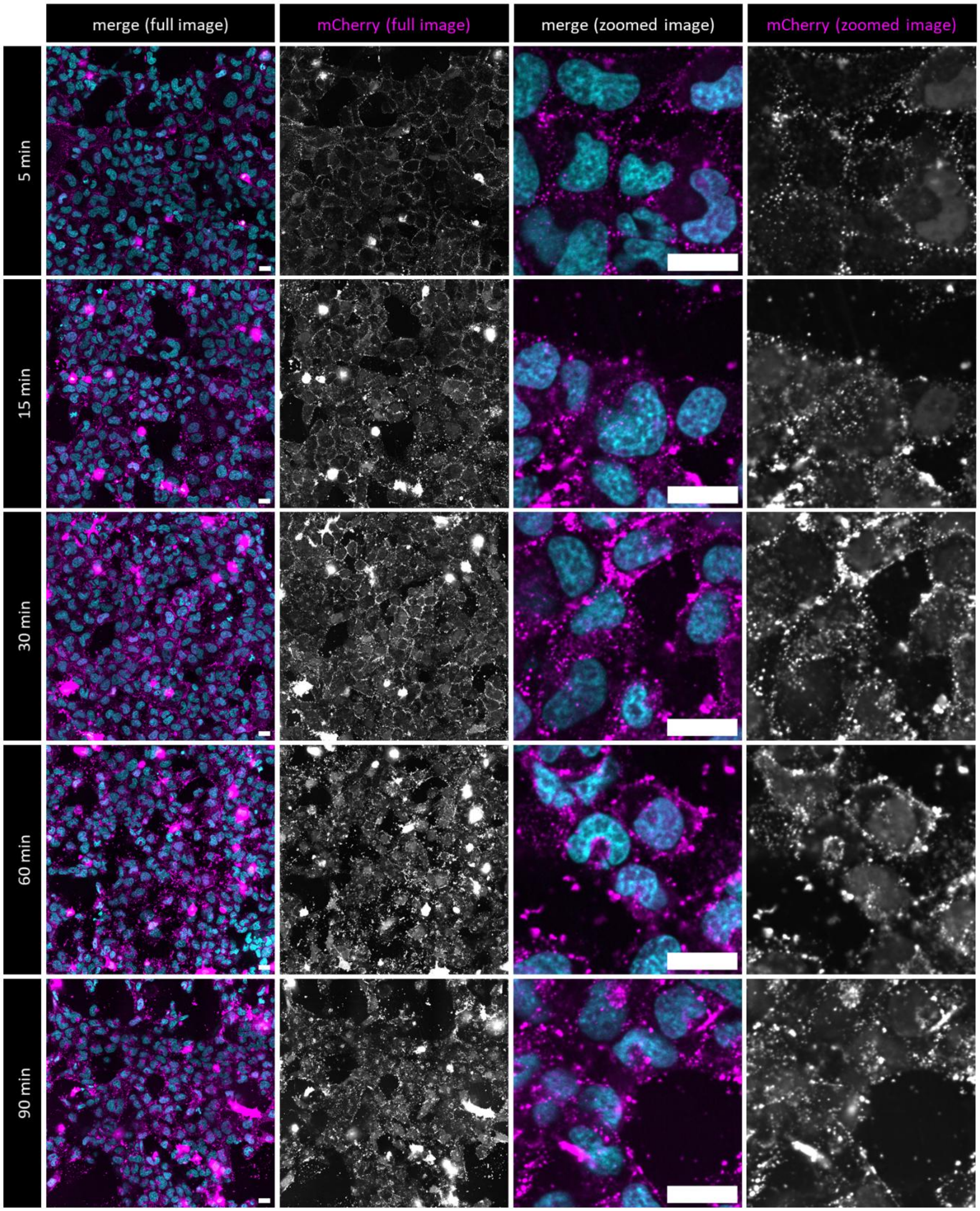

Hela cells treated with 2.5 µM NLS-mCherry **G** in the presence of 5 µM CPP-Additive **1** for the indicated time in serum free FluoroBrite DMEM medium. Pictures are representative regions of images acquired during automated acquisition and are basis for the quantification for nuclear fluorescence as described in the methods. Nuclei counterstained with Hoechst (cyan), Scale bars = 20 µM.

#### Supplementary Fig. 14: Co-delivery of GBP1 at high bystander concentrations.

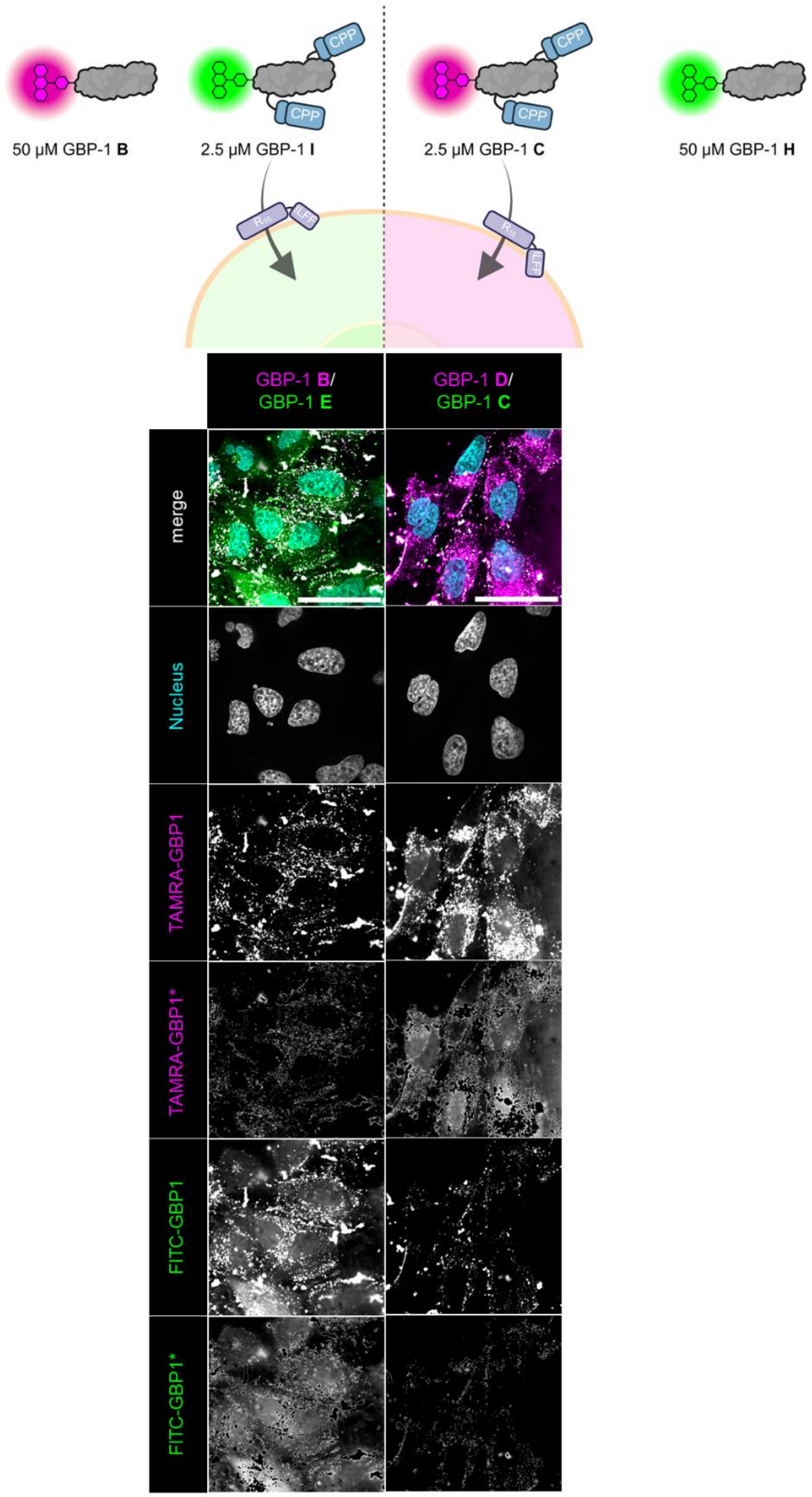

Co-delivery of 2.5 µM fluorescently labeled, CPP-modified GBP1 **C**/**I** in the presence of 50 µM GBP1 **B**/**H**. (green – GBP1 **H**/**I**, magenta – GBP1 **B**/**C**, cyan – nucleus, counterstained using Hoechst). *endosome exclusion was performed by Fiji macro using thresholding of intense signals.

#### Supplementary Fig. 15: Full SDS-PAGE gel of labeled, cell permeable GBP1 nanobodies.

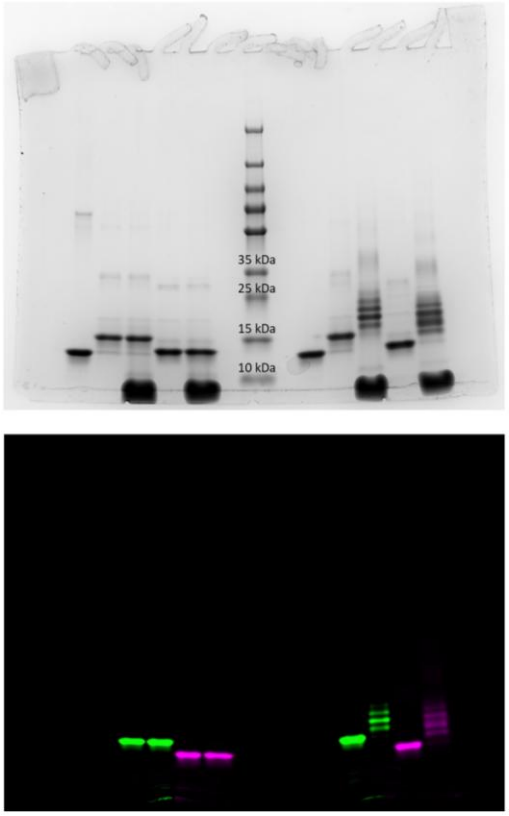

SDS-PAGE analysis of GBP1 proteins generated. Reducing samples (left) include β-mercaptoethanol causing the reductive cleavage. CPP-modifications from the protein. Coomassie stained gel (top) and fluorescent gel (bottom).

#### Supplementary Video 1

Time-lapse live cell microscopy of HeLa cells upon treatment with Cy5-CPP-Additives **2** (2.5 μM) and quenched qCPP **4** (2.5 μM) at 37 °C under 5% $CO_2$. Upon cytosolic entry, qCPP is reduced to fluorescent fCPP **3**. Cytosol is counterstained with cytosolic marker CMFDA.

#### Supplementary Video 2

Pore formation and translocation of linear polyarginine $R_{10}$. Representative atomistic MD simulation over 500 ns using CompEL set up with ion imbalance of 18 ions. POPC lipid headgroups in red (extracellular) dark blue (intracellular); water molecules in light blue.

#### Supplementary Video 3

Time-lapse live cell microscopy of HeLa cells upon treatment with Cy5-CPP-Additives **2** (2.5 μM) and quenched GBP1 **E** (2.5 μM) at 37 °C under 5% $CO_2$. Upon cytosolic entry, GBP1 **E** is reduced to fluorescent GBP1. Cytosol is counterstained with cytosolic marker CMFDA.

### General Information

#### Chemicals, solvents, and materials

Chemicals and solvents were purchased from Sigma-Aldrich (Merck Group, Germany), TCI (Tokyo chemical industry CO., LTD., Japan) and Acros Organics (Thermo Fisher scientific, USA), BLD Pharm (BLD Pharmatech, China) and used without further purification. Dry solvents were purchased from Acros Organics (Thermo Fisher scientific, USA). Aminoacids, coupling reagents and resins for SPPS were purchased from Novabiochem (Merck, USA) or Iris Biotech GmbH (Germany). Amino acids, rink amide resin and coupling reagents were purchased from Iris Biotech (Germany). HATU was purchased from Bachem (Switzerland). DIEA, TFA, Salts, LB medium, antibiotics and other buffer components were purchased from Carl Roth (Germany). Mammalian cell culture media and fetal bovine serum were purchased from BioWest and Gibco, respectively. Zeba™ Spin Desalting columns were purchased from Thermo Scientific (USA).

#### Flash and thin layer chromatography

Flash column chromatography was performed, using NORMASIL 60® silica gel 40-63 μm (VWR international, USA). Analytical thin layer chromatography (TLC) was performed on aluminum foil pre-coated with $SiO_2$-60 F254 (Macherey-Nagel, DE). Spots were visualized by fluorescence depletion with a 254nm lamp or manganese staining (10 g $K_2CO_3$, 1.5 g $KMnO_4$, 0.1 g NaOH in 100 mL $H_2O$), followed by heating.

#### Semi-preparative high-performance liquid chromatography (HPLC)

Semi-preparative HPLC was performed on a Shimadzu prominence HPLC system (Shimadzu Corp., Japan) with a CBM20A communication bus module, an FRC-10A fraction collector, 2 pumps LC-20AP, and a SPD-20A UV/VIS detector, using a VP250/21 Macherey-Nagel Nucleodur C18 HTec Spum column (Macherey-Nagel GmbH & Co. Kg, Germany).

#### NMR Spectroscopy

NMR spectra were recorded with a Bruker Avance III 600 MHz spectrometer (Bruker Corp., USA) at ambient temperature. Chemical shifts δ are reported in ppm relative to residual solvent peak ($CDCl_3$: 7.26 [ppm]; DMSO-$d_6$: 2.50 [ppm]; 4.79 $D_2O$ [ppm] for $^1$H-spectra and $CDCl_3$: 77.16 [ppm]; DMSO-$d_6$: 39.52 [ppm]; for $^{13}$C-spectra. Coupling constants *J* are stated in Hz. Signal multiplicities are abbreviated as follows: s: singlet; d: doublet; t: triplet; q: quartet; m: multiplet.

#### UPLC-UV/MS UPLC-UV/MS

Samples were recorded on a Waters H-class instrument equipped with a quaternary solvent manager, a Waters autosampler, a Waters TUV detector and a Waters Acquity QDa detector with an Acquity UPLC BEH C18 1.7 μm, 2.1 x 50 mm RP column with a flow rate of 0.6 mL/min (Waters Corp., USA). The following gradient was used: 0.1 % TFA in $H_2O$; B: 0.1% TFA in MeCN. 5 % B 0 - 0.5 min, 5-95 % B 0.5-3 min, 95 % B 3-3.9 min, 5 % B 3.9-5 min.

#### High resolution mass spectroscopy (HR-MS)

High resolution ESI-MS spectra were recorded on a Waters H-class instrument equipped with a quaternary solvent manager, a Waters sample manager-FTN, a Waters PDA detector and a Waters column manager with an Acquity UPLC protein BEH C18 column (1.7 μm, 2.1 mm x 50 mm). Samples were eluted with a flow rate of 0.3 mL/min. The following gradient was used: "QTof": 0.01% FA in $H_2O$; B: 0.01% FA in MeCN. 5 % B: 0-1 min; 5 to 95 % B: 1-7min; 95 % B: 7 to 8.5 min. Mass analysis was conducted with a Waters XEVO G2-XS QTof analyzer. (A significant portion of small molecules could not be detected by QTof HR-MS due to their inherent instability upon ionization.)

#### BioRAM conjugation to aliphatic amines

100 µM of protein was incubated with respective peptide (**13:** BioRAM -$R_8$ or **14:** BioRAM-$R_4$) at 1200−2000 µM (12-20 eq.) in HEPES buffer (200 mM HEPES at pH = 8.25, 140 mM NaCl). Test reactions were performed in 20 µL, upscale reactions were done using a total volume of 850 µL for NLS-mCherry and 50 µL for GBP1. The reaction mixture was incubated at 20 °C for 3 hours at 500 rpm shaking. The resulting modified protein was separated from excess peptide by using an Amicon® spin filtration (Amicon® Ultra-0.5, 3-10k MWCO) until a 1:100 dilution, followed by desalting by Zeba™ Spin purification to remove remaining small molecules and potential aggregates. Proteins were rebuffered within the Zeba™ Spin purification step in storage buffer (5 mM HEPES at pH = 7.5, 140 mM NaCl, 2.5 mM KCl, 5 mM Glycine). Protein concentration was determined using a Nanodrop (Nanodrop One/OneC, Thermo Scientific, USA). The samples were frozen in liquid nitrogen and stored at -80 °C for up to 3 months.

#### SDS-PAGE gel analysis

SDS-PAGE gels were self-casted using 12 or 15 % acrylamide and run in 1x Tris-Glycine-SDS buffer (Carl Roth, Germany) at 180 V. Samples were diluted in water or PBS and supplemented by 1x Laemmli buffer (Bio-Rad, USA) with (reducing, '+') or without (non-reducing, '-') the addition of β-mercaptoethanol (ACROS Organics) and heated for 10 min at 95 °C. The proteins were stained with Blue stain (FastGene) for at least 2 h and destained with water. The images were acquired using a ChemiDoc™ Imager (Bio-Rad, USA).

#### Cell Culture

All cell incubation and treatments were done in a humidified incubator at 37 °C with 5 % $CO_2$. Cells were split at 70%–90% confluency and used in passages 4-15. HeLa cells were obtained from ATCC (ATCC, catalogue number: CCL-2). U2OS were kindly donated by Dorothea Fiedler. HeLa, U2OS were cultivated in Dulbecco's Modified Eagle Medium (DMEM) high glucose (HG) medium with stable L-glutamine, 10% fetal calf serum (FCS). A549 cells were obtained from (DSMZ, catalogue number: ACC 107). A549 were maintained in DMEM/Ham's F-12, 2 mM stable L-glutamine and 10% FCS. SJSA1 were generously provided by Andreas L. Marzinzik (Novartis). MiaPaCa-2 were obtained from ATCC (ATCC catalog number: CRL1420) . MCF-7 were obtained from DSMZ (ACC 115, LOT23). SJSA1, MiaPaCa-2 and MCF-7 cells were maintained in Roswell Park Memorial Institute (RPMI) 1640 medium with 10% FCS. All cells were passaged every 48 h or at 80% confluency.

#### Sortase mediated ligation

General procedure for sortase mediated ligation is as follows: 1 eq N-terminal glycine containing protein was labeled using 0.3 eq eSrtA 5M-H6 and 10 eq sortase peptide in 20 mM Hepes, 1 mM $CaCl_2$, pH 7.4. The reaction was left to proceed at 4° C, shaking at 450 rpm for 60 minutes. Pre-equilibrated Ni-NTA beads with 20 mM Hepes, 150 mM NaCl, pH 7.4 were added to the reaction and left for 10−12 minutes to bind eSrtA 5M-H6 and reaction intermediates. The beads were washed again with specified equilibration buffer and the flow through and washes were collected and combined in a 3 kDa MWCO amicon and concentrated to the desired working concentration. The remaining product was desalted with 0.5 mL Zeba™ Spin (ThermoFisher) to remove excess peptide.

#### Statistical treatment

All biological experiments were performed in three biological replicates unless otherwise specified. The statistical tests and values of *N* are reported in the figure legends where appropriate. Graphs and statistical significances were determined using GraphPad Prism 10.

### Data Availability

All experimental data, materials and methods, analytical procedures, cell assays, copies of spectra are available in the Main Text and the Supplementary Information (ESI).

### Experimental procedures

#### Cytotoxicity experiment

Lactate dehydrogenase (LDH) and Mitochondrial dehydrogenase (WST-1) were tested in parallel to determine cytotoxicity of qCPP **4** and CPP-additive **1** treatment.

5000 HeLa cells per well were seeded in a 96-well plate (Sarstedt, F-bottom, Item No.: 83.3924) and incubated for 48 h at 37°C, 5 % $CO_2$ to settle. Cells were treated with qCPP **4** at 2.5 µM, 5 µM and 10 µM in the presence and absence of 5 µM CPP-additive **1** (TNB-$R_{10}$-ILFF) in 100 µL of FluoroBrite DMEM with 0% FCS at 37°C, 5% $CO_2$ for 1h.

##### Lactate dehydrogenase (LDH) release

Lactate dehydrogenase (LDH) release was measured using the CytoTox-ONE™ Homogeneous Membrane Integrity Assay from Promega (Cat. G7890). 3 min before the end of the 1 h - incubation time, 2 µL of lysis solution was added to the three control wells for maximum LDH release. The medium (100 µL) was transferred to a fresh 96 well plate and fresh DMEM-HG medium with 10% FCS was added to the cells, which were transferred back to the incubator for parallel WST-1 assessment after 24 hours. The supernatant was equilibrated for 10 min at 22 °C and background fluorescence (ex.: 560 nm; em.: 590 nm) of the fCPP was recorded using a M200Pro plate reader (TECAN, Switzerland). LDH release was tested by addition of 100 µL of prepared reagent (substrate + assay buffer). The reaction mixture was incubated for 10 min at 22 °C in the dark. 50 µL of Stop solution was added to each well and fluorescence (ex.: 560 nm; em.: 590 nm) was recorded after 10s of shaking using the plate reader. Background fluorescence was subtracted at the respective concentration of peptide. Data was normalized to medium treated cells (0% release) and maximum LDH release (100% release).

##### WST-1 assay

For cell viability measurements using Mitochondrial dehydrogenase (WST-1) assay cells were incubated for 24h after compound treatment and exchange of medium. Then, 10 µL of WST-1 reagent was added to each well and incubated for 60 min at 37 °C, 5% $CO_2$. Absorbance at 460 nm was measured using a M200Pro plate reader (TECAN, Switzerland). Data was normalized to medium treated cells (100 % viability) and 0.1 % Triton-X100 (0 % viability).

#### Cellular uptake

HeLa cells were seeded in a black-walled 96-well glass-bottom plate with flat transparent bottom (Screenstar, microplate, F-bottom, Item No.: 655866) at 10'000 cells/well in growth medium and allowed to attach for 48 h. Cells were washed with PBS once in case of serum-free incubation. 5 µM TNB-$R_{10}$-ILFF were mixed with the indicated amount of NLS-mCherry in 100 µL growth medium with 0 or 10 % FCS for 1 hour at 37 °C, 5% $CO_2$. Treated cells were washed with prewarmed 200 µL PBS containing 0.5 mg/mL heparin. The nucleus is counterstained with Hoechst 33342 at (1:1000) in FluroBrite DMEM + 10% FCS for 5 min. The media was replaced with fresh FluoroBrite DMEM + 10% FCS followed by confocal microscopy.

#### Microscopy

Confocal microscopy was done using a Nikon-CSU spinning disk microscope with a CSU-X1 (Yokogawa with modifications by Andor) confocal scanner unit and a live cell imaging chamber with temperature and $CO_2$ control (OKOlab). Images were captured using either a PlanApo 60x NA 1.4 oil objective (Nikon) or a PlanApo 40x NA 1.4 air objective (Nikon) and an EMCCD (AU888, Andor). The automated quantification was performed using a 40x air NA 0.95. Brightfield images were acquired along with fluorescence images.

Standard laser, a quad Dicroic (400-410,486-491, 560-570, 633-647, AHF) and Emission filters were used in the acquisition of confocal fluorescence images [BFP (Hoechst 33342), ex.: 405 nm em.:450/50; GFP (Atto488, mVenus), ex.: 488 em.:525/50; RFP (mCherry), ex.: 561 em.:600/50 nm.]

### Time lapse microscopy

HeLa cells were seeded at 1 × $10^5$ cells per well in an µ-Slide 8 well$^{high}$ Glass bottom chamber (ibidi, item no. 80807-90) in 300 µL of medium and allowed to attach for 48 h. Medium was aspirated and cells were incubated with 150 µL FluoroBrite DMEM, 0% FCS and 5 µM of CellTracker™ CMFDA stain (Thermo Fisher, item no. C7025) for 30 min. Cells were washed once with PBS and the medium was exchanged to 150 µL FluoroBrite DMEM 0% serum before the slide was mounted to the microscope. After focus adjustment on the microscope using the GFP channel (CMFDA signal), automated time-lapse acquisition was initiated. Subsequently, 150 µL FluoroBrite DMEM containing 5 µM qCPP **4** (final concentration 2.5 µM), and 5 µM Cy5-CPP-additive **1** (final concentration 2.5 µM) were added.

Images were acquired every 5s−10s for GFP-, RFP-, and iRFP-channel using a Plan Apo 40×/1.4 NA air objective (Nikon) over a total imaging period of 5 min.

##### Colocalization analysis during time lapse microscopy

We filtered for areas of intense fluorescence of Cy5-CPP-additive **2** (iRFP channel), transforming them into a mask and measuring fCPP **3** fluorescence (RFP channel) inside and outside these areas over time. We used top hat filter and the dilate function to expand the foreground regions in the binary mask and thereby slightly enlarge selected zones. We displayed the quotient over time to describe the co-occurrence of nucleation zones (CPP-additive signal) and activated fCPP (entry points of fCPP).

$$f(t) = \frac{I_{RFP}(inside)}{I_{RFP}(outside)}$$

Please refer to section "Microscopy Macros", "Time-lapse microscopy to estimate co-localization" for FIJI-macro.

### DiBAC$_4$(3) membrane potential experiments

<u>Time lapse microscopy</u>

HeLa cells were seeded at 1 × $10^5$ cells per well in an µ-Slide 8 well$^{high}$ Glass bottom chamber (ibidi, item no. 80807-90) in 300 µL of medium and allowed to attach for 48 h. The medium was aspirated, and cells were incubated with 500 nM DIBAC dye (Thermo Scientific) in 150 µL FluoroBrite DMEM (10% FCS) for 30 min. The incubation solution was then replaced with 200 µL FluoroBrite-DMEM (0% FCS) containing 500 nM DiBAC$_4$(3).

After focus adjustment on the microscope using the GFP channel (DIBAC signal), automated time-lapse acquisition was initiated. Subsequently, 100 µL FluoroBrite DMEM (0% FCS) containing 500 nM DiBAC$_4$(3), 7.5 µM qCPP **4** (final concentration 2.5 µM), and 15 µM CPP-additive **3** (final concentration 5 µM) were added. For the DiBAC$_4$(3) control, 100 µL FluoroBrite-DMEM (0% FCS) containing 500 nM DIBAC was added.

For gramicidin-pretreated samples, the DiBAC$_4$(3) incubation solution was exchanged for 150 µL FluoroBrite DMEM (0% FCS) containing 500 nM DiBAC$_4$(3). Prior to time-lapse imaging, 50 µL of 4× gramicidin solution in FluoroBrite DMEM (0% FCS, 500 nM DiBAC$_4$(3)) was added to the wells to reach a final concentration of 1.2 µg/mL and incubated for 5 min. Imaging was then started, followed by the addition of 100 µL FluoroBrite DMEM (0% FCS) containing 500 nM DiBAC$_4$(3), 7.5 µM qCPP **4** (final concentration 2.5 µM), and 15 µM CPP-additive **1** (final concentration 5 µM).

Images were acquired every 15 s at three positions for GFP- and RFP-channel using a Plan Apo 40×/1.4 NA air objective (Nikon) over a total imaging period of 7 min.

Note: Bleaching of the $DiBAC_4(3)$ has been observed over time (decline of signal intensity of control sample Fig. 3b). We have not corrected samples for bleaching for the data displayed.

Flow cytometry-based membrane potential quantification. Measurement of resting membrane potential using $DiBAC_4(3)$ was performed as described previously.[1] Briefly, cells were seeded into 10-cm culture dishes for 48 h or until 80% confluency. The cells were detached with Accutase, centrifuged (800 × *g,* 5 min, 28 °C), and resuspended in their respective cell culture media indicated in the 'Cell Culture' section to a concentration of 4 × $10^5$ cell/mL. The cell suspension was aliquoted at 500 µL in each prepared 1.5 microcentrifuge tubes (2 × $10^5$ cell/tube). For the standard curve (6 tubes), cells were depolarized by adding 1.2 µg/mL gramicidin and incubating the tubes on a heated benchtop shaker (37 °C, 800 rpm) for 5 min. $DiBAC_4(3)$ was directly added to individual tube to a final concentration of 400, 200, 100, 50, 25, 0 nM and further incubated for 20 min. For measuring resting membrane potential, $DiBAC_4(3)$ was directly added to the tube to a final concentration of 200 nM. For cells pre-treated with TRAM-34 and SKA-31, detached cells were treated with respective 10 µM of respective small molecule for 5 min prior to the addition of $DiBAC_4(3)$ to a final concentration of 200 nM and 20 min incubation. For measuring the membrane potential at different KCl concentrations, detached cells in microcentrifuge tubes were centrifuged (800 × *g,* 5 min, 28 °C) and resuspended in HEPES buffer with indicated KCl concentration and incubated on a heated benchtop shaker (see above) for 30 min prior to the addition of $DiBAC_4(3)$ (final concentration of 200 nM) and incubation for 20 min. Mean fluorescence intensity of DiBAC4(3) was measured with 10,000 gated events using an LSRFortessa (BD Biosciences).

Calculation of membrane potential was performed as follows.[1] The intracellular $DiBAC_4(3)$ concentration ($D_i$) for each setup was determined using the standard curve of depolarized cells (*x*-axis: $DiBAC_4(3)$, *y*-axis: mean fluorescence intensity), where $D_i$ was assumed to be equal to the initial extracellular dye ($D_e$). The membrane potential (mV) was then calculated using Nernst equation for the anionic $DiBAC_4(3)$ dye at 37°C below.

$$\mathrm{mV} = -60\ log_{10}\left(\frac{D_e}{D_i}\right)$$

#### Fluid marker uptake

HeLa cells (2.5 × $10^4$ cells/well) were seeded onto 98-well glass bottom plate (Cellvis). After 48 h of incubation, the cells were washed twice with pre-warmed PBS, followed by treatment with 50 µg/mL propidium iodide, 400 µg/mL neutral 10 kDa tetramethylrhodamine(TMR) dextran, neutral 40 kDa tetramethylrhodamine(TMR) dextran, or 70 kDa tetramethylrhodamine(TMR) dextran (Thermo Scientific) with or without 5 µM CPP-additive **1** in serum-free FluoroBrite DMEM for 15 min. Treated cells were washed with heparin-PBS and counterstained with counterstained with Hoechst 33342 at (1:1000) in FluroBrite DMEM + 10% FCS for 5 min. Imaging was performed according the 'Microscopy' section above. For cells treated with 1.2 µg/mL gramicidin, 20 µM TRAM34, or 20 µM SKA-31, cells were pre-incubated with the indicated small molecule reagents in serum-free FluoroBrite DMEM for 5 min prior to fluid marker incubation in the presence of the small molecule (except gramicidin). For Figure 3, cells with diffused cytosolic signal (%)" corresponds to the number of cells exhibiting diffused cytosolic signal (see below) divided by the total number of counted nucleus. Bar graphs shows the mean value ± standard error of three biological replicates. Data presented as mean ± SE of three biological replicates. *P*-values were determined by one-way analysis (ANOVA) followed by Dunnett's post hoc test.

#### NLS-mCherry-BioRAM uptake

Cells (2.5 × $10^4$ cells/well) were seeded onto 96-well glass bottom plate (Cellvis). After 48 h of incubation, the cells were washed twice with pre-warmed PBS, followed by treatment with 2.5 µM NLS-mCherry-BioRAM **G** with 5 µM CPP-additive **1** serum-free FluoroBrite DMEM for 15 min. Treated cells were washed with heparin-PBS and counterstained with Hoechst 33342 at (1:1000) in FluroBrite DMEM + 10% FCS for 5 min prior to confocal imaging. Quantification of nuclear fluorescence (Median fluorescence intensity) was performed according to previously published procedures.[2] For time-course uptake experiment, prepared cells were treated with NLS-mCherry-BioRAM **G** and CPP-additive **1** for 5, 15, 30, 60, 90 min. Succeeding steps were similar as above. For small molecule pre-treatment, prepared cells were incubated with DMSO (control), 1.2 µg/mL gramicidin, 20 µM TRAM34, or 20 µM SKA-31 in serum-free FluoroBrite DMEM for 5 min. Protein treatment and following steps were similar as above. For uptake in different KCl concentrations, protein treatment was performed in HEPES buffer (1 mM MgCl2, 1.8 mM CaCl2, 10 mM D-glucose, 20 mM HEPES at pH 7.4) having different concentrations of KCl (5.4, 10, 20, 40, 80, 140 mM) and NaCl (140, 135.4, 125.4, 105.4, 65.4, 5.4 mM) for 15 min. Succeeding procedures are the same as above.

### Co-delivery experiment

HeLa cells were seeded at 0.5 × $10^5$ cells/mL in 96-well plates (100 µL per well) and allowed to attach for 48 hours. Nanobodies were prepared as 5 µM stocks in FluoroBrite DMEM (0% FCS) containing 5 µM CPP-additive **1** and mixed with other Nanobody as indicated to a final concentration of 2.5 µM for each nanobody in solution. After a single wash with FluoroBrite DMEM (0% FCS), cells were incubated for 15 minutes with the respective nanobody mixture. Subsequently, cells were washed twice with heparin and stained with Hoechst in FluoroBrite DMEM (10% FCS) for 5 minutes. Finally, the medium was replaced with FluoroBrite DMEM (10% FCS), and cells were imaged.

### Computational simulations

Molecular dynamics (MD) simulations were performed using GROMACS 2021.7[3,4] with the CHARMM36m all-atom force field[5-7] to investigate the translocation mechanism of peptides. To this end, four distinct peptide-containing models were generated: (i) linear deca-arginine (linear $R_{10}$), (ii) cyclic deca-arginine (cyclic $R_{10}$), (iii) deca-glycine ($G_{10}$), and (iv) hydrophobic polyarginine $R_{10}$-ILFF. Initial three-dimensional peptide structures were generated using AlphaFold2[8,9]**,** and a symmetric bilayer composed of 256 1-palmitoyl-2-oleoyl-sn-glycero-3-phosphocholine (POPC) lipids was built using the CHARMM-GUI Membrane Builder.[10-12] The bilayer was solvated with a 20 Å layer of TIP3P water[13] and neutralized with 150 mM $K^+$/$Cl^-$. The peptide and membrane models were then combined, and the peptides were positioned approximately 5 Å from the membrane surface in the aqueous phase.

Following the model building, each single-bilayer system was energy minimized and thermally equilibrated applying the CHARMM-GUI protocol.[14-16] The equilibrated models were then duplicated along the membrane normal (z-axis) to generate a double-bilayer configuration, thereby creating two distinct aqueous compartments (C1 and C2) within a single periodic simulation box. To impose and maintain an ionic gradient mimicking a transmembrane potential, the subsequent production simulations were performed using the computational electrophysiology (CompEL) framework (Fig.1A), originally described by Kutzner *et al*.[17] . In our simulations, an ionic charge imbalance of 18e (where *e* is the elementary charge) was chosen between the two compartments preserving overall system electroneutrality. Specifically, C1 carried a net charge of +9e, whereas C2 carried a net charge of −9e. The details for each system are summarized in Table S1. This high charge imbalance corresponding to an applied potential of ca. 2 V was chosen based on an initial screening. In these preceding simulations, charge imbalances ranging from 10e and 18e were applied and the membrane stability was evaluated to determine the minimal potential required to induce electroporation within a 100 ns timescale (Figure S5).

Production simulations of 500 ns were conducted in the isothermal–isobaric (NPT) ensemble. The temperature of 303.15 K was kept constant by the velocity-rescale (V-rescale) thermostat[18] with a coupling time constant of 1.0 ps, applied separately to the lipid–peptide complex and the solvent. The pressure was maintained at 1 bar using the stochastic cell-rescale (C-rescale) barostat[19] under semi-isotropic conditions, with a coupling time constant of 5.0 ps and a compressibility of 4.5 × $10^{-5}$ $bar^{-1}$. A cutoff distance of 1.2 nm was applied for both electrostatic and van der Waals interactions. Long-range electrostatic interactions were treated with the Particle-Mesh Ewald (PME) method.[20] Periodic boundary conditions were imposed in all three spatial dimensions. All bonds involving hydrogen atoms were constrained using the LINCS algorithm[21], enabling a stable integration time step of 2 fs. To ensure statistical robustness and reproducibility, seven independent replicas were performed for each system, starting from identical coordinates but with different initial velocity assignments.

**Table S1.** List of simulation systems, summarizing the applied ionic imbalance, numbers of water molecules, ions, peptides, and lipids, as well as the size of the original simulation box.

| System | Ionic Imbalance | Peptides | POPC (per bilayer) | TIP3 Water Molecules | Ions ($Na^+$/ $Cl^-$) | Original box size (Lx × Ly × Lz, nm) |
|---|---|---|---|---|---|---|
| cyclic $R_{10}$ | 10e | 1 | 256 | 42946 | 25 / 35 | 9.08 × 9.08 × 23.42 |
| cyclic $R_{10}$ | 12e | 1 | 256 | 42946 | 24 / 34 | 9.08 × 9.08 × 23.42 |
| cyclic $R_{10}$ | 14e | 1 | 256 | 42946 | 23 / 33 | 9.08 × 9.08 × 23.42 |
| cyclic $R_{10}$ | 16e | 1 | 256 | 42946 | 22 / 32 | 9.08 × 9.08 × 23.42 |
| cyclic $R_{10}$ | 18e | 1 | 256 | 42946 | 21 / 31 | 9.08 × 9.08 × 23.42 |
| linear $R_{10}$ | 10e | 1 | 256 | 42904 | 25 / 35 | 9.09 × 9.09 × 23.30 |
| linear $R_{10}$ | 12e | 1 | 256 | 42904 | 24 / 34 | 9.09 × 9.09 × 23.30 |
| linear $R_{10}$ | 14e | 1 | 256 | 42904 | 23 / 33 | 9.09 × 9.09 × 23.30 |
| linear $R_{10}$ | 16e | 1 | 256 | 42904 | 22 / 32 | 9.09 × 9.09 × 23.30 |
| linear $R_{10}$ | 18e | 1 | 256 | 42904 | 21 / 31 | 9.09 × 9.09 × 23.30 |
| $G_{10}$ | 10e | 1 | 256 | 43270 | 35 / 35 | 9.12 × 9.12 × 23.30 |
| $G_{10}$ | 12e | 1 | 256 | 43270 | 34 / 34 | 9.12 × 9.12 × 23.30 |
| $G_{10}$ | 14e | 1 | 256 | 43270 | 33 / 33 | 9.12 × 9.12 × 23.30 |
| $G_{10}$ | 16e | 1 | 256 | 43270 | 32 / 32 | 9.12 × 9.12 × 23.30 |
| $G_{10}$ | 18e | 1 | 256 | 43270 | 31 / 31 | 9.12 × 9.12 × 23.30 |
| $R_{10}$ ILFF | 10e | 1 | 256 | 42780 | 25 / 35 | 9.07 × 9.07 × 23.44 |
| $R_{10}$ ILFF | 12e | 1 | 256 | 42780 | 24 / 34 | 9.07 × 9.07 × 23.44 |
| $R_{10}$ ILFF | 14e | 1 | 256 | 42780 | 23 / 33 | 9.07 × 9.07 × 23.44 |
| $R_{10}$ ILFF | 16e | 1 | 256 | 42780 | 22 / 32 | 9.07 × 9.07 × 23.44 |
| $R_{10}$ ILFF | 18e | 1 | 256 | 42780 | 21 / 31 | 9.07 × 9.07 × 23.44 |

**Table S2.** List of simulation systems, summarizing the applied ionic imbalance, initial transmembrane voltage, net peptide charge, number of ions in each compartment (comp.), simulation replicas, and simulation runs.

| System | Ionic imbalance | Replicas | Time (ns) | Net peptide charge | Comp. 1 ($Na^+$/$Cl^-$) | Comp. 2 ($Na^+$/$Cl^-$) | Initially applied voltage (ΔU) |
|---|---|---|---|---|---|---|---|

| | | | | | | | |
|---|---|---|---|---|---|---|---|
| cyclic $R_{10}$ | 10 e | 7 | 500 | 10 | 10 / 15 | 15 / 20 | 1.21 ± 0.04 |
| cyclic $R_{10}$ | 12 e | 7 | 500 | 10 | 10 / 14 | 14 / 20 | 1.42 ± 0.05 |
| cyclic $R_{10}$ | 14 e | 7 | 500 | 10 | 10 / 13 | 13 / 20 | 1.66 ± 0.05 |
| cyclic $R_{10}$ | 16 e | 7 | 500 | 10 | 10 / 12 | 12 / 20 | 1.93 ± 0.06 |
| cyclic $R_{10}$ | 18 e | 7 | 500 | 10 | 10 / 11 | 11 / 20 | 2.02 ± 0.05 |
| linear $R_{10}$ | 10 e | 7 | 500 | 10 | 10 / 15 | 15 / 20 | 1.19 ± 0.04 |
| linear $R_{10}$ | 12 e | 7 | 500 | 10 | 10 / 14 | 14 / 20 | 1.41 ± 0.09 |
| linear $R_{10}$ | 14 e | 7 | 500 | 10 | 10 / 13 | 13 / 20 | 1.62 ± 0.06 |
| linear $R_{10}$ | 16 e | 7 | 500 | 10 | 10 / 12 | 12 / 20 | 1.83 ± 0.03 |
| linear $R_{10}$ | 18 e | 7 | 500 | 10 | 10 / 11 | 11 / 20 | 1.98 ± 0.04 |
| $G_{10}$ | 10 e | 7 | 500 | 0 | 20 / 15 | 15 / 20 | 1.29 ± 0.03 |
| $G_{10}$ | 12 e | 7 | 500 | 0 | 20 / 14 | 14 / 20 | 1.47 ± 0.05 |
| $G_{10}$ | 14 e | 7 | 500 | 0 | 20 / 13 | 13 / 20 | 1.65 ± 0.04 |
| $G_{10}$ | 16 e | 7 | 500 | 0 | 20 / 12 | 12 / 20 | 1.81 ± 0.06 |
| $G_{10}$ | 18 e | 7 | 500 | 0 | 20 / 11 | 11 / 20 | 1.99 ± 0.09 |
| $R_{10}$ ILFF | 10 e | 7 | 500 | 10 | 10 / 15 | 15 / 20 | 1.20 ± 0.06 |
| $R_{10}$ ILFF | 12 e | 7 | 500 | 10 | 10 / 14 | 14 / 20 | 1.56 ± 0.03 |
| $R_{10}$ ILFF | 14 e | 7 | 500 | 10 | 10 / 13 | 13 / 20 | 1.67 ± 0.06 |
| $R_{10}$ ILFF | 16 e | 7 | 500 | 10 | 10 / 12 | 12 / 20 | 1.84 ± 0.06 |
| $R_{10}$ ILFF | 18 e | 7 | 500 | 10 | 10 / 11 | 11 / 20 | 2.02 ± 0.12 |

### Protein expression and purification

| Description | Assigned letter | Abbreviation |
|---|---|---|
| GBP1 | A | GBP1 A |
| TAMRA-GBP1 | B | GBP1 B |
| TAMRA-GBP1-BioRAM-$R_8$ | C | GBP1 C |
| TAMRA/BHQ-2-GBP1 | D | GBP1 D |
| TAMRA/BHQ-2-GBP1-BioRAM-$R_8$ | E | GBP1 E |
| NLS-mCherry | F | NLS-mCherry F |
| NLS-mCherry BioRAM-$R_4$ | G | NLS-mCherry G |
| FITC-GBP1 | H | NLS-mCherry H |
| FITC-GBP1-BioRAM-$R_8$ | I | NLS-mCherry I |

#### Sortase A 5M-H6(eSrtA)

Protein sequence:

MQAKPQIPKDKSKVAGYIEIPDADIKEPVYPGPATREQLNRGVSFAEENESLDDQNISIAGHTFIDRPNYQF
TNLKAAKKGSMVYFKVGNETRKYKMTSIRNVKPTAVEVLDEQKGKDKQLTLITCDDYNEETGVWETRKIF
VATEVKLEHHHHHH

For the expression, BL21 (DE3) cells were transformed with the pET29 plasmid (*addgene, #75144*). A single colony from an agar plate was picked and grown for 24 hours at 37 °C in 5 mL of LB medium with 40 µg/mL Kanamycin. The starting culture was then expanded to 1.0 L at 37 °C until $OD_{600}$ reached between 0.6-0.8, followed by induction using 0.4 mM IPTG and incubation at 30 °C for four hours under agitation.

Cells were collected by centrifugation at 5000× *g* for 15 min. The cells were washed once in PBS, then resuspended in PBS with 20 mM imidazole and 2 mM dithiothreitol (DTT) and lysed using sonication (3×2 min, 25 % Amplitude), followed by centrifugation at 25000× *g* for 30 min. For the purification, the clear lysate was loaded on PureCube 100 Ni-INDIGO agarose. The beads were washed with 20 column volumes (CV) of 10 mM Dulbecco's phosphate buffered saline (DPBS) with 20 mM imidazole. The protein was then eluted using 10 mM DPBS containing 500 mM imidazole. 5M SrtA-H6 was concentrated to desired stock concentration, liquid $N_2$ shock-frozen as aliquots, and stored at −70 °C.

intact Q-TOF HR-MS (ESI) mass calcd.: 17853 Da; found: 17854 Da.

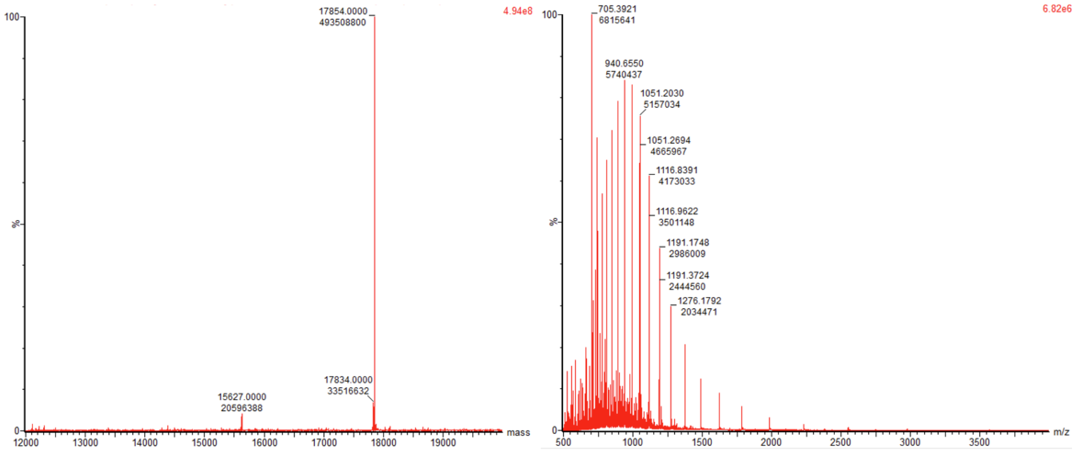

#### GBP1 (A)

Protein sequence (sequence after *Bovine Thrombin (Thr)* cleavage site is underlined):

MGSSHHHHHHSSGLVPRGSHDVQLVESGGALVQPGGSLRLSCAASGFPVNRYSMRWYRQAPGKERE
WVAGMSSAGDRSSYEDSVKGRFTISRDDARNTVYLQMNSLKPEDTAVYYCNVNVGFEYWGQGTQVTV
SS

For the expression Rosetta Gammi™ 2 (DE3) cells were transformed with a pET28a in-house generated plasmid. A single colony from an agar plate was picked and grown for 24 hours at 37 °C in 5 mL of TB medium with 40 µg/mL Kanamycin supplemented with 8 mM MgCl2. The starting culture was then expanded to 6 L (12 x 500 mL in 2 L glass flasks) at 37 °C, at 200 rpm until $OD_{600}$ reached 0.6-0.8, followed by induction using 0.5 mM IPTG and incubation at 18 °C overnight under agitation at 180 rpm.

Cells were collected by centrifugation at 5000 × *g* for 15 min. The cells were washed once in PBS, then resuspended in PBS with 5 mM imidazole and lysed using sonication (6×2 min, 25 % Amplitude), followed by centrifugation at 20000× *g* for 30 min. For the purification, the clear lysate was loaded on PureCube 100 Ni-INDIGO agarose. The beads were washed with 20 column volumes (CV) of 10 mM Dulbecco's phosphate buffered saline (DPBS) with 20 mM imidazole. The protein was then eluted using 10 mM DPBS containing 500 mM imidazole. The purification tag was removed by the addition of Thrombin (1:1000 (*v/v*)) overnight at 16 °C in dialysis against 10 mM DPBS followed size exclusion chromatography on Superdex 75 16/60 column. The GBP1 was concentrated to desired stock concentration, $LN_2$-shock-frozen as aliquots, and stored at −70 °C.

intact Q-TOF HR-MS (ESI) mass calcd.:12983 Da; found: 12983 Da.

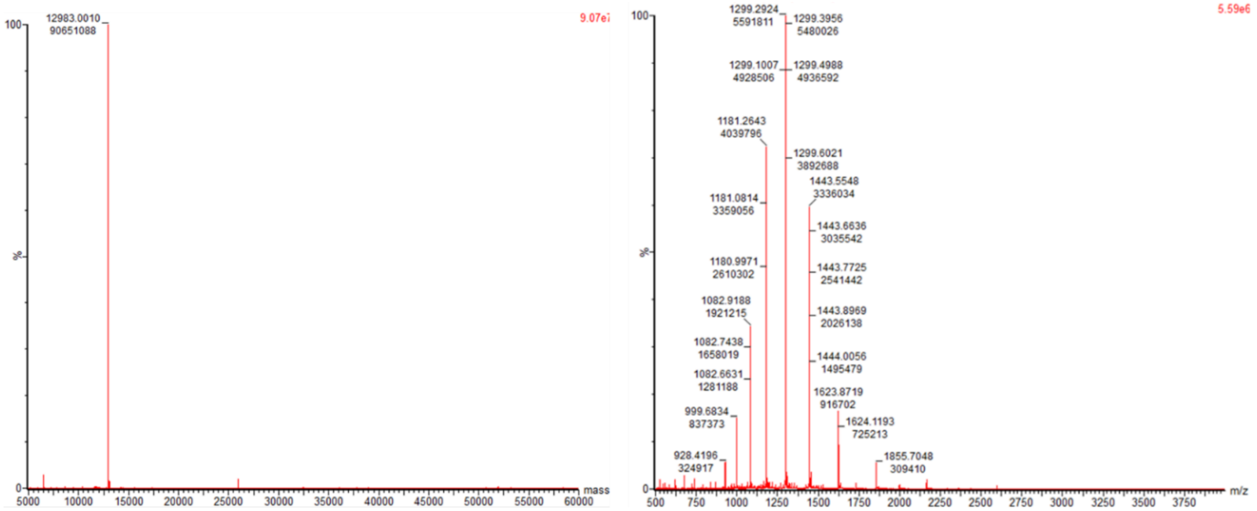

#### TAMRA-GBP1 (B)

General procedure for Sortase mediated ligation was followed using a final concentration of 50 µM GBP1 **B** (75 µL), 0.5 mM peptide **10** (3 µL) in 217.91 µL reaction buffer (300 µL total reaction volume).

intact Q-TOF HR-MS (ESI) mass calcd.:14128 Da; found: 14127 Da.

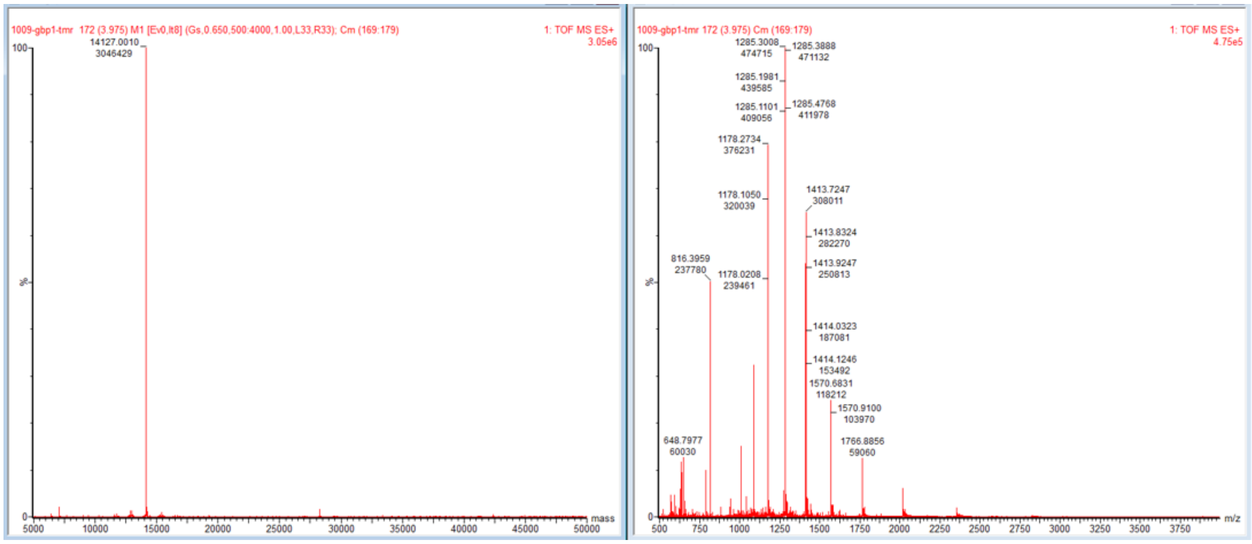

#### TAMRA-GBP1-BioRAM-$R_8$ (C)

General procedure for BioRAM modification was followed using a final concentration of 100 µM TAMRA-GBP1 **B** (14.83 µL), 2 mM peptide **13** (10 µL) in 25 µL reaction buffer (50 µL total reaction volume).

| **number of modifications** | **mass of modification (Da)** | **mass calculated (Da)** | **deconvoluted mass observed (m/z)** | **peak intensity (AU)** |
|---|---|---|---|---|
| **0** | | 14127 | 14127 | 0 |
| **1** | 1673.0 | 15800 | 15799 | 6192420 |
| **2** | 3346.0 | 17473 | 17472 | 18733116 |
| **3** | 5019.1 | 19146 | 19145 | 9619870 |
| **4** | 6692.1 | 20819 | 20818 | 1929402 |
| **5** | 8365.1 | 22492 | 0 | 0 |
| **6** | 10038.1 | 24165 | 0 | 0 |
| | | | average protein modification: | 2.20 |

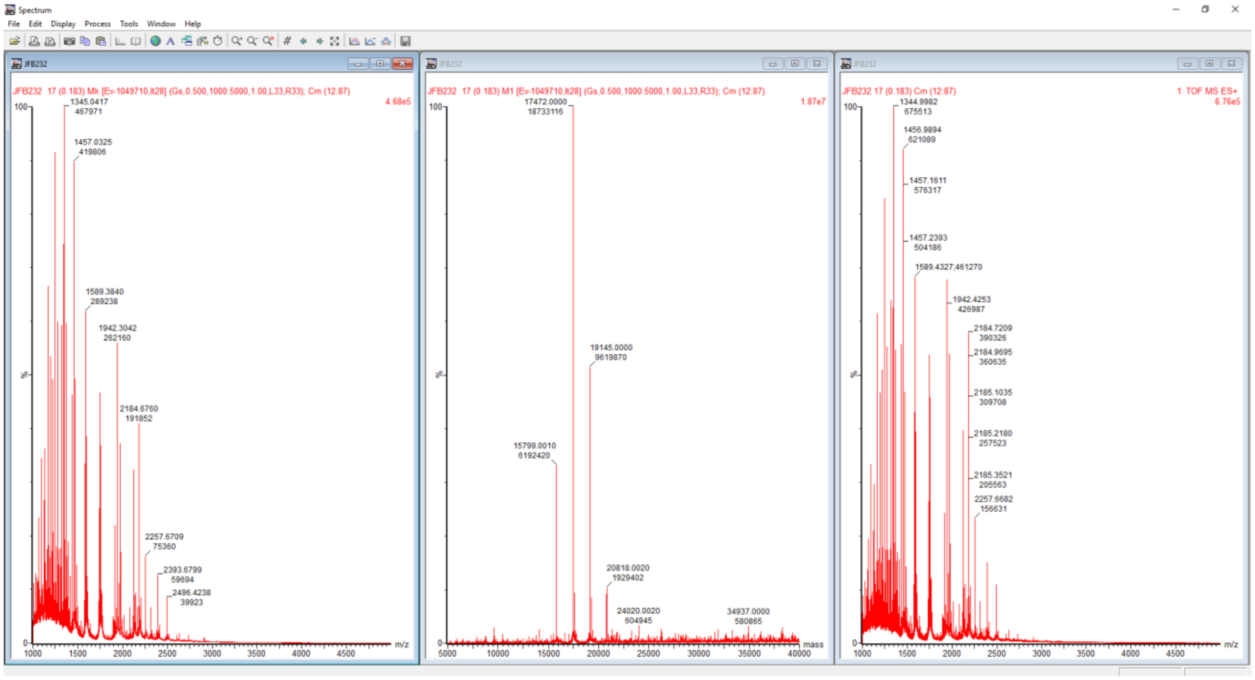
Spectrum
File Edit Display Process Tools Window Help
JFB232
JFB232 17 (0.183) Mk [Ev-1049710,It28] (Gs,0.500,1000:5000,1.00,L33,R33); Cm (12:87)
4.68e5
1345.0417
467971
1457.0325
419806
1589.3840
289238
1942.3042
262160
2184.6760
191852
2257.6709
75360
2393.6799
59694
2496.4238
39923
m/z
JFB232 17 (0.183) M1 [Ev-1049710,It28] (Gs,0.500,1000:5000,1.00,L33,R33); Cm (12:87)
1.87e7
17472.0000
18733116
19145.0000
9619870
15799.0010
6192420
20818.0020
1929402
24020.0020
604945
34937.0000
580865
mass
JFB232 17 (0.183) Cm (12:87)
1: TOF MS ES+
6.76e5
1344.9982
675513
1456.9894
621089
1457.1611
576317
1457.2393
504186
1589.4327;461270
1942.4253
426987
2184.7209
390326
2184.9695
360635
2185.1035
309708
2185.2180
257523
2185.3521
205563
2257.6682
156631
m/z

#### TAMRA/BHQ-2-GBP1 (D):

General procedure for Sortase mediated ligation was followed using a final concentration of 100 µM GBP1 **B** (150 µL), 1 mM peptide **12** (6 µL), 30 µM enzyme (8.2 µL) in 135.8 µL reaction buffer (300 µL total reaction volume).

intact Q-TOF HR-MS (ESI) mass calcd.: 15635 Da; found: 15634 Da.

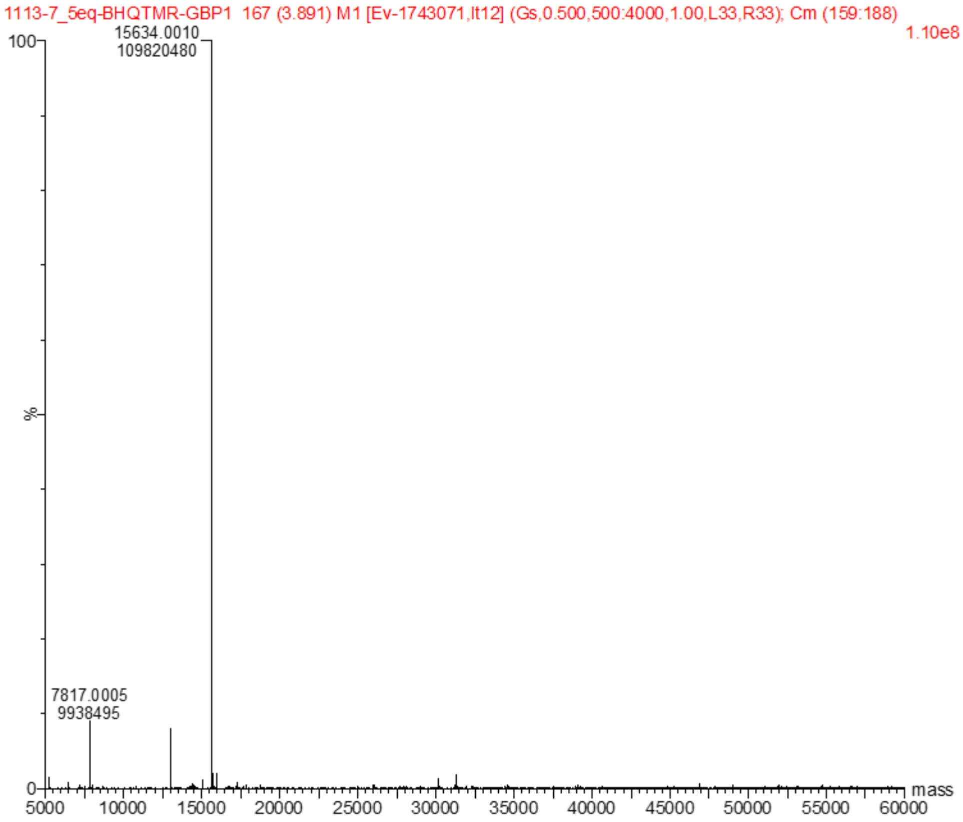

#### TAMRA/BHQ-2-GBP1-BioRAM-$R_8$ (E):

General procedure for BioRAM modification was followed using 100 µM TAMRA/BHQ-2-GBP1 **D** (44 µL), 1.85 mM peptide **13** (18.5 µL) in 100 µL reaction volume (37.5 µL reaction buffer).

| number of modifications | mass of modification | mass calculated | mass observed | peak intensity |
|---|---|---|---|---|
| **0** | | 15634 | - | 0 |
| **1** | 1673.0 | 17307 | - | 0 |
| **2** | 3346.0 | 18980 | 18990 | 4523082 |
| **3** | 5019.1 | 20653 | 20663 | 8992625 |
| **4** | 6692.1 | 22326 | 22336 | 3141151 |
| **5** | 8365.1 | 23999 | - | 0 |
| **6** | 10038.1 | 25672 | - | 0 |
| | | | average protein modification: | 2.92 |

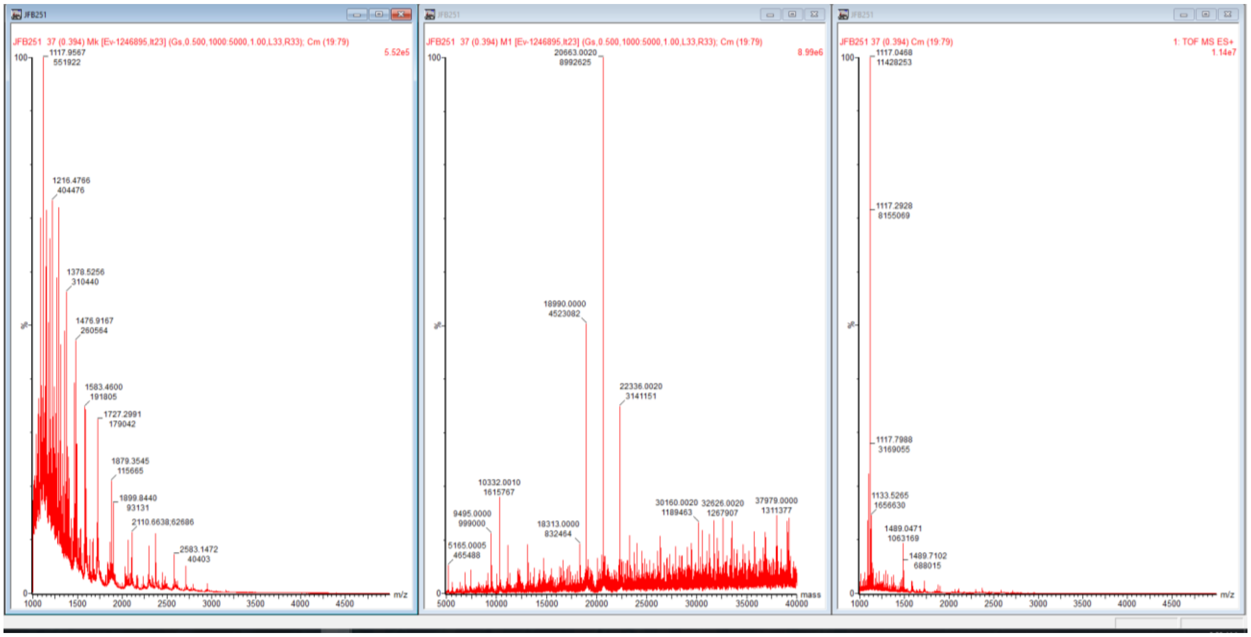

#### FITC-GBP1 (H)

General procedure for Sortase mediated ligation was followed using a final concentration of 50 µM GBP1 **B** (75 µL), 0.5 mM peptide **11** (3 µL),15 µM enzyme in 217.91 µL reaction buffer (300 µL total reaction volume).

intact Q-TOF HR-MS (ESI) mass calcd.:14075 Da; found: 14073 Da.

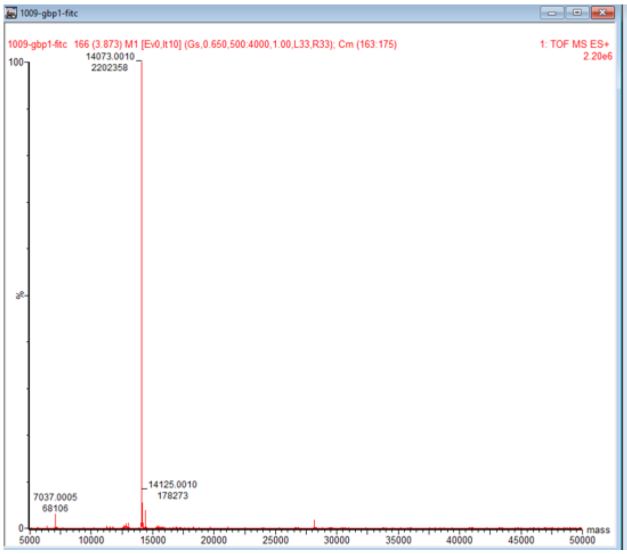
1009-gbp1-fitc
1009-gbp1-fitc 166 (3.873) M1 [Ev0,It10] (Gs,0.650,500:4000,1.00,L33,R33); Cm (163:175)
1: TOF MS ES+
2.20e6
14073.0010
2202358
14125.0010
178273
7037.0005
68106
100
%
0
5000
10000
15000
20000
25000
30000
35000
40000
45000
50000
mass

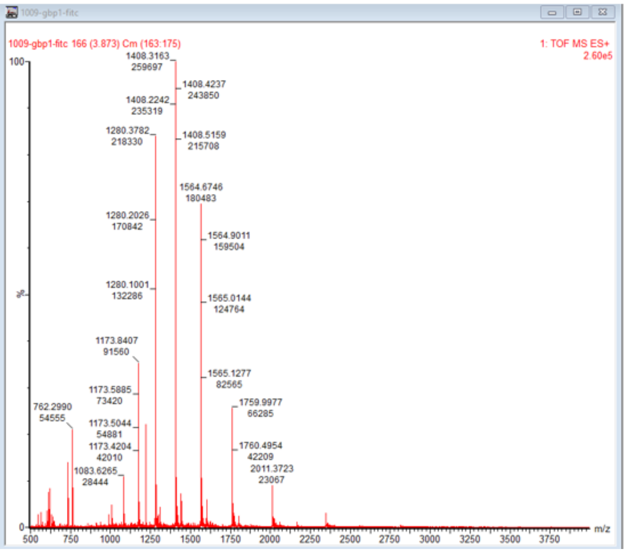
1009-gbp1-fitc
1009-gbp1-fitc 166 (3.873) Cm (163:175)
1: TOF MS ES+
2.60e5
1408.3163
259697
1408.4237
243850
1408.2242
235319
1280.3782
218330
1408.5159
215708
1564.6746
180483
1280.2026
170842
1564.9011
159504
1280.1001
132286
1565.0144
124764
1173.8407
91560
1565.1277
82565
1173.5885
73420
762.2990
54555
1759.9977
66285
1173.5044
54881
1173.4204
42010
1760.4954
42209
2011.3723
23067
1083.6265
28444
100
%
0
500
750
1000
1250
1500
1750
2000
2250
2500
2750
3000
3250
3500
3750
m/z

### FITC-GBP1-BioRAM-$R_8$ (I)

General procedure for BioRAM modification was followed using a final concentration of 100 µM FITC-GBP1 **H** (15 µL), 2 mM peptide **13** (10 µL) in 25 µL reaction buffer (50 µL total reaction volume).

| **number of modifications** | **mass of modification (Da)** | **mass calculated (Da)** | **deconvoluted mass observed (m/z)** | **peak intensity (AU)** |
|---|---|---|---|---|
| **0** | | 14073 | - | 0 |
| **1** | 1673.0 | 15746 | 15744 | 1654146 |
| **2** | 3346.0 | 17419 | 17417 | 6640157 |
| **3** | 5019.1 | 19092 | 19090 | 6370799 |
| **4** | 6692.1 | 20765 | 20763 | 1846969 |
| **5** | 8365.1 | 22438 | - | 0 |
| **6** | 10038.1 | 24111 | - | 0 |
| | | | average protein modification: | 2.51 |

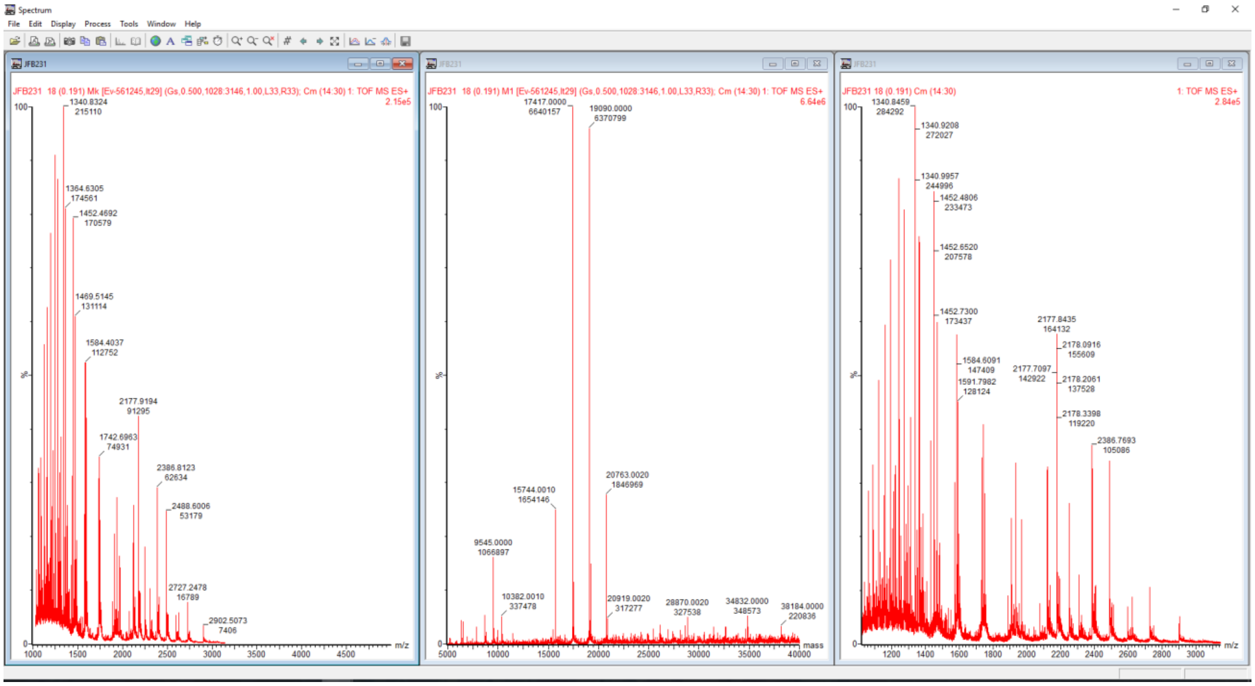

#### NLS-mCherry (F)

The protein was expressed as published previously.[22,23] Protein sequence (sequence after thrombin cleavage underlined, engineered cysteine in blue):

MGSSHHHHHHSSGLVPRGSHMPAAKRVKLDMVSKGEEDNMAIIKEFMRFKVHMEGSVNGHEFEIEG EGEGRPYEGTQTAKLKVTKGGPLPFAWDILSPQFMYGSKAYVKHPADIPDYLKLSFPEGFKWERVMN FEDGGVVTVTQDSSLQDGEFIYKVKLRGTNFPSDGPVMQKKTMGWEASSERMYPEDGALKGEIKQRL KLKDGGHYDAEVKTTYKAKKPVQLPGAYNVNIKLDITSHNEDYTIVEQYERAEGRHSTGGMDELYKACA*

Therefore, the thiol was alkylated prior to CPP to lysine conjugations using 10 equiv. iodoacetamide for 1 h at pH 7.4. The protein was desalted using Zeba™ Spin (ThermoFisher) filtration (7k MWCO).

QTof HR-MS analysis: calcd.: 28394; found: 28395.

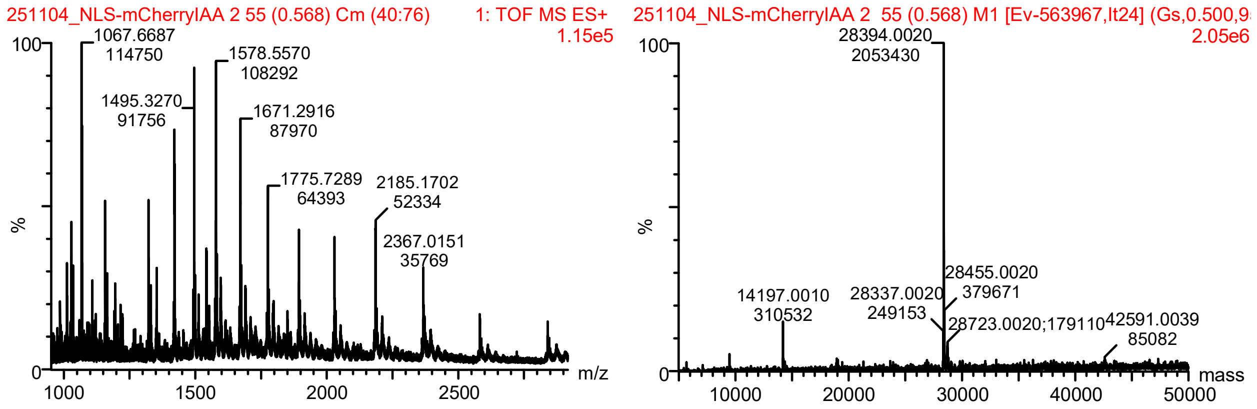

#### NLS-mCherry-BioRAM-R$_4$ (G)

General procedure for BioRAM modification was followed using a final concentration of 200 µM NLS-mCherry **F** (425 µL), 1.2 mM peptide **14** (103 µL) in 322 µL reaction buffer (850 µL total reaction volume).

| number of modifications | mass of modification (Da) | mass calculated (Da) | deconvoluted mass observed (m/z) | peak intensity (AU) |
|---|---|---|---|---|
| **0** | | 28395 | - | 0 |
| **1** | 1048.3 | 29443 | 29444 | 468081 |
| **2** | 2096.5 | 30492 | 30491 | 3117282 |
| **3** | 3144.8 | 31540 | 31539 | 6887892 |
| **4** | 4193.0 | 32588 | 32587 | 9696214 |
| **5** | 5241.3 | 33636 | 33636 | 10086924 |
| **6** | 6289.6 | 34685 | 34684 | 6881597 |
| **7** | 7337.8 | 35733 | 35732 | 2964004 |
| **8** | 8386.1 | 36781 | 36780 | 768933 |
| **9** | 9434.3 | 37829 | - | 0 |
| | | | average protein modification: | 4.52 |

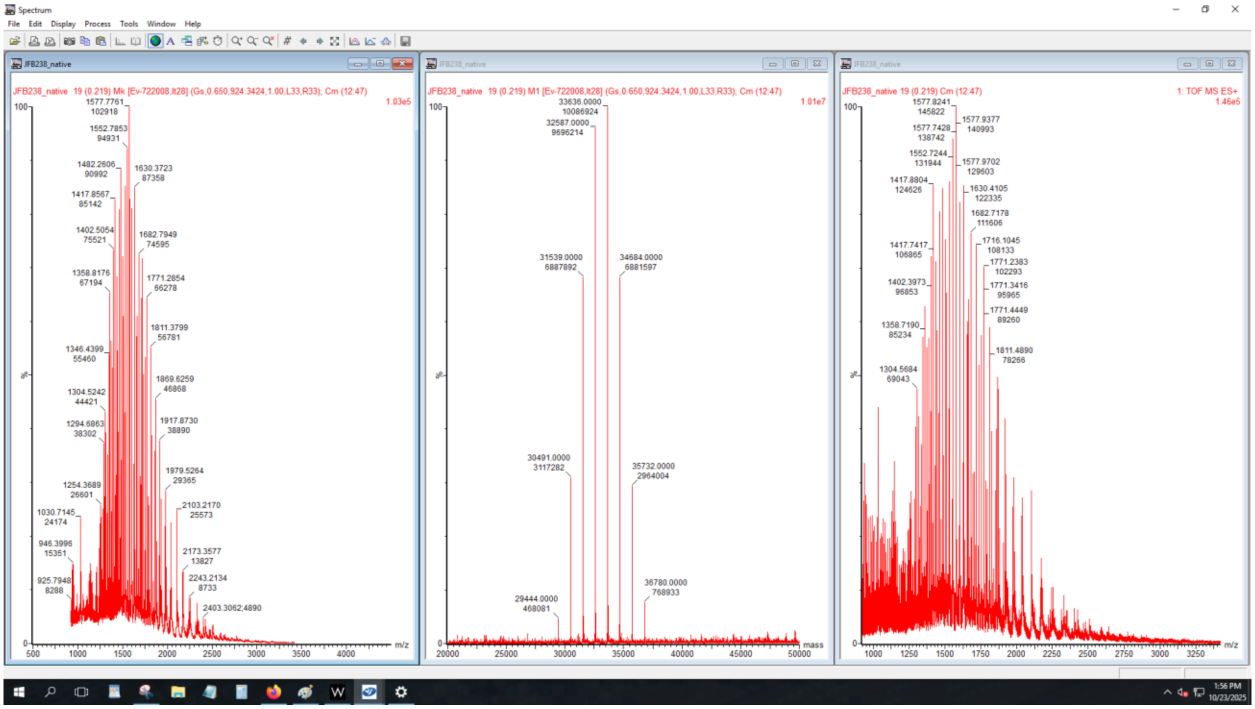
Spectrum
File Edit Display Process Tools Window Help
JFB238_native
JFB238_native 19 (0.219) Mk [Ev-722008,It28] (Gs,0.650,924:3424,1.00,L33,R33); Cm (12:47)
1.03e5
1577.7761 102918
1552.7853 94931
1482.2606 90992
1630.3723 87358
1417.8567 85142
1402.5054 75521
1682.7949 74595
1358.8176 67194
1771.2854 66278
1811.3799 56781
1346.4399 55460
1869.6259 46868
1304.5242 44421
1294.6863 38302
1917.8730 38890
1979.5264 29365
1254.3689 26601
1030.7145 24174
2103.2170 25573
946.3996 15351
2173.3577 13827
925.7948 8288
2243.2134 8733
2403.3062;4890
m/z
JFB238_native 19 (0.219) M1 [Ev-722008,It28] (Gs,0.650,924:3424,1.00,L33,R33); Cm (12:47)
1.01e7
33636.0000 10086924
32587.0000 9696214
31539.0000 6887892
34684.0000 6881597
30491.0000 3117282
35732.0000 2964004
29444.0000 468081
36780.0000 768933
mass
JFB238_native 19 (0.219) Cm (12:47)
1: TOF MS ES+
1.46e5
1577.8241 145822
1577.9377 140993
1577.7428 138742
1552.7244 131944
1577.9702 129603
1417.8804 124626
1630.4105 122335
1682.7178 111606
1716.1045 108133
1417.7417 106865
1771.2383 102293
1402.3973 96853
1771.3416 95965
1771.4449 89260
1358.7190 85234
1811.4890 78266
1304.5684 69043
m/z
1:56 PM
10/23/2025

### Microscopy Macros

**Time-lapse microscopy to estimate co-localization of CPP-additive and fCPP signal**

```
// FIJI macro for batch quantification of TAMRA signal in defined ROIs
// Images are imported as hyperstacks via Bio-Formats and processed frame by frame.

// setBatchMode(true);
// Optional: can be uncommented to run the macro without continuously displaying images
// (recommended for large datasets to improve performance).

dir  = getDirectory("Choose Source Directory ");

dir2 = getDirectory("Choose Output Directory ");

list = getFileList(dir);

// Loop over all files in the source directory
for (i = 0; i < list.length; i++) {

    file = list[i];
    run("Bio-Formats Importer",
        "open=[" + dir + "" + file + "] " +
        "autoscale " +
        "color_mode=Default " +
        "concatenate_series " +
        "open_all_series " +
        "rois_import=[ROI manager] " +
        "view=Hyperstack " +
        "stack_order=XYCZT");

    // Retrieve image dimensions, including the number of time frames.
    getDimensions(width, height, channels, slices, frames);

    // Loop over all time frames of the current hyperstack
    for (f = 1; f <= frames; f++) {

        Stack.setFrame(f);
        name = getTitle();
        nameseries = name + "_" + f;
        // Segmentation of the iRFP channel (channel 3) to define ROIs
        // Channel 3 is assumed to contain the iRFP signal used to define nucleation
        // and inverse regions of interest.

        run("Duplicate...", "duplicate channels=3 frames=" + f);
        run("Top Hat...", "radius=6 stack");
        // Applies a morphological top-hat filter with a radius of 6 pixels to enhance
        // local bright structures on a slower varying background across the stack.

        setAutoThreshold("Triangle dark stack no-reset");
        // Automatically determines a global threshold using the Triangle method,
        // assuming objects are darker than the background, and applies it over the stack.

        run("Create Mask");
```

```
    // Converts the thresholded image into a binary mask (foreground vs. background).

    run("Dilate");
    // Expands the foreground regions in the binary mask by one pixel, reducing
    // potential fragmentation and tightening the segmented objects.

    run("Create Selection");

    // Quality control: skip frames where no ROI could be defined
    if (selectionType == -1) {
        selectWindow(name);
        close("\\Others");
        continue;
    }

    roiManager("Add");
    // Adds the current selection (iRFP-positive region) to the ROI Manager.
    // This ROI is interpreted as the "nucleation" region of interest.

    run("Make Inverse");
    // Inverts the current selection, generating the complementary ROI that
    // corresponds to all pixels outside the nucleation region within the image.

    if (selectionType == -1) {
        // If the inverse selection fails (no valid complementary region),
        // the frame is skipped.
        selectWindow(name);
        close("\\Others");
        continue;
    }

    roiManager("Add");
    // Adds the inverse selection as a second ROI in the ROI Manager.
    // This ROI corresponds to the non-nucleation (background or surrounding) region.

    // Quantification of RFP signal (channel 2) inside the defined ROIs
    selectWindow(name);
    Stack.setFrame(f);
    Stack.setChannel(2);
    // Selects channel 2, assumed here to represent the RFP signal

    roiManager("measure");
    roiManager("Delete");

    // Cleanup for the current frame
    selectWindow(name);
    close("\\Others");
    // Closes all non-active windows created during processing
    // (e.g., duplicated channel images, masks) while keeping the main
    // hyperstack open for the next iteration.
}
// Export measurements for the current file and reset the Results table
saveAs("Results", dir2 + name + "_Results.csv");
// Saves all accumulated measurements for the current hyperstack as a CSV file
```

```
    // in the output directory, with a file name derived from the image title.

    run("Clear Results");
}
```

**Time-lapse experiments using DIBAC$_4$(3) to measure change of membrane potential upon CPP-treatment**

```
setBatchMode(true);
// Enables batch mode, which suppresses image display and accelerates processing
// when large numbers of images are analyzed sequentially.

// User input: define input (raw data) and output (results) directories
dir  = getDirectory("Choose Source Directory ");
// Prompts the user to select the directory containing the raw microscopy files

out  = getDirectory("Choose Destination for Results ");
// Prompts the user to select the directory in which the quantitative results
// tables (.csv) will be stored.

list = getFileList(dir);
// Retrieves an array containing the names of all files

// Loop over all files in the source folder
for (i = 0; i < list.length; i++) {

   file = list[i];
   // Restrict processing to specific microscopy formats
   if (endsWith(file, ".tif") || endsWith(file, ".lif") || endsWith(file, ".nd2")) {

      // Import image as a multi-dimensional hyperstack using Bio-Formats
      run("Bio-Formats Importer",
         "open=[" + dir + file + "] autoscale color_mode=Default " +
         "concatenate_series open_all_series rois_import=[ROI manager] " +
         "view=Hyperstack stack_order=XYCZT");
      // Opens the current file with Bio-Formats, preserving channel, z-slice,
      // and time information and representing the data as an XYCZT hyperstack.

      getDimensions(width, height, channels, slices, frames);
      // Retrieve image dimensions, including the number of time frames.

      // Analysis restricted to channel 1
      Stack.setChannel(1);
      // fluorescence channel of the DIBAC probe (green/GFP) to be quantified

      imageName = getTitle();
      // Stores the window title (derived from the file name) for use in
      // naming output files.

      // Loop through all time frames in the selected channel
      for (f = 1; f <= frames; f++) {

         Stack.setFrame(f);
         // Activates the current time frame in channel 1.

         name = getTitle();
         nameseries = name + "_" + f;
         // Frame-specific identifier
```

```
        // Duplicate the current frame for image processing
        run("Duplicate...", "title=Proc_Dup_Frame_" + f + " frame=" + f);
        selectWindow("Proc_Dup_Frame_" + f);
        // Brings the duplicated frame window to the foreground for processing.

        // Preprocessing and segmentation by intensity thresholding
        run("Gaussian Blur...", "sigma=10");
        // Applies a Gaussian blur with sigma = 10 pixels to smooth intensity
        // facilitating robust global thresholding.

        setAutoThreshold("MinError dark");
        // Computes an automatic intensity threshold using the \"Minimum Error\"
        // method, assuming that objects of interest are darker than the
        // background, and applies this threshold to the current image.

        run("Create Selection");
        roiManager("Reset");
        // Clears any pre-existing ROIs from the ROI Manager to avoid mixing

        roiManager("Add");
        // Adds the current selection from the processed duplicate image to
        // the ROI Manager as the ROI to be quantified in this frame.

        run("Measure");
        // Quantitative measurement of the selected ROI
        close();
        // Closes the duplicated processed frame window to free memory.
      }

      // Export the accumulated measurements for the current image
      saveAs("Results", out + name + "_Results_green.csv");
      // Saves all rows currently present in the Results table as a CSV file

      run("Clear Results");
      // Clears the Results table so that measurements from the next input file

      run("Close All");
      // Closes all open image windows (including the original hyperstack)

      roiManager("Reset");
      // Clears all ROIs from the ROI Manager
    }
  }
```

**Endsome substraction and quantification of nuclear signal to quantify cytosolic delivery of protein**

```
// setBatchMode(true);
// Optional: suppresses image display and speeds up batch processing.

// Select directory containing multi-channel image stacks
dir = getDirectory("Choose Source Directory ");
list = getFileList(dir);

// Loop over all files in the selected folder
for (i = 0; i < list.length; i++) {
  file = list[i];

  // Import file as XYCZT hyperstack via Bio-Formats
  run("Bio-Formats Importer", "open=[" + dir + "" + file + "] autoscale color_mode=Default "
    + "concatenate_series open_all_series rois_import=[ROI manager] view=Hyperstack
stack_order=XYCZT");
  getDimensions(width, height, channels, slices, frames);

  // Each file is a time series (frames) with two channels (Hoechst, RFP)
  for (f = 1; f <= frames; f++) {
    Stack.setFrame(f);
    name = getTitle;
    nameseries = name + "_" + f;

    // Hoechst channel (1): segment nuclei and exclude edge-touching objects
    run("Duplicate...", "duplicate channels=1 frames=f");
    run("Median...", "radius=5");                    // Noise reduction
    setAutoThreshold("Huang dark no-reset");         // Automatic global threshold
    run("Convert to Mask");                          // Binary mask of nuclei
    run("Fill Holes");                               // Solid nuclear regions
    run("Analyze Particles...",
      "size=20-Infinity circularity=0.20-1.00 show=Outlines exclude add");
    // Detect nuclei as ROIs; exclude objects touching image borders.

    // RFP channel (2): define non-endosomal region by manual threshold
    selectWindow(name);
    run("Duplicate...", "duplicate channels=2 frames=f");

    // Threshold should be adjusted per experiment (based on maximal uptake condition).
    setThreshold(6000, 65535, "raw");
    run("Create Selection");
    run("Make Inverse");
    roiManager("Add");                               // Add non-endosomal ROI
    // Identify index of non-endosomal ROI (last entry in ROI Manager)
    roiCount     = roiManager("count");
    nonEndosomeROI = roiCount - 1;
    roiManager("Select", nonEndosomeROI);
    roiManager("Rename", "nonEndosomeROI");

    // Intersect each nuclear ROI with the non-endosomal ROI
    // and reorganize ROI Manager accordingly
    for (j = 0; j < nonEndosomeROI; j++) {
      // Select current first ROI and the non-endosomal ROI
```

```
            roiManager("Select", newArray(0, nonEndosomeROI - j));
            roiManager("AND");// Geometric intersection

            // Skip empty intersections (no overlap)
            type = selectionType();
            if (type == -1) {
                print("null");
            } else {
                roiManager("add"); // Store intersected ROI
            }

            // Remove the original first ROI that has now been processed
            roiManager("Deselect");
            roiManager("Select", 0);
            roiManager("Delete");
        }

        // After all nuclei have been processed, remove the non-endosomal ROI
        roiManager("Select", 0);
        roiManager("Delete");

        // Measure intensity in all newly generated ROIs
        run("Select All");
        roiManager("Show All");
        roiManager("Measure");

        // Create (if needed) and define output directory for this experiment
        dir2 = dir + "/filename/";
        File.makeDirectory(dir2);

        // Optionally save ROIs instead of deleting them
        // roiManager("Deselect");
        // roiManager("save", dir2 + name + ".zip");

        // Reset ROI Manager and close auxiliary windows before next frame
        roiManager("Delete");
        selectWindow(name);
        close("\\Others");
    }

    // Export per-image measurements and clear Results table
    saveAs("Results", dir2 + name + "_Results.csv");
    run("Clear Results");
}
```

**Endosome versus nuclear signal to estimate direct translocation over NLS-mCherry over time**

```
// Batch macro for endosome/nucleus segmentation and quantification

dir = getDirectory("Choose Source Directory ");
list = getFileList(dir);

for (i = 0; i < list.length; i++) {
   file = list[i];

   // Import as XYCZT hyperstack
   run("Bio-Formats Importer", "open=[" + dir + "" + file + "] autoscale color_mode=Default "
      + "concatenate_series open_all_series rois_import=[ROI manager] view=Hyperstack
stack_order=XYCZT");
   getDimensions(width, height, channels, slices, frames);

   for (f = 1; f <= frames; f++) {
      Stack.setFrame(f);
      name = getTitle();
      nameseries = name + "_" + f;

      // Hoechst nuclei segmentation
      run("Duplicate...", "duplicate channels=1 frames=f");
      run("Median...", "radius=5");                      // Noise reduction
      setAutoThreshold("Huang dark no-reset");           // Global threshold
      run("Convert to Mask");
      run("Fill Holes");
      run("Create Selection");
      roiManager("Add");                                 // Add nuclei ROI

      // Non-endosomal region (RFP channel 2, threshold >15000)
      selectWindow(name);
      run("Duplicate...", "duplicate channels=2 frames=f");
      setThreshold(15000, 65535, "raw");                 // Endosomal signal cutoff
      run("Create Selection");
      run("Make Inverse");
      roiManager("Add");                                 // Add non-endosomal ROI

      // Intersect nuclei with non-endosomal region
      selectWindow(name);
      roiManager("Select All");
      roiManager("AND");                                 // Nuclei ∩ non-endosomes
      roiManager("add");                                 // Store intersected ROI
      run("Select All");
      roiManager("Measure");                             // Measure intersected regions

      roiManager("Delete");

      // Whole cell segmentation (RFP total signal)
      selectWindow(name);
      run("Duplicate...", "duplicate channels=2 frames=f");
      run("Gaussian Blur...", "sigma=20 stack");         // Smooth for cell detection
      setAutoThreshold("MinError dark");
      run("Create Selection");
```

```
        roiManager("add");                              // Add cell ROI

        // Measure total RFP signal in cells
        selectWindow(name);
        run("Duplicate...", "duplicate channels=2 frames=f");
        run("Measure");                                 // Whole cell intensity

        // Cleanup
        selectWindow(name);
        close("\\Others");
        roiManager("Delete");
    }

    // Save results
    dir2 = dir + "/endosomesVSnucleus/";
    File.makeDirectory(dir2);
    saveAs("Results", dir2 + name + "_Results.csv");
    run("Clear Results");
}
```

### Peptide Synthesis

#### Peptide Synthesis and modifications

Peptides were synthesized by Fluorenylmethoxycarbonyl (Fmoc)-solid-phase peptide synthesis (SPPS) using a peptide synthesizer (Automated microwave peptide synthesizer, CEM) on TentaGel S Ram resin (RappPolymere, Tübingen, Germany, 0.05–0.1 mmol scale, 0.22 mmol/g). Couplings were achieved by reacting 0.2 M Fmoc-AA-OH with 0.25 M *N*,*N'*-Diisopropylcarbodiimide (DIC) and 0.25 M ethyl cyanoacetate (Oxyma) in dimethylformamide (DMF). A solution of 20% Piperidine in DMF was used to remove the Fmoc protection group. The acetylation is carried out with 3.5 ml DMF, 1 ml acetic anhydride and 0.5 ml diisopropylamine (DIPEA) overnight at room temperature. Arginine was incorporated with 2,2,4,6,7-pentamethyldihydro-benzofuran-5-sulfony (Pbf) protection using double couplings, cysteine was incorporated on the *N*-termini with Trityl-, 3-nitro-2-pyridinesulfenyl- (NPYS) or tert-butyl- ($S^tBu$) protection, in case activation of the disulfide was performed on resin.

#### Peptide final deprotection and cleavage, purification, and characterization

Peptides were deprotected and cleaved from the resin using a mixture of 10 mL trifluoroacetic acid (TFA), 0.75 g phenol, 0.5 mL water, 0.5 mL methylphenylsulfide, and 0.25 mL 1,2-ethandithiol (EDT). After at room temperature, the cleavage solution was collected, and the crude peptides were precipitated from ice-cold $^t$Butyl-methyl ether. Crude peptides were washed five times with dry diethyl ether. Final deprotection was done by adding 95:2.5:2.5 of TFA/trifluoroacetic acid (TIS)/$H_2O$ overnight at 25 ºC. Peptides were purified by Preparative reverse phase-high performance liquid chromatography (RP-HPLC) on a Shimadzu system using a VP 250/21Nucleodur C18 Htec column (Macherey-Nagel, 100 A, 5 µm, 250 mm x 21 mm, 10 mL/min). The following gradient was used in all purifications: A = $H_2O$ + 0.1 % TFA, B = acetonitrile (MeCN) + 0.1% TFA 5 % B 0–5 min, 5–95 % B 5-60 min, 95 % 63-70 min. UPLC-UV traces were obtained on a Waters H-class instrument equipped with a Quaternary Solvent Manager, a Waters autosampler and a Waters TUV detector with an Acquity UPLC-BEH C18 1.7 µm, 2.1× 50 mm RP column. The Empower 3 software (Waters) was used. The following gradient was used: A = $H_2O$ + 0.1 % TFA, B = MeCN + 0.1 % TFA 5-95 % B 0-5 min, flow rate 0.6 mL/min. UPLC-UV chromatograms were recorded at 220 nm. Masses were measured using a Xevo G2-XS QTof (Waters) high-resolution mass spectrometer coupled to an Acquity UPLC system running on water and acetonitrile, both with 0.01% formic acid using the MassLynx software (V4.1, Waters).

| Entry | Compound Nr. | Peptide | Sequence | remark | reference |
|---|---|---|---|---|---|
| 1 | **1** | TNB-$R_{10}$-ILFF (CPP-additive) | $NH_2$-C(TNB)-Peg-Peg-RRRRRRRRRR-ILFF-$CONH_2$ | a | Arafiles et al. [24] |
| 2 | **2** | TNB-Cy5-$R_{10}$-ILFF (Cy5-CPP-additive) | $NH_2$-C(TNB)-Peg-K(Cy5)-Peg-RRRRRRRRRR-ILFF-$CONH_2$ | a | Arafiles et al. [2424] |
| 3 | **3** | fCPP | TAMRA-G-C(BHQ-2)-Peg-RRRRRRRRRR-$CONH_2$ | b | - |
| 4 | **4** | qCPP | TAMRA-G-C(BHQ-2)-Peg-RRRRRRRRRR-$CONH_2$ | c | - |
| 5 | **10** | TAMRA-sortase peptide | TAMRA-Peg-Peg-LPETGG-$CONH_2$ | b | - |
| 6 | **11** | FITC-sortase peptide | TAMRA-Peg-Peg-LPETGG-$CONH_2$ | b | - |
| 7 | **12** | TAMRA/BHQ-2-sortase peptide | TAMRA-ER-C(BHQ-2)-ERER-Peg-LPETG-$CONH_2$ | c | - |
| 8 | **13** | BioRAM-$R_4$ | ac-C(2-Mp-4-Np-carbonate)-Peg-RRRR-$CONH_2$ | - | Franke et al. [2] |

| | | | | | |
|---|---|---|---|---|---|
| 9 | **14** | BioRAM-$R_8$ | ac-C(2-Mp-4-Np-carbonate)-Peg-RRRRRRRR-$CONH_2$ | - | Franke et al[2] |

Peg = 8-amino-3,6-dioxaoctanoic acid.
(2-Mp-4-Np-carbonate) = 2-mercaptopropyl (4-nitrophenyl) carbonate (*via disulfide)*
TAMRA= 5-Carboxytetramethylrohodamine
ac = acetylated *N*-terminus

a)

The activation of $NH_2$-C-Peg-Peg-$R_{10}$-ILFF-$CONH_2$ to $NH_2$-C(TNB)-Peg-Peg-$R_{10}$-ILFF-$CONH_2$ was performed on resin before cleavage. Therefore Fmoc-Cys($S^tBu$)-COOH building block was used during microwave synthesis. After Fmoc-deprotection, the $S^tBu$ protecting group was removed using 20 % β-mercaptoethanol in DMF at room temperature overnight. After thorough washing, cysteine activation was performed using Ellman´s reagent (5,5-dithio-bis-(2-nitrobenzoic acid)) in 25 % ethanol in DMF. Cleavage was performed using only TFA/TIS/$H_2O$ (95:2.5:2.5).

b)

To the resin-bound peptide, 3 eq. of the respective fluorophore carboxylic acid, 3 eq. of *O*-(7-Azabenzotriazol-1-yl)-*N*,*N*,*N′*,*N′*-tetramethyluronium-hexafluorphosphat (HATU), and 10 eq. of DIPEA were added in DMF (0.1 M in coupling reagent). The resin was incubated with the reaction mixture for 2 hours at room temperature. After washing with 3×DMF and 3×DCM, cleavage was conducted following the general protocol (**3**: 31% yield*, **10**: 56% yield*, **11**: 42% yield*; *calculated from resin loading).

c)

Peptide sequence was synthesized on the solid phase. TAMRA acid was conjugated on resin to the free *N*-terminus using 3 eq. of HATU, 10 eq. of DIPEA with 3 eq. of the respective acid in DMF for 2 hours at room temperature. After washing with 3×DMF and 3×DCM, cleavage was conducted following the general protocol. Peptide was purified using RP-HPLC as described in the general methods. Lyophilized peptide (**3**: 10.74 mg; for **12**: 7.2 mg) was then incubated with 1.1 eq. of disulfide activated BHQ-2 **9** and 6 eq. DIPEA in 200 µL DMF for 2 h. Peptides were precipitated using ice-cold diethyl ether (-20 °C). After centrifugation the supernatant was discarded and the precipitate was air-dried for 10 minutes. The pallet was dissolved in ACN/water (1:10), filtered and purified by RP-HPLC as described in the general methods. Peptide was obtained as dark purple powder (**4**: 66%; **12**: 33% from purified precursor peptide)

**3** – fCPP:

**Q-TOF HR-MS (ESI)** *m/z* calcd. for $C_{96}H_{168}N_{46}O_{19}S^{6+}$: 383.7219 [*M*+6H]$^{6+}$; found: 383.7254 [*M*+H]$^{6+}$.

**UPLC-MS (ESI):**

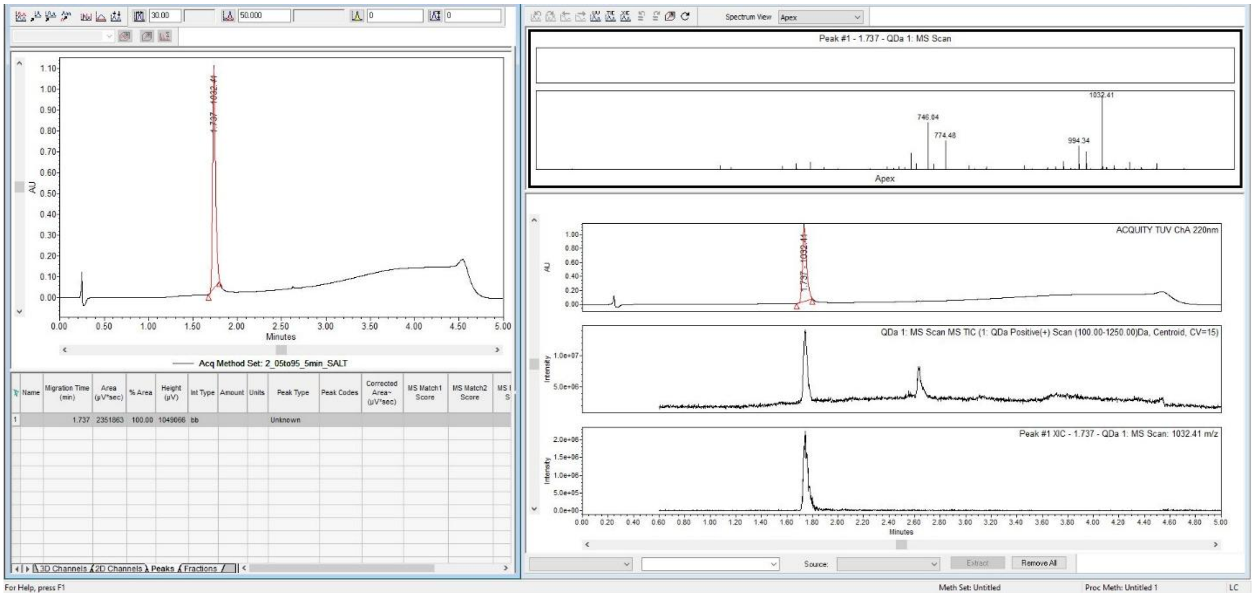

**4** – qCPP:

**Q-TOF HR-MS (ESI)** *m/z* calcd. for $C_{122}H_{195}N_{53}O_{24}S_2^{6+}$: 475.2518 [*M*+6H]$^{6+}$; found: 475.2509 [*M*+H]$^{6+}$.

**UPLC-MS (ESI):**

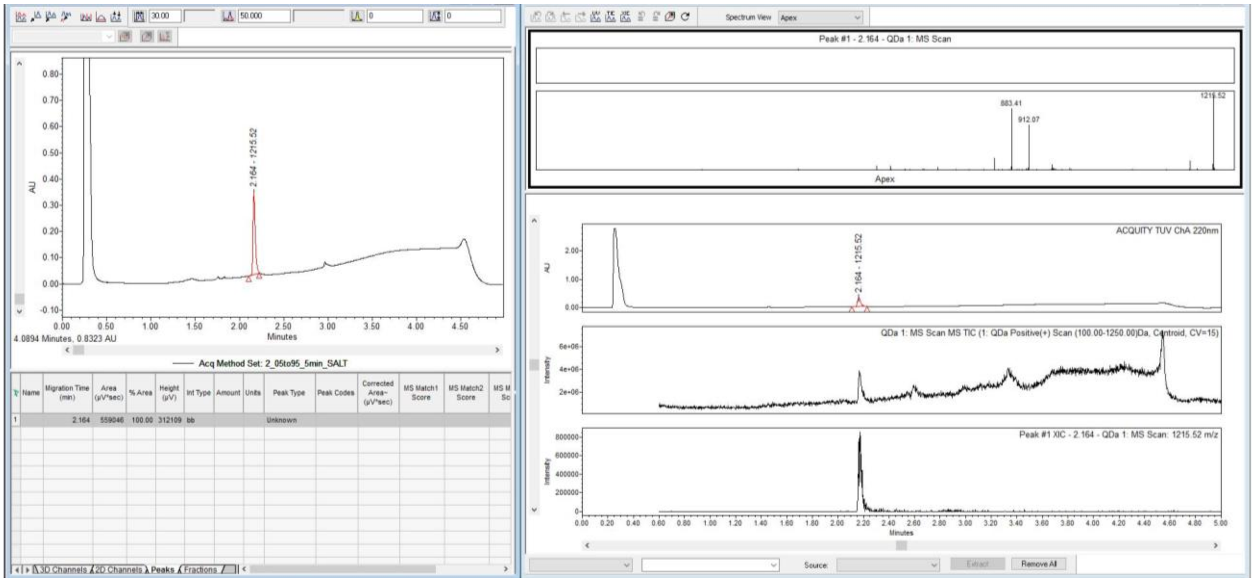

**10** – TAMRA-sortase peptide:

**Q-TOF HR-MS (ESI)** *m/z* calcd. for $C_{61}H_{83}N_{11}O_{19}$: 638.19 [*M*+2H]$^{2+}$; found: 638.21 [*M*+H]$^{2+}$.

**UPLC-MS (ESI):**

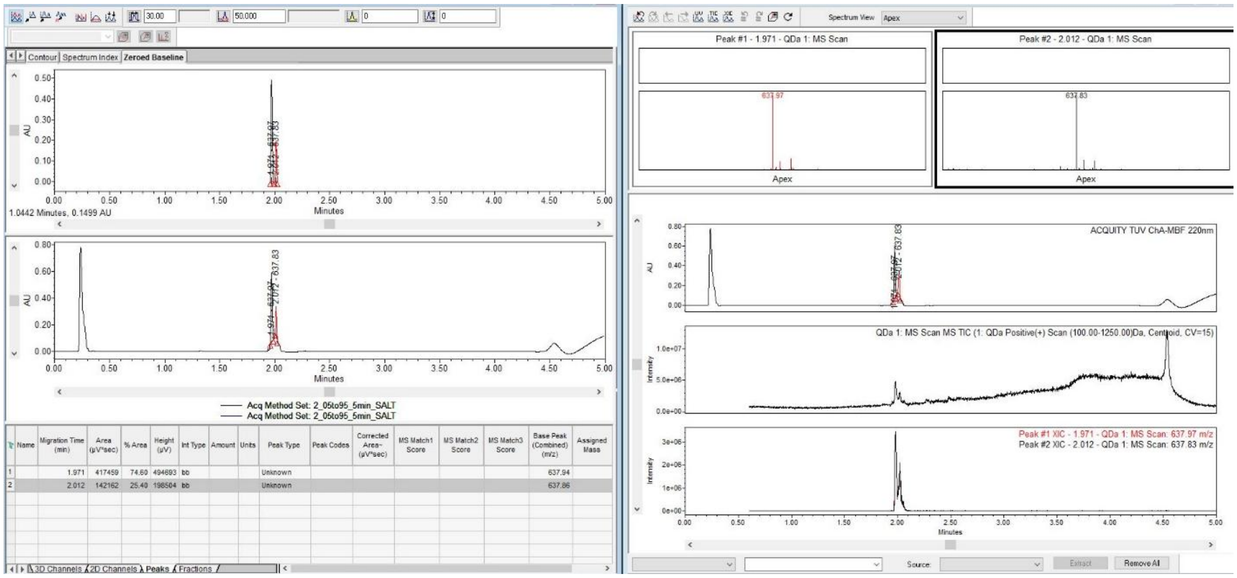

*the two peaks correspond to the respective TAMRA isomers.*

**11** – FITC-sortase peptide

**Q-TOF HR-MS (ESI)** *m/z* calcd. for $C_{57}H_{74}N_{10}O_{20}S^{2+}$: 610.12 [*M*+H]$^{2+}$; found: 610.74 [*M*+H]$^{2+}$.

**UPLC-MS (ESI):**

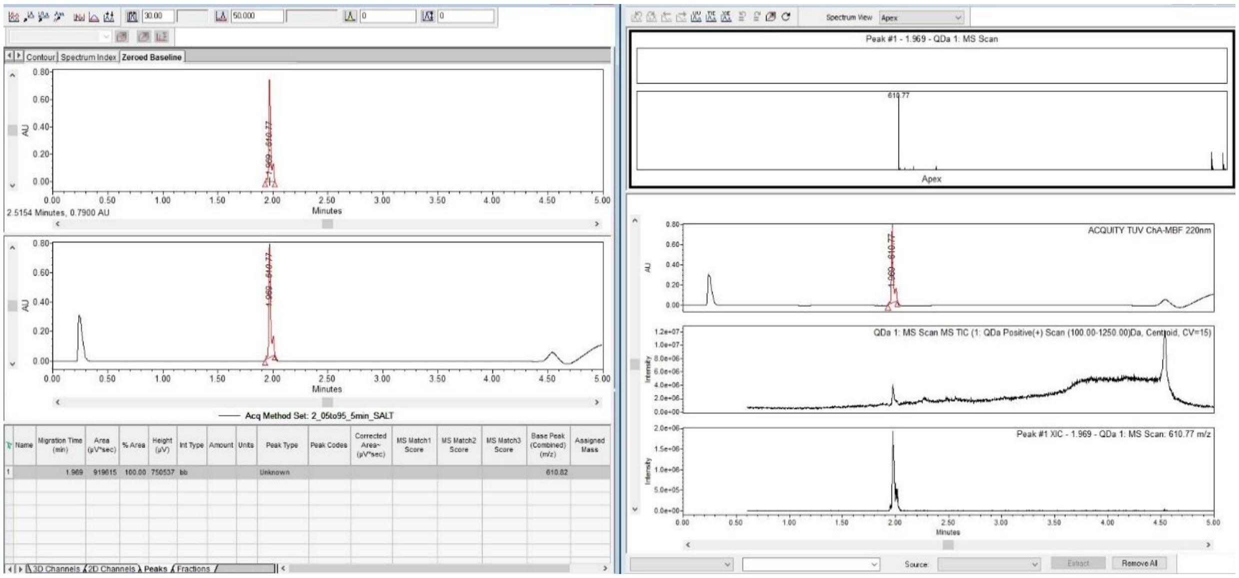

**12** – TAMRA/BHQ-2-sortase peptide:

**Q-TOF HR-MS (ESI)** *m/z* calcd. for $C_{96}H_{168}N_{46}O_{19}S^{6+}$: 383.7219 [*M*+6H]$^{6+}$; found: 383.7254 [*M*+H]$^{6+}$.

**UPLC-MS (ESI):**

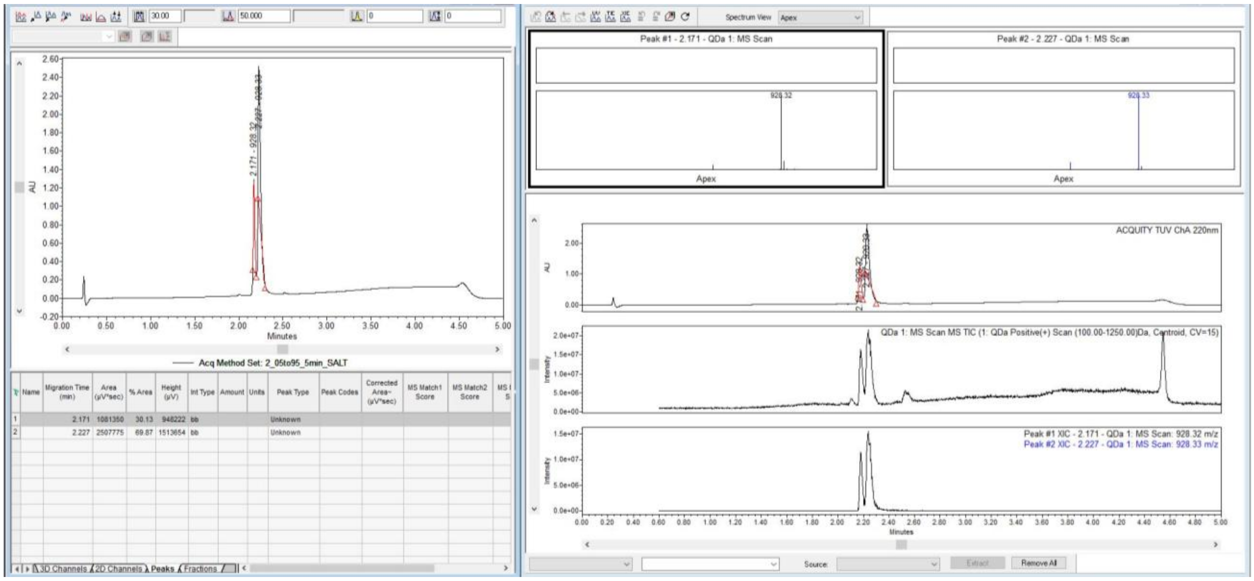

**the two peaks correspond to the respective TAMRA isomers.*

### Organic Synthesis

##### 6 - 3-(methyl(phenyl)amino)propanoic acid

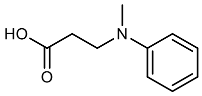

**6**

Synthesis was performed according to the literature.[25] In short: To a solution of N-methylaniline (2.2 mL, 2.14 g, 20 mmol, 2.0 eq.) in EtOH (30 mL) were added 3-bromopropionic acid (1.53 g, 10 mmol, 1.0 eq.) and AcONa (1.64 g, 20 mmol, 2.0 eq.). The reaction mixture was stirred for 6 h at reflux temperature followed by 16h at room temperature. After the addition of water, the mixture basified using NaOH (aq.) and washed five times with diethylether (5x30 mL). The aqueous layer was acidified with HCl aq. to pH ~ 5 and extracted with diethylether (30 mL). The organic layer was evaporated and the residue was purified by column chromatography on silica gel (eluent: $CH_2Cl_2$/MeOH = 98/2→97/3) to afford 2 (1.18 g, 6.58 mmol) as a white solid in 66% yield.

**$^1$H-NMR** (600 MHz, $CDCl_3$): δ [ppm] = 7.21–7.19 (m, 2H), 6.79–6.69 (m, 3H), 3.66 (t, 2H, J = 7.1 Hz), 2.93 (s, 3H), 2.53 (t, 2H, J = 7.0 Hz).

##### 7 - 4-((4-((E)-(2,5-dimethoxy-4-((E)-(4-nitrophenyl)diazenyl)phenyl)diazenyl)phenyl)(methyl)amino)butanoic acid

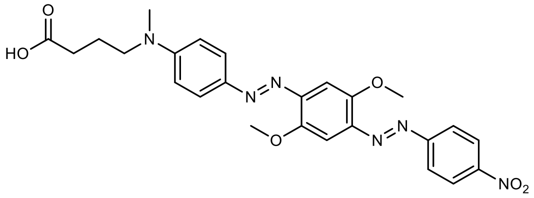

**7**

Compound 3 was synthesized following the general procedure for bioconjugatable BHQ-2s.[26] Fast Black K Salt hemi zinc chlorid (1.2 eq., 6.70 mmol, 2.34 g) was dissolved in a mixture of NaOAc buffer (0.1 M, pH 4.0) and MeCN (1:1, v/v, 20 mL). The mixture was sonicated for 3 min and stirred for 10 to reach almost complete solubilization. In another round bottom flask, the 3-(methyl(phenyl)amino)propanoic acid (1 eq., 5.58 mmol, 1.0 g) was dissolved in 5 mL MeCN and cooled to 0 °C using an ice bath. The solubilized black salt was filtered and added to the aniline in solution. The reaction mixture was stirred for 2 hours at room temperature. The precipitate was centrifuged and washed twice with water in MeCN (1:1, v/v, 2x20 mL). The residue was lyophilized overnight resulting in a deep purple-black powder (>95% purity by LC-MS) and was not purified further.

**Q-TOF HR-MS (ESI)** *m/z* calcd. for $C_{25}H_{27}N_6O_6^+$: 493.1830 [*M*+H]$^+$; found: 493.1804 [*M*+H]$^+$.

**UPLC-MS (ESI)** *m/z* calcd. for $C_{25}H_{27}N_6O_6^+$: 493.18 [*M*+H]$^+$; found: 492.98 [*M*+H]$^+$:

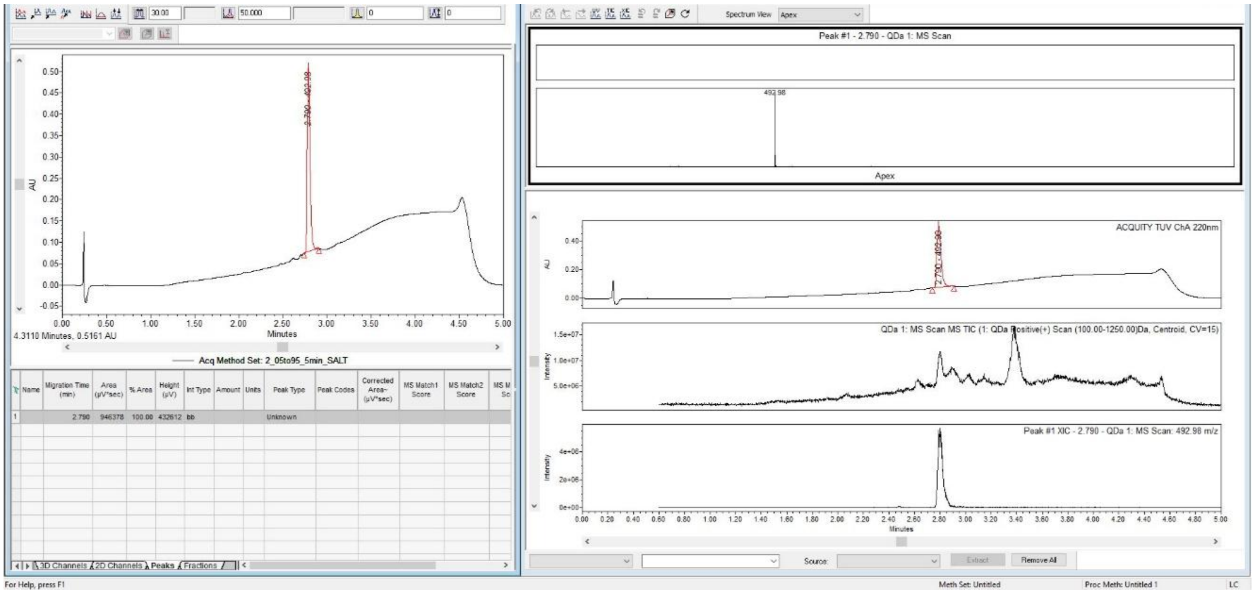
Spectrum View Apex
Peak #1 - 2.790 - QDa 1: MS Scan
492.98
Apex
AU
Minutes
4.3110 Minutes, 0.5161 AU
Acq Method Set: 2_05to95_5min_SALT
Name
Migration Time (min)
Area (µV*sec)
% Area
Height (µV)
Int Type
Amount
Units
Peak Type
Peak Codes
Corrected Area~ (µV*sec)
MS Match1 Score
MS Match2 Score
1 2.790 946378 100.00 432612 bb Unknown
3D Channels
2D Channels
Peaks
Fractions
ACQUITY TUV ChA 220nm
QDa 1: MS Scan MS TIC (1: QDa Positive(+) Scan (100.00-1250.00)Da, Centroid, CV=15)
Intensity
Peak #1 XIC - 2.790 - QDa 1: MS Scan: 492.98 m/z
Source:
Extract
Remove All
For Help, press F1
Meth Set: Untitled
Proc Meth: Untitled 1
LC

**8 - 2-((5-nitropyridin-2-yl)disulfaneyl)ethan-1-amine**

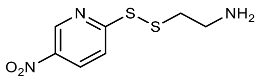

**8**

Compound 8 was synthesized following the procedure of Genentech (WO2017/201449). 1,2-bis(5-nitropyridin-2-yl)disulfane (1.2 eq, 8.17 mmol, 2.54 g) and 2-aminoethane-1-thiol (1.0 eq, 6.81 mmol, 0.525 g) were dissolved in 10 mL DCM:MeOH (8:2, *v/v*). Acetic acid (2 eq, 13.62 mmol, 0.82 g) was added and the reaction mixture was stirred for 72 hours. The solvent was evaporated under reduced pressure and the residue was resuspended in 100 % DCM. The mixture was centrifuged and the supernatant discarded. After washing twice using 2x20 mL of DCM the precipitate was obtained as pale-yellow solid (93% purity by LC-MS).

**UPLC-MS (ESI)** *m/z* calcd. for $C_{25}H_{27}N_6O_6^+$: 232.02 [*M*+H]$^+$; found: 231.8 [*M*+H]$^+$:

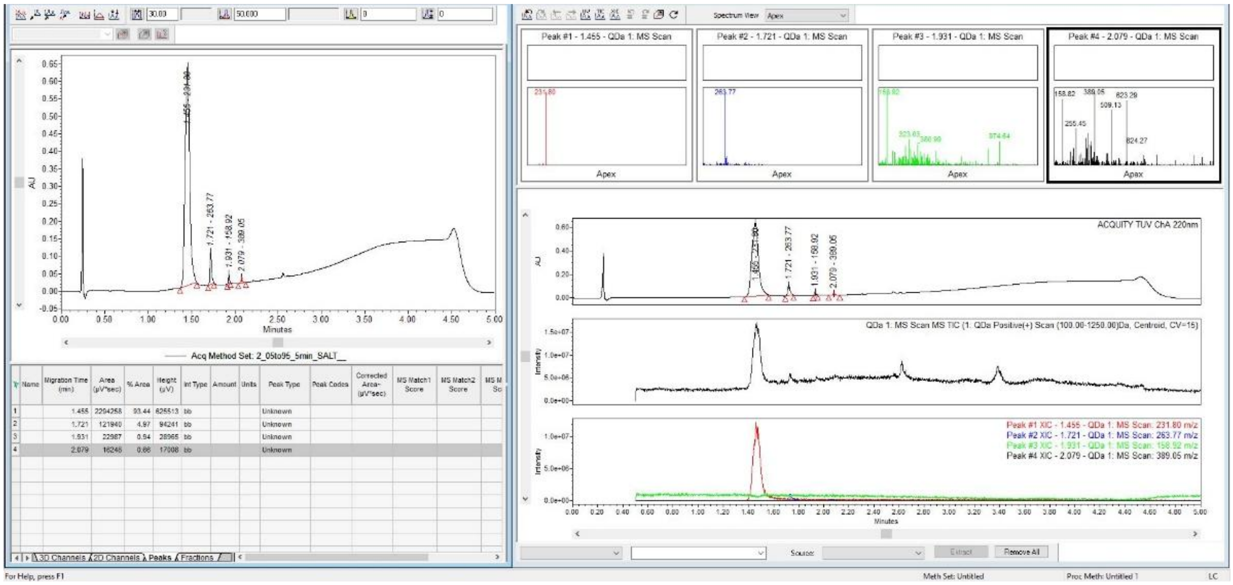

**9 - 3-((4-((E)-(2,5-dimethoxy-4-((E)-(4-nitrophenyl)diazenyl)phenyl)diazenyl)phenyl)(methyl)amino)-N-(2-((5-nitropyridin-2-yl)disulfaneyl)ethyl)propanamide**

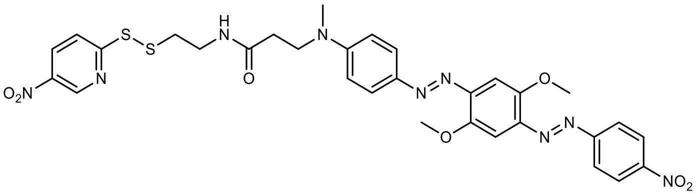

**9**

A 50 mL round bottom flask was charged with BHQ-2 acid 3 (1 eq., 118 µmol, 58 mg), HOBt (1.5 eq, 177 µmol, 27 mg), DIPEA (5 eq., 589 µmol, 76 mg, 103 µL) and 10 mL DCM. The reaction mixture was cooled to 0 °C followed by the addition of EDC (1.5 eq., 177 µmol, 34 mg). Compound 4 (1.5 eq., 177 µmol, 41 mg) was dissolved in 4 mL DMF and added to the reaction mixture. The reaction was stirred at 0 °C for 1 hour, allowed to warm up to room temperature and stirred for another 5 hours. The solvent was evaporated under reduced pressure and the residue was taken up in EtOAc and washed with 0.1 M NaOH twice. The organic solvent was evaporated and purified by Flash-Chromatography (eluent: $CH_2Cl_2$/MeOH = 99/1→95/5). The product eluted as dark purple band. Evaporation of the solvent afforded a black solid (~80% purity by LC-MS) which was used for conjugation to peptides without further purification. Subsequent purification using RP-HPLC yielded the product with a purity exceeding 95% (LC-MS, see spectrum attached).

**UPLC-MS (ESI)** *m/z* calcd. for $C_{31}H_{32}N_9O_7S_2^+$: 706.19 [*M*+H]$^+$; found: 706.18 [*M*+H]$^+$.

**Q-TOF HR-MS (ESI)** *m/z* calcd. for $C_{31}H_{32}N_9O_7S_2^+$: 706.1861 [*M*+H]$^+$; found: 706.1826 [*M*+H]$^+$.

**UPLC-MS (ESI):**

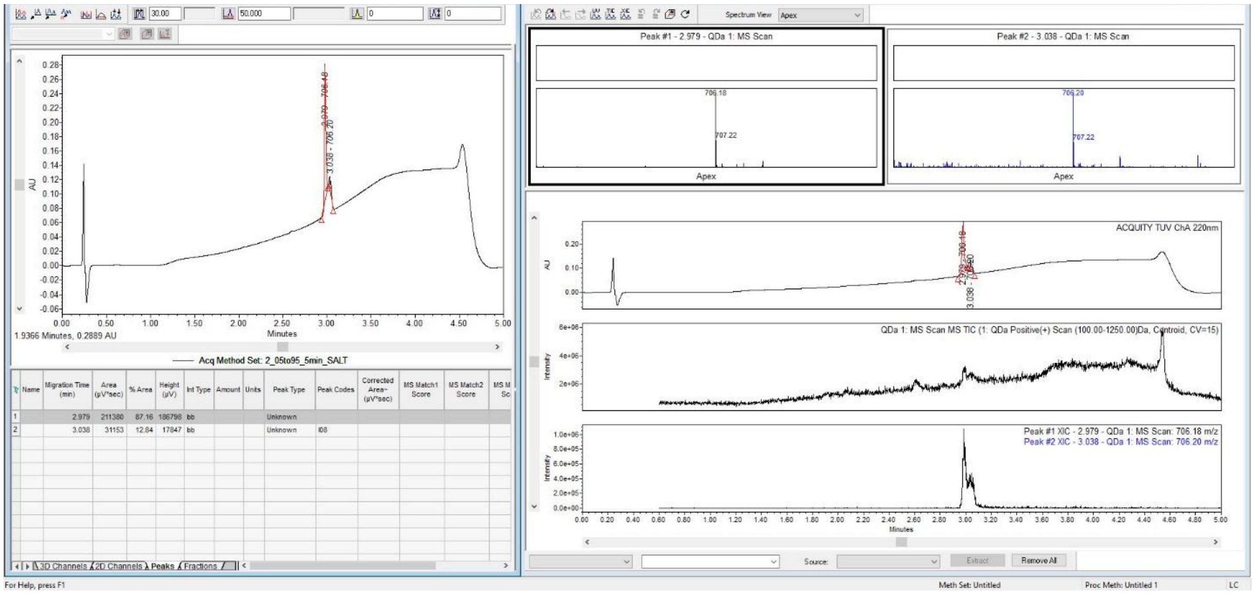

**2 peaks of the same mass might be a result of the azo-isomers*
